# Hitchhiking of bactericidal/permeability-increasing protein-like gene with the fibromelanosis locus in Kadaknath black-bone-chicken

**DOI:** 10.1101/2022.07.28.501823

**Authors:** Sagar Sharad Shinde, Ashutosh Sharma, Nagarjun Vijay

## Abstract

Black-bone-chicken (BBC) meat is popular for its distinctive taste and texture. A complex chromosomal rearrangement at the fibromelanosis (*Fm*) locus on the 20th chromosome results in increased expression of the endothelin-3 (*EDN3*) gene and is responsible for melanin hyperpigmentation in BBC. We use public long-read sequencing data of the silkie breed to resolve high-confidence haplotypes at the *Fm* locus spanning both Dup1 and Dup2 regions and establish that the *Fm_2* scenario is correct of the three possible scenarios of the complex chromosomal rearrangement. The relationship between Chinese and Korean BBC breeds with Kadaknath native to India is underexplored. Our data from whole-genome re-sequencing establishes that all BBC breeds, including Kadaknath, share the complex chromosomal rearrangement junctions at the fibromelanosis (*Fm*) locus. We also identify two *Fm* locus proximal regions (∼70Kb and ∼300 Kb) with signatures of selection unique to Kadaknath. These regions harbor several genes with protein-coding changes, with the Bactericidal/permeability-increasing-protein-like (*BPIL*) gene having two Kadaknath-specific changes within protein domains. Our results indicate that protein-coding changes in the *BPIL* gene hitchhiked with the *Fm* locus in Kadaknath due to close physical linkage. Identifying this *Fm* locus proximal selective sweep sheds light on the genetic distinctiveness of Kadaknath compared to other BBC.

## Introduction

Domestication of chicken during the Neolithic period involved a complex pattern of interbreeding with various jungle fowl species[1–4]. After domestication, chicken has spread worldwide and occurs as commercial, exotic, and indigenous village breeds. Humans use chicken as a research model due to their physiology and behaviour[5], as game fowl and for religious reasons, or more commonly for egg or meat production[6]. Chicken is the most preferred source of meat for humans due to its easy availability and affordability[7]. Hence, understanding the genetics of meat features is commercially relevant[8]. Although understudied, indigenous village chicken breeds with unique properties provide an opportunity to understand the genetics of meat. For instance, the black-bone chicken (BBC) is a delicacy due to its texture, color, firmness, flavor, and use in traditional medicine[9, 10]. The black color of BBC results from melanin deposition throughout the body, i.e., melanin hyperpigmentation or fibromelanosis caused by the *Fm* allele[11]. Bateson and Punnet were the pioneers in identifying the autosomal dominant *Fm* allele[12]. Modern studies found a chromosomal rearrangement on chromosome 20 is involved in the *Fm* locus[13, 14]. The overexpression of the endothelin-3 (*EDN3*) gene located within the *Fm* locus is responsible for hyperpigmentation seen in BBC[13,15,16].

BBC breeds have anti-fatigue and anti-hypoxic abilities, with its meat having antioxidant properties[9, 17], high carnosine[18, 19], and lower fat and cholesterol content[10, 20]. Some BBC breeds also have local adaptations. For instance, the Korean Ogye has improved fetal viability and innate immunity against microbial and viral infections[21]. BBC breeds occur globally and have distinct names, such as Ayam Cemani (Indonesia), Black H’Mong (Vietnam), Tuzo (Argentina), Svarthöna (Sweden)[14, 22], Yeonsan Ogye (Korea)[23], and Thai BBC (Thailand)[24]. China has a high diversity of BBC breeds, including Silkie, Jiangshan, Lueyang, Sichuan, Xingwen, Yugan, Dehua, Jinhu, Muchuan, Wumeng, Yanjin, Xichuan, Tuanfu, Wuliangshan, Emei and Miyi fowl[9,25–28]. It is unclear whether all BBC breeds share the same rearrangement at the *Fm* locus, as independent structural variants can produce similar phenotypes[29]. India has a single breed of BBC, commonly known as Kadaknath[19].

Kadaknath is considered native to the Jhabua, Alirajpur, and Dhar districts of Madhya Pradesh, but its farming has recently spread across India [30, 31]. The Kadaknath breed has long been documented as a distinctive indigenous Indian breed and is also called Kali Masi or Karaknath [32]. At least three distinctive phenotypes (i.e., Jetblack, Pencil, and Golden) occur within the Kadaknath breed[33]. In Jetblack, all the body parts like plumage, comb, internal organs, eyes, skin, beak, shank, and claw are entirely black, whereas Pencil and Golden have white and golden color patches on the plumage, respectively[33]. Several unique characteristics, such as earlier egg-laying maturity, high protein content, better disease resistance, and adaptation to the local environment, are attributed to Kadaknath chicken [19,30,33–41]. Understanding the genetics behind these traits will help establish the uniqueness of Kadaknath and guide de-extinction efforts and breeding programs.

Despite its immense popularity and commercial value, the genomics of the Kadaknath chicken breed has received limited attention. Therefore, further research and genomic analyses are required to understand adaptations in this breed and its genetic history. The aims of this study are : (1) Perform whole-genome re-sequencing of Kadaknath and evaluate its relationship with other black-bone and non-black-bone chicken breeds. (2) Assess whether Kadaknath and other BBC breeds share a common origin for the *Fm* locus by comparing the chromosomal rearrangement junction. (3) Evaluate how the BBC breeds dispersed to various parts of Southeast Asia. (4) Use population genetic statistics to identify signatures of selection in Kadaknath compared to other BBC breeds. Our results offer an example of the co-selection of traits regulating immune response, meat quality, and plumage color providing new approaches to improve breed quality and productivity in chicken simultaneously.

## Materials and Methods

### Population sampling

The study was approved by the Institutional Ethics Committee (IEC) of the Indian Institute of Science Education and Research, Bhopal, vide reference number IISERB/IEC/Certificate/2018-11/03 dated 8^th^ June 2018. We purchased the meat of six individuals (two individuals each from Jetblack, Pencil, and Golden morphs (**Fig. S1**)) of Kadaknath from an FSSAI (Food Safety and Standards Authority of India) licensed shop in Bhopal, Madhya Pradesh, India. We procured two other individuals with a black-bone phenotype from the same FSSAI-licensed shop to examine hybrids. One had a Golden-Pencil external appearance, and another was completely white on the exterior. We also sampled two non-black-bone chicken from the same shop to determine the genetic relationship of Kadaknath with native-village chicken and broiler reared in the same poultry (see **Fig. S1**). To reduce the chances of inter-breeding and have relatively pure Kadaknath samples, we obtained three (one individual each from Jetblack, Pencil, and Golden morphs) additional Kadaknath breed chickens from Jhabua, Madhya Pradesh. Whole-genomic DNA with high purity and quality was extracted from the tissue samples using DNeasy Blood & Tissue Kits (QIAGEN). We generated >25x coverage whole-genome short-read paired-end data (using Illumina Novaseq) for all 13 individuals sampled (see **Table S1**).

To evaluate the relationship of Kadaknath chicken globally, we compared our dataset with publicly available chicken re-sequencing data. High-quality re-sequencing datasets of 88 chicken individuals from other BBC, commercial chicken lines, and other distinctive chicken breeds with a coverage >20x were selected and obtained from ENA (Europen Nucleotide Archive) and (KNABIC) Korean National Agricultural Biotechnology Information Center (https://nabic.rda.go.kr/). Out of 88 individuals, 22 are BBC, which we considered for further analysis. Hence, we analyze a dataset of 101 individuals (88 public + 13 sequenced as part of this study) from different breeds (see **Table S2** for more details)[21,23,27,42–47].

### Read mapping, variant calling, and phylogeny

We mapped the paired-end raw reads of 101 individuals to the chicken genome assembly (genome assembly version is Gallus_gallus.GRCg6a) using the BWA[48] (Burrows-Wheeler Aligner) v0.7.17-r1188 read mapper with default parameters. We added the read group information using Picard tools and removed duplicate reads in all 101 individual bam files (https://github.com/broadinstitute/picard). We performed the variant calling using FreeBayes[49]with different quality control flags such as --min-alternate-count -C 10, --min-mapping-quality -m 20, --min-base-quality -q 20, and --min-coverage 10. Similarly, we used the bcftools[50],[51] with mapping quality flags such as --mapping quality -C 50, --min base quality -Q 20, and --min mapping quality -q 20 for robust variant calling. We removed the indels from variant calls using vcftools[52] with the -remove-indels flag and extracted the common SNPs using both SNP callers for retaining reliable SNP calls. The Single Nucleotide Polymorphisms (SNPs) identified by both variant callers (bcftools v1.9 and FreeBayes v1.0.0) were used for subsequent analysis. To identify the effect of genetic variants, we used the snpEff[53]. Common SNPs from both vcf files were used to construct phylogeny for 34 and 101 individuals using SNPhylo[54] v20180901, based on a maximum-likelihood tree with 1000 bootstrap values. We excluded the scaffolds, Z, W, and MT chromosomes in phylogeny analysis. The local phylogeny for Dup1 and Dup2 regions was generated using the vk phylo command (using both NJ and UPGMA methods) implemented in VCF-kit[55]. The Mt genome haplotype median-joining networks were constructed using PopART[56] and SplitsTree[57].

### *Fm* locus junction identification

The *Fm* locus consists of a complex chromosomal rearrangement composed of two different non-paralogous regions (Dup1 (∼127 Kb) and Dup2 (∼170 Kb)) separated by an intermediate (Int) region. Dup1 and Dup2 regions are both duplicated and are involved in a complex rearrangement consisting of two junctions (**A**) Dup1 + (inverted Dup2) and (**B**) (inverted Dup1) + Dup2. To identify the base-pair level positions of Dup1, Dup2, and Int regions, we compared the short-read coverage of black and non-black chicken in 1Kb windows along chromosome 20. We used the makewindows command of bedtools[58] (v2.26.0) to create 1Kb non-overlapping windows along chromosome 20. The number of reads in each 1Kb window was calculated using the bedtools coverage command. We shortlisted adjacent windows with drastically different read coverage in black individuals but not in non-black individuals. The base-pair level coordinates of Dup1 and Dup2 in the Gallus_gallus.GRCg6a genome was narrowed down further using coverage estimates in 1bp windows. Dup1 starts at 20:10766772 and ends at 20:10894151. Dup2 occurs further along the chromosome and starts at 20:11306686 and ends at 20:11477501. The read coverage at these junctions consistently differed between all black-bone and non-black-bone chicken. In addition to the Dup1, Dup2, and Int regions, we defined ∼500 Kb flanking regions as Flank1 (20:10263555-10766771) and Flank2 (20:11477502-11980000).

### Black-bone-specific *Fm* locus junction

The junction between the rearranged regions in the BBC does not occur in the Gallus_gallus.GRCg6a genome (**Fig. S2, 3**). Hence, we searched for the *Fm* locus junction in the Korean BBC PacBio data from ENA (SRR6189090). Based on our search of PacBio reads that mapped to the Gallus_gallus.GRCg6a genome, we shortlisted reads that simultaneously aligned to two of the five (Flank1, Dup1, Int, Dup2, and Flank2) genomic regions we have defined. We found several reads spanning Flank1-Dup1, Dup1-Int, Int-Dup2, and Dup2-Flank2, which is unsurprising because these regions are adjacent.

We also found five reads (SRR6189090.111279, SRR6189090.56386, SRR6189090.880702, and SRR6189090.387043 and SRR6189090.54824) that span across both Dup1 and Dup2. Dup1 and Dup2 are far apart in their genomic location in Gallus_gallus.GRCg6a assembly. Hence, these reads support a rearrangement that leads to junctions between these two regions. Three of these five reads (SRR6189090.111279, SRR6189090.880702, and SRR6189090.387043) support the junction Dup1 + (Inverted Dup2) (i.e., START-DUP1-END-END-DUP2-START), and the other two reads (SRR6189090.54824 and SRR6189090.56386) support the junction (inverted Dup1) + Dup2 (i.e., END-DUP1-START-START-DUP2-END). The read coverage at these junctions consistently differed between all black-bone and non-black-bone chicken. We identified 34 out of 101 samples analyzed as BBC based on the read coverage across the *Fm* locus junctions.

### Long-read-based resolution of haplotypes at Dup1 and Dup2 regions

Recently, a new high-coverage multi-platform genomic public dataset for the silkie black-bone chicken became available on the ENA (Europen Nucleotide Archive) as part of the Bioproject# PRJNA805080 (we thank China Agricultural University for publishing this data). With a ∼65X (Nanopore) and >660X coverage (PacBio) of the same silkie individual, this dataset is well suited to resolve the haplotypes at the *Fm* locus and identify the exact order of rearrangement. Upon visual inspection of the long-read alignments at the Flank-1-Dup1 and Dup2-Flank2 junctions region, three heterozygous sites were recognized at the Dup1 start region, and Dup2 end region, potentially separating the two haplotypes of Dup1 and Dup2. A detailed blastn-based search of reads from both haplotypes anchored one haplotype to the Flank1 + Dup1 junction and another haplotype to the (inverted Dup2) + Dup1 junction. The Flank1 + Dup1 junction containing haplotype is referred to as Dup1 haplotype-1 (D1H1) and contains the allele T at position 10766895. The (inverted Dup2) + Dup1 junction containing haplotype is referred to as Dup1 haplotype-2 (D2H2) and contains the allele A at position 10766895. We extended these haplotypes toward the end of Dup1 by relying upon overlapping long-reads at haplotype-defining sites. To ensure the robustness of our approach, we required that more than ten sequencing reads should support haplotype-specific alleles at each adjacent site (see **Table S3**).

In the Dup2 region, one haplotype was anchored to the Dup2+ Flank2 junction and another haplotype to the Dup2 + (inverted Dup1) junction. The Dup2 + Flank2 junction containing haplotype is Dup2 haplotype-1 (D2H1) and contains the allele A at position 11476819. The Dup2 + (inverted Dup1) junction containing haplotype is Dup2 haplotype-2 (D2H2) and contains the allele G at position 11476819. We extended these haplotypes toward the start of Dup2 by relying upon overlapping long-reads at haplotype-defining sites. We identified 25 well-resolved, high-confidence haplotype-defining sites spanning from the end of Dup2 to the beginning of Dup2 (see **Table S3**). Haplotype-1 of Dup2 contains reads spanning from the end of the Int region to the start of Flank2, and Haplotype-2 of the Dup2 region spans from the end of the Haplotype-1 of Dup1 (D1H1) region to the beginning of Haplotype-2 of Dup1 (D1H2) region.

### Principal component analysis and admixture

Principal components analysis (PCA) was performed using PCAngsd[59] based on genotype likelihood estimates from ANGSD[60] (Analysis of Next Generation Sequencing Data) v0.935. PCA was done both genome-wide and *Fm* locus region-wise (Flank1, Dup1, Int, Dup2, and Flank2) for the 34 black-bone and all 101 chicken individuals. We used several flags in ANGSD for population structure analysis as follows: -GL 2, -doMaf 1, -minMapQ 30, -minQ 20, -doGlf 2, and -SNP_pval < 1e-6. Genotype likelihood values from ANGSD were used to identify principal components using PCAngsd and genome-wide admixture proportions using NGSadmix[61]. NGSadmix was run for different values of K from K1 to K10 using each K with 15 iterations with flags -minMaf 0.05 and -minInd as specified. We used the online Clumpak[62] web server to find the best K.

### Population genetic analysis

Chinese black-bone breeds have diverged to differing extents from Kadaknath and have limited sample sizes. Hence, we combined the individuals from closely related XBBC, LCEM, and LCMY breeds into a single population representative of Chinese black-bone (CHIN) chicken and JETB, PENC, and GOLD into another population representative of Kadaknath (KADK) (see **Table S4**). We calculated population genetic statistics genome-wide to identify signatures of selection. However, to avoid false positives, we excluded genomic regions (50 Kb windows with <80 percent callable sites) with poor callability. We used the CallableLoci walker of GATK[63] on the bam files to quantify callability. For identifying callable regions from mapped bam files for CHIN and KADK individuals, we used GATK with different flags such as --minMappingQuality 20, --minBaseQuality 20, --minDepth 10, --minDepthForLowMAPQ 20, and --maxFractionOfReadsWithLowMAPQ 20. Furthermore, we filtered 50Kb windows with >0.1 repeat element fraction to rule out the possibility of false positives (see **Fig. S4-6**). We used stringent coverage and quality criteria (-GL 2, -dosaf 1, -baq 1, -C 50, -setMinDepthInd 6, -minInd 3, 4 or 9, -minMapQ 30, -minQ 20, and -doCounts 1) in ANGSD to calculate all population genetic statistics. The list of individuals in each population and the population pairs compared is in **Table S4**. We found that at least 20,000 of the 21,659 50Kb windows had sufficient high-quality data in all populations.

We used the folded site frequency spectrum (SFS) approach implemented in ANGSD to calculate genome-wide population-specific estimates of _π_ (the average pairwise differences), Watterson’s _θ_ (the average number of segregating sites), (Tajima’s D), and Fu and Li’s D. We estimated inter-population genomic differentiation (F_ST_) and divergence (D_xy_) using ANGSD and popgenWindows python script (https://github.com/simonhmartin/genomics_general) for each population pair. The F_ST_ estimates from the two methods were highly correlated (Pearson’s _ρ_= 0.98, p-value < 2.2 e-16) (see **Fig. S7, 8**). We defined F_ST_ outlier regions as 50Kb windows in the top 1 percent of the genome-wide estimates and merged adjoining windows using bedtools. Similarly, the 50Kb windows in the top 10 percent of the genome-wide estimates of D_xy_ were deemed to have high levels of divergence. We identified fixed sites as those sites with F_ST_ > 0.9. We estimate haplotype-based statistics iHS and XP-EHH in the rehh[64] R package with option polarised=FALSE using genotype data phased with SHAPEIT[65].

## Results

### Whole-genome re-sequencing

We generated >25X coverage whole-genome re-sequencing Illumina data for 13 chicken individuals from India. Our collection includes 9 Kadaknath samples as representatives of all three extant morphs Jetblack (JETB n=3), Golden (GOLD n=3), and Pencil (PENC n=3). We also sampled one individual with a golden-pencil-like phenotype (GOPE n=1), one with complete white plumage with black-bone (CROS n=1), one non-black-bone individual where plumage was black (NONB), and one individual of broiler breed (**Fig. S1** and **Table S1**). We compared these Kadaknath chicken genomes with public re-sequencing datasets of chicken breeds. The sampling location (see **Fig. 1a** for black-bone chicken) of individuals analyzed in this study is spread across Asia (for more detail, see **Table S2,** which includes the complete list of all chicken breeds examined). A detailed map of China depicts the locations of all the Chinese black-bone breeds (**Fig. S9**).

**Fig. 1:**
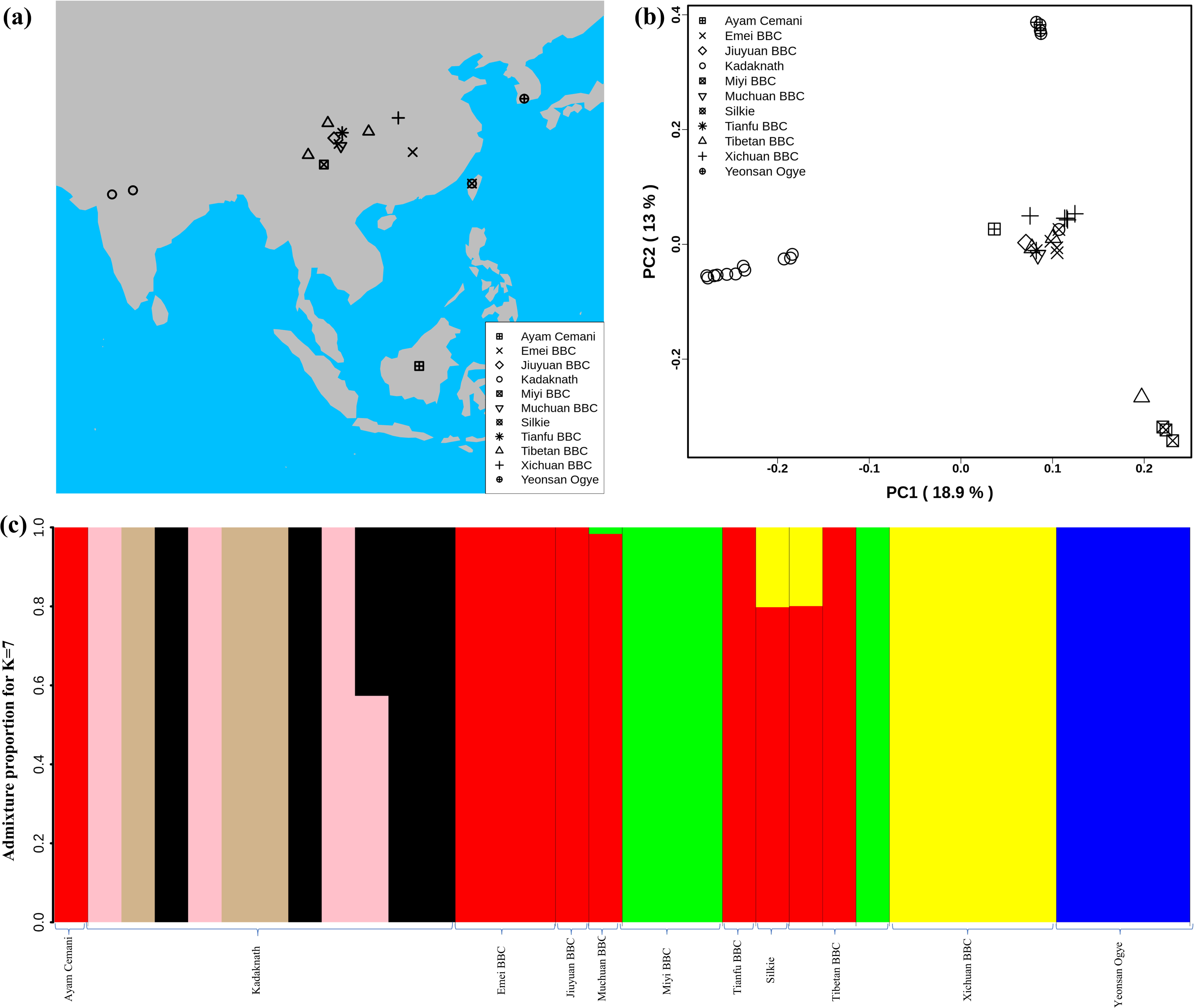
Population structure analysis : **(a)** Geographical locations of different black-bone chicken breeds used in this study are shown on the map using different shapes. The map was generated using rworldmap, map, and mapdata R packages. **(b)** Genome-wide principal component analysis (PCA) reveals the genetic relationship of 34 black-bone chicken individuals. PC1 and PC2 explained 18.9% and 13% variance, respectively. **(c)** Population genetic structure and individual ancestry were estimated using NGSadmix for 34 black-bone chickens from different breeds based on best K=7.

### Kadaknath is a distinct breed of black-bone fowl

The major axes of genetic variation in BBC (PC1:18.9% and PC2:13%) separated the Kadaknath (KADK), Yeonsan Ogye (YOSK), and Chinese BBC (see **Fig. 1b** and **Fig. S10**). We observed that all Kadaknath individuals form a single cluster distinct from other black-bone (from Indonesia, China, and South Korea) (**Fig. 1b** and **Fig. S10**). Similarly, in the genome-wide PCA analysis of 101 individuals, Kadaknath forms a separate cluster from other commercial, native, and BBC breeds in different PC comparisons (see **Fig. S11**). Admixture analysis suggests Kadaknath has sub-structure but remains distinct from the other black-bone breeds at best K=7 (see **Fig. 1c** and **Fig. S12, 13**). In Chinese BBC, the admixture analysis identified three groups that support the clustering pattern in PCA, while YOSK forms a separate cluster (see **Fig. 1c** and **Fig. S10**). We observed lower genetic diversity (_π_ = 0.002, _θ_ = 0.002) in YOSK, while Kadaknath and Chinese BBC have comparable genetic diversity (_π_ = 0.004 and 0.003, _θ_ = 0.003 in both) (**Fig. S14a** and **b**). In contrast to the nuclear genome, the mitochondrial genome haplotype network did not separate Kadaknath from other BBC breeds (**Fig. S15, 16**). The *Fm* locus region on chromosome 20, which codes for the black-bone phenotype, is the defining feature of all BBC. Even after excluding chromosome 20, the PCA of the remaining chromosomes finds that Kadaknath is genetically distinct from other BBC breeds (**Fig. S17**). Hence, the genetic distinctiveness of Kadaknath is spread across the entire genome.

### All black-bone chicken breeds share the same rearrangement junctions at *Fm* locus

In the non-black chicken, the Dup1 (∼127 Kb), Int (∼412 Kb), and Dup2 (∼170 Kb) regions occur in a single copy, are arranged sequentially, and are flanked by Flank1 (∼500 Kb) and Flank2 (∼500 Kb) regions (see **Fig. 2a**). While the Dup1 region contains five protein-coding genes (*EDN3*, *ZNF831*, *SLMO2*, *ATP5E*, *TUBB1*), the Dup2 region probably consists of only long-non-coding RNA genes. All non-black chickens have a single copy of this region, referred to as *\*N* locus (see **Fig. 2a**). The corresponding locus in black chickens is known as the *Fm* locus. The *Fm* locus consists of three different non-paralogous regions (Dup1, Int, and Dup2) that form a complex chromosomal rearrangement in which both the Dup1 and Dup2 regions are duplicated, giving rise to two junctions (**A**) Dup1 + (inverted Dup2) and (**B**) (inverted Dup1) + Dup2. Although the exact ordering of these regions in the rearrangement is not conclusively established, both regions are known to be duplicated due to these regions having a sequencing coverage that is twice the genomic average [13]. The presence of Dup1 + (inverted Dup2) and (inverted Dup1) + Dup2 junctions has been verified in several black chicken breeds [13,14,23]. Based on this information, three possible scenarios have been proposed by earlier studies (see **Fig. 2b**). The \**Fm_2* scenario is supported based on crosses between the black bone and non-black bone chicken [13]. However, both \**Fm_2* and \**Fm_3* scenarios require two rearrangement events compared to a single rearrangement event needed for the \**Fm_1* scenario [23].

**Fig. 2:**
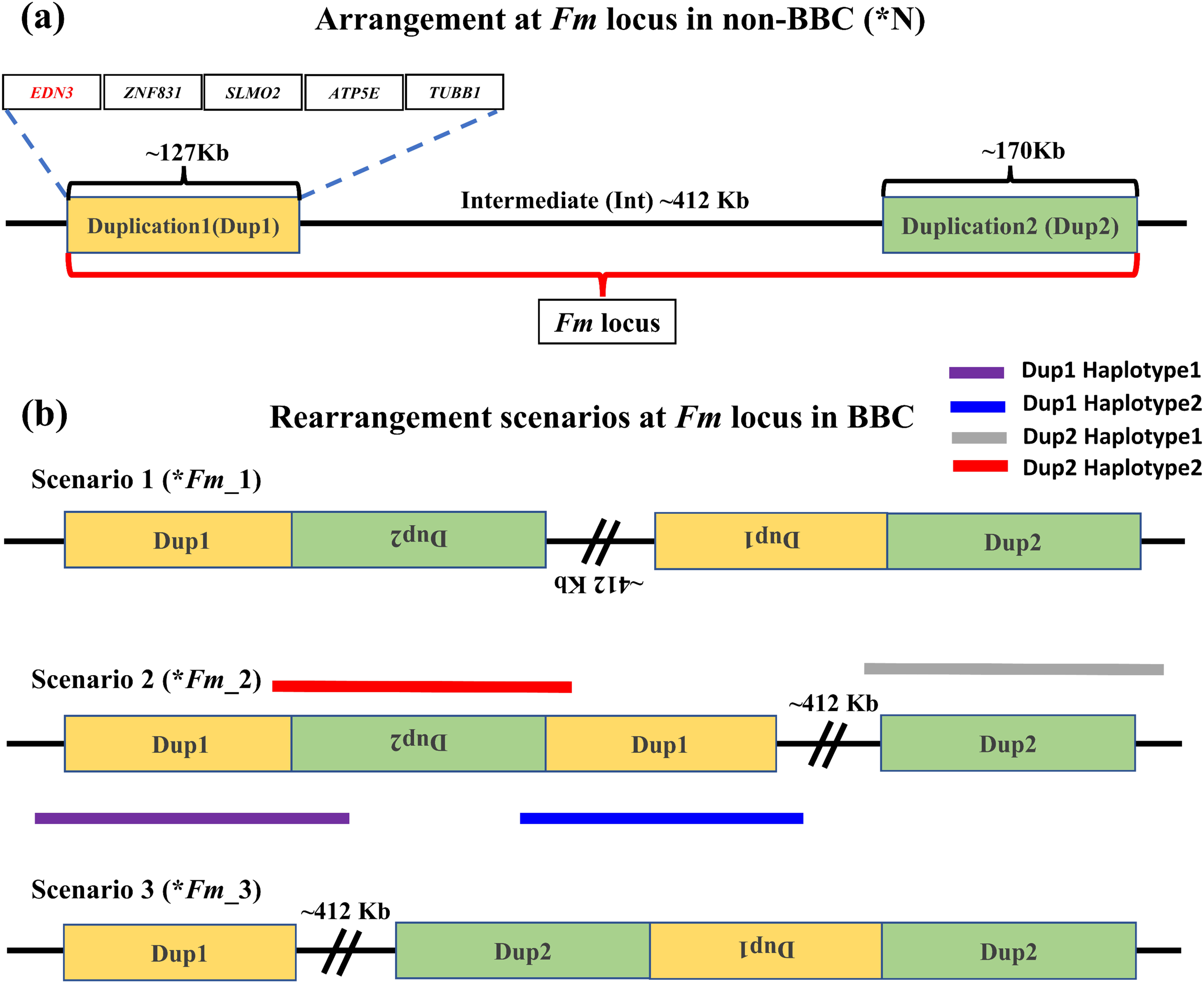
Arrangement of *Fm* locus region in non-BBC and BBC breeds: Two different non-paralogous regions on chromosome 20 are referred to as Duplication 1 (Dup1) and Duplication 2 (Dup2), shown in Gold and light green colors, respectively. The length of the Dup1 region is ∼127 Kb, the Intermediate (Int) is ∼412 Kb, and Dup2 is ∼170 Kb in size. (**a**) These regions are neither duplicated nor rearranged in nonblack bone chicken (*N). The Dup1, Int, and Dup2 regions are collectively referred to as the *Fm* locus. Dup1 contains five genes, *EDN3*, *ZNF831*, *SLMO2*, *ATP5E*, and *TUBB1*, whereas the Dup2 region does not have any protein-coding genes. (**b**) Three possible scenarios (\**Fm_1*, \**Fm_2*, and \**Fm_3*) for *Fm* locus have been proposed in BBC (earlier described in Dorshorst et al. 2011, Dharmayanthi et al. 2017, and Shon et al. 2018). The dark solid purple line represents haplotype-1, which spans across Dup1 from Flank1 to inverted Dup2 (i.e., Flank-1 + Dup1 + (inverted Dup2)). The blue line represents haplotype-2, which spans Dup1 from inverted Dup2 to Int (i.e., (inverted Dup2) + Dup1 + Int).

Distinguishing between these three (\**Fm_1*, \**Fm_2*, and \**Fm_3*) scenarios requires long-range connectivity information such as long-read sequencing data (PacBio, Nanopore, Synthetic long reads, etc.), Hi-C (high-resolution chromosome conformation capture)[66, 67], or Optical Mapping[68]. Both \**Fm_1* and \**Fm_2* contain the same adjacent regions for Dup1 (defined as Haplotype-1 and Haplotype-2 in **Fig. 2b**) inconsistent with \**Fm_3*. Hence, long-range information that can span the entire ∼127 Kb Dup1 region in a single read will be able to distinguish between (\**Fm_1* and \**Fm_2*) vs. \**Fm_3*. However, to differentiate between \**Fm_1* and \**Fm_2* scenarios, we need to span the entire ∼170 Kb Dup2 region in a single read. Spanning the Dup1 or Dup2 regions with PacBio (average read lengths of 10 to 25 Kb) or Nanopore (average read lengths of 10 to 30 Kb) technologies with a single read is challenging but not impossible as reads longer than 1 Mb are possible [69, 70]. Without such individual reads that can span the entire region, it is possible to perform read-based phasing to infer haplotypes that extend to distinct junctions [71].

Using a public long-read dataset, we have inferred distinct high-confidence Dup1 and Dup2 haplotypes (supporting \**Fm_2* scenario). Dup1 haplotypes are anchored by (Flank-1 end)/(Dup2 start) at the Dup1 start and (Int start)/(Dup2 end) at the Dup1 end (**Fig 3**). Each haplotype can be distinguished at 24 sites (see **Table S3**) along the Dup1 region. Each haplotype-defining site shares more than ten long reads with haplotype-specific alleles at adjacent positions. Haplotype-1 (see **Fig. 3a** and **Fig. S18a**, **b**) spans Dup1 from Flank-1 to haplotype-2 Dup2 (D2H2) end, and haplotype-2 (see **Fig. 3b** and **Fig. S18a**, **b**) spans Dup1 from haplotype-2 Dup2 (D2H2) start to Int start. Our inference of Dup1 haplotype-1 (i.e., Flank-1 + Dup1 + (inverted Dup2)) and haplotype-2 (i.e., (inverted Dup2) + Dup1 + Int) sequences rule out the *\*Fm_3* scenario, suggesting that either \**Fm_1* and \**Fm_2* scenario is possible. To verify the correct arrangement, we used the read phasing approach on the Dup2 region, and we could distinguish the two haplotypes at the Dup2 region based on 25 sites (see **Table S3**). Haplotype-1 of the Dup2 region (see **Fig. 4a** and **S19a, b**) spans Dup2 from the end of Int to that start of Flank2, and haplotype-2 (see **Fig. 4b** and **S19a, b**) spans Dup2 from haplotype-2 Dup1 (D1H2) start to haplotype-1 Dup1 (D1H1) end. Our inference of haplotype-1 Of Dup2 (i.e., Int+ Dup2 + Flank2) and haplotype-2 (i.e., start of Dup1 + (inverted Dup2) + Dup1 end) sequences rule out the \**Fm_1* and \**Fm_3* scenarios, suggesting that \**Fm_2* scenario is correct (see **Fig. S20**).

**Fig. 3:**
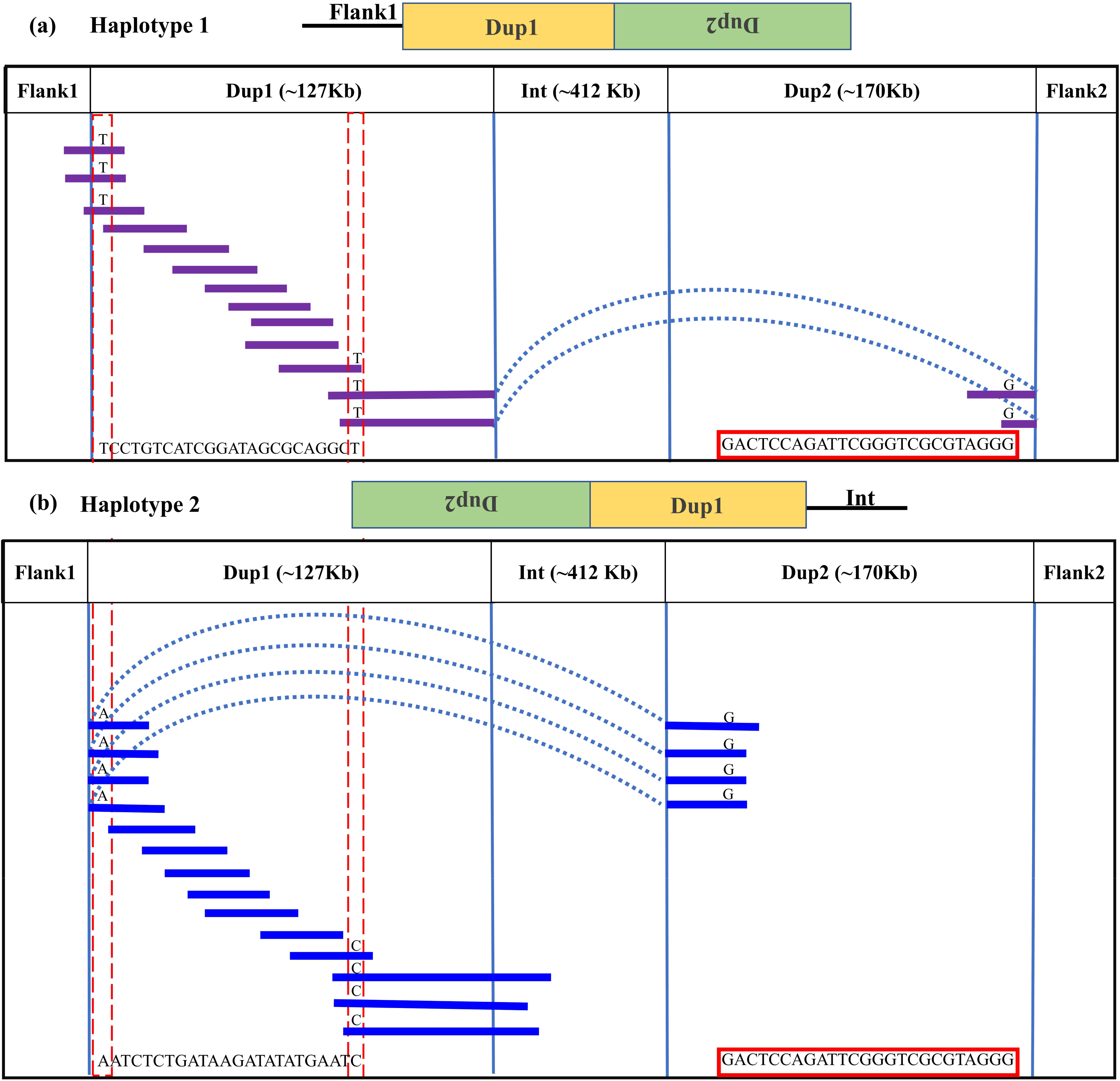
Long-read-based haplotypes resolve the sequence spanning the Dup1 region at *Fm* locus: The two haplotypes spanning the Dup1 region are distinguished by distinct alleles at 24 positions using long sequencing reads. Red dotted vertical boxes highlight the alleles that differ between haplotype-1 and haplotype-2 at the same positions. The alleles at 24 sites that separate these two haplotypes at various positions along Dup1 are presented sequentially between the red dotted boxes. (**a**) Haplotype-1 of Dup1 spans from Flank1 to inverted Dup2 (i.e., Flank-1 + Dup1 + (inverted Dup2)). The purple lines represent overlapping reads of Nanopore containing Dup1 haplotype-1 alleles. The light blue dotted line represents the span of the same read from the end of haplotype-1 of Dup1 to the end of haplotype-2 of Dup2. (**b**) Haplotype-2 spans Dup1 from haplotype-2 of Dup2 to the Int region (i.e., (inverted Dup2) + Dup1 + Int). The blue lines represent the overlapping reads of Nanopore containing haplotype-2 alleles. The light blue dotted line represents the span of the same read from the start of haplotype-2 of Dup1 to the start of haplotype-2 of Dup2. Both haplotypes of Dup1 are connected to the single haplotype of Dup2 (i.e., haplotype-2 of Dup2) but at different ends.

**Fig. 4:**
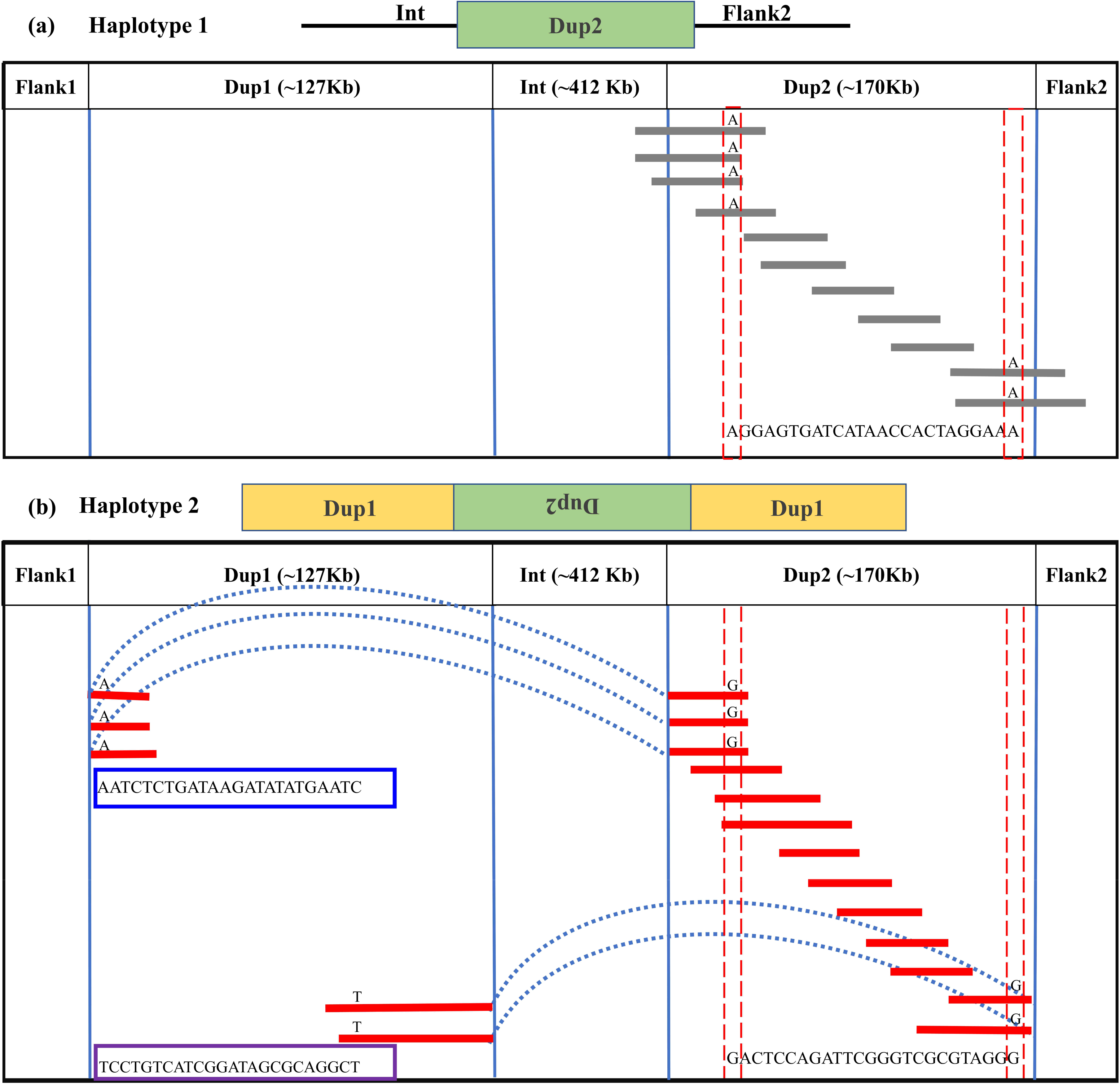
Long-read-based haplotypes resolve the sequence spanning the Dup2 region at *Fm* locus: The two haplotypes spanning the Dup2 region are distinguished by distinct alleles at 25 positions using long sequencing reads. Red dotted vertical boxes highlight the alleles that differ between haplotype-1 and haplotype-2 at the same positions. The alleles at 25 sites that separate these two haplotypes at various positions along Dup2 are presented sequentially between the red dotted boxes. (**a)** Haplotype-1 of Dup2 spans from Int to Flank2 (i.e., Int + Dup2 + Flank2). The grey lines represent overlapping reads of Nanopore containing Dup2 haplotype-1 alleles. (**b**) Haplotype-2 of Dup2 spans from the start of haplotype-2 of Dup1 to the end of haplotype-1 of Dup1 region (i.e., Dup1 +(inverted Dup2)+ Dup1). The red lines represent the overlapping reads of Nanopore containing Dup2 haplotype-2 alleles. The light blue dotted line represents the span of the same reads from the start of haplotype-2 of Dup1 to the start of haplotype-2 of Dup2. Similarly, the same reads span the end of haplotype-1 of Dup1 to the end of haplotype-2 of Dup2.

Conclusive inference regarding which scenario is present in BBC has been extremely challenging due to the large (∼1Mb) size and complexity of the rearrangement. However, in the case of Kadaknath, it has not even been established whether the Dup1 + (inverted Dup2) and (inverted Dup1) + Dup2 Junctions identified in other BBC are present. We have compared the normalized short-read coverage at these junctions to evaluate whether all black-bone chicken breeds share the same rearrangement junctions at the *Fm* locus. The normalized short-read coverage in black-bone chicken (**Fig. 5a-e**) abruptly increases at Dup1 and Dup2 regions, while no such increase occurs in non-black-bone (**Fig. 5f**) chicken. The drastic change in coverage at Dup1 and Dup2 boundaries occurs at the same base in all black-bone chickens (see **Fig. S21-22a-d**). In the case of non-black-bone chickens, no change in coverage occurs at this position (see **Fig. S23**). We also identified the black-bone-specific *Fm* locus junctions Dup1 + (inverted Dup2) and (inverted Dup1) + Dup2 using published PacBio data (see **Fig. 6a, b**, and **Methods**). Short-read coverage spanning these black-bone-specific *Fm* locus junctions is present in black-bone and missing in non-black-bone chicken (**Fig. S24-26**). We can infer that black-bone chicken defining *Fm* locus originated through a single event based on coverage and rearranged junction sequence.

**Fig. 5:**
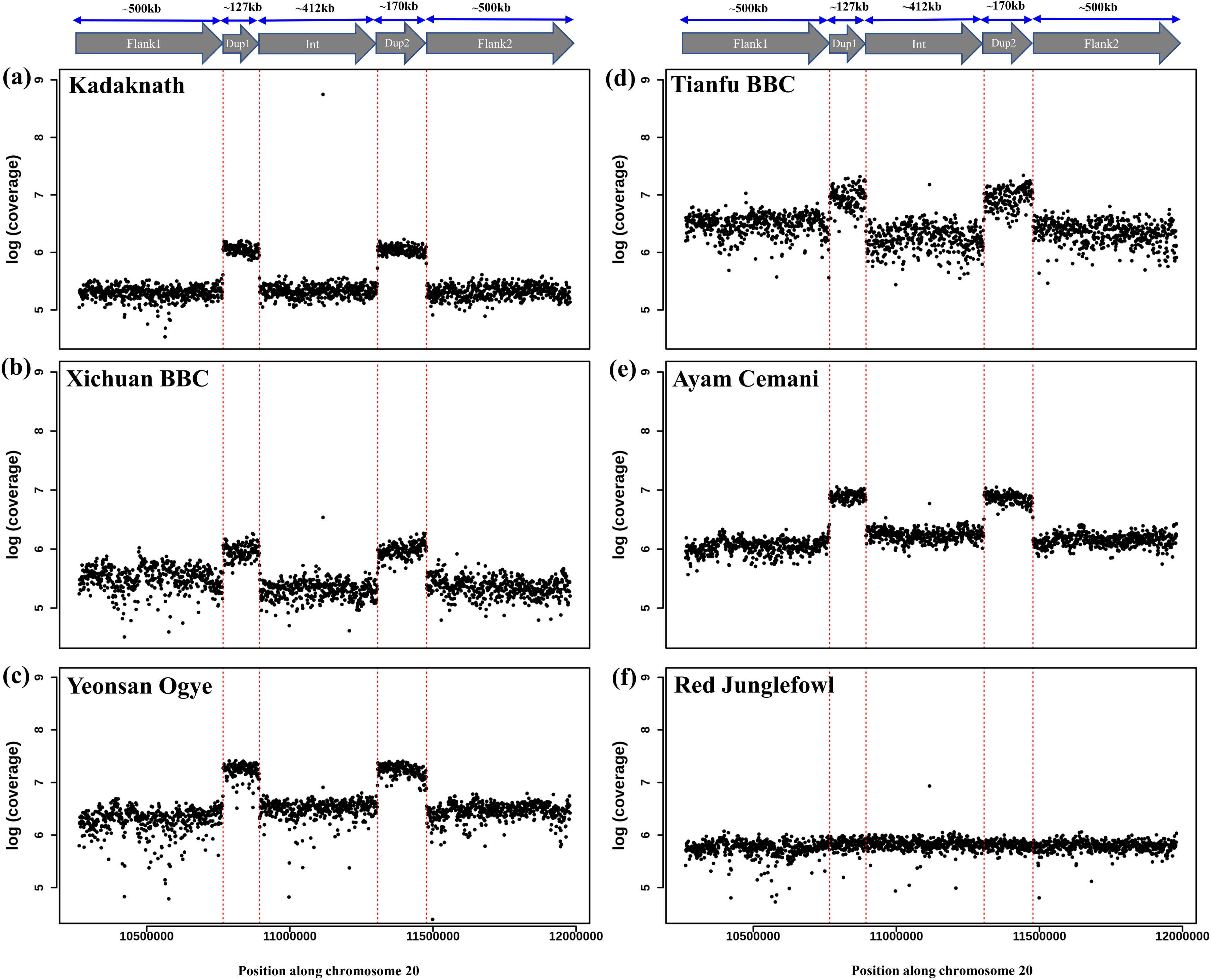
Comparison of *Fm* locus read coverage in black-bone vs. non-black-bone chicken breeds: Read coverage along chromosome 20 at the *Fm* locus is shown in 1Kb sliding windows. Two duplicated genomic loci, Dup1 and Dup2, are denoted by vertical dotted red lines and have a higher coverage in black-bone chicken breeds: **(a)** Kadaknath, **(b)** Xichuan, **(c)** Yeonsan Ogye, **(d)** Tianfu, **(e)** Ayam Cemani than non-black bone chicken: **(f)** Red junglefowl (RJF).

**Fig 6:**
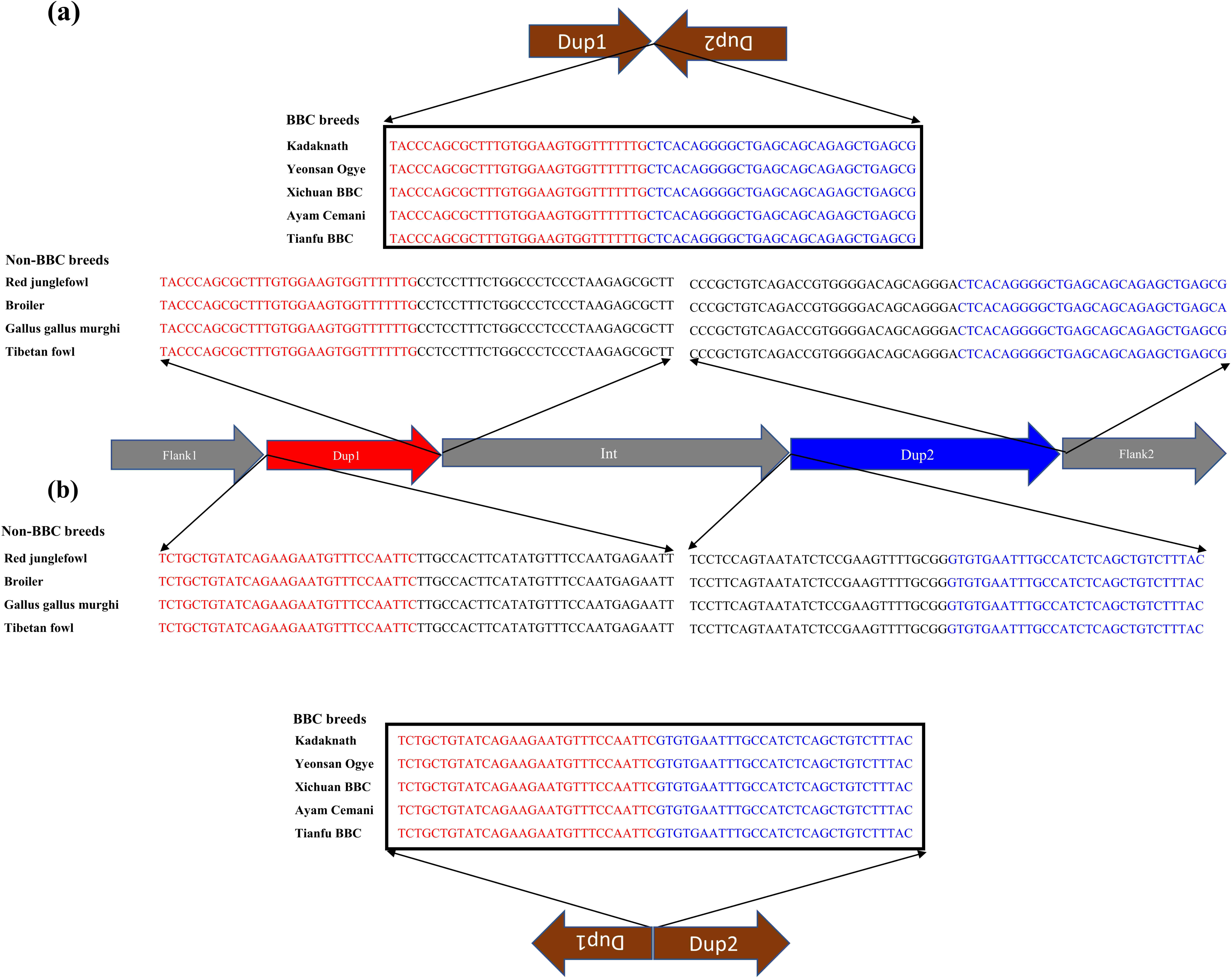
The nucleotide sequence of rearranged junctions at the *Fm* locus in black-bone vs. non-black-bone chicken: **(a)** Dup1 + (inverted Dup2) **(b)** (Inverted Dup1) + Dup2. Brown arrows represent the direction of rearranged junction sequence, which is present as a continuous sequence (verified by their presence in a single read) in BBC breeds. In contrast, non-BBC breeds did not contain these junctions. Sequences highlighted in red and blue represent the Dup1 and Dup2 sequences, respectively. Sequences in black color represent the subsequent nucleotides.

Crossbreeding with native and commercial breeds affects the genome-wide patterns of genetic variation in black-bone breeds (see **Fig. S27-29**). Hence, we focussed on the *Fm* locus to investigate the history of the BBC breeds. The F_ST_ between black-bone and non-black-bone chicken along chromosome 20 is elevated at the Dup1 and Dup2 regions (see **Fig. S30**). However, other population genetic parameters, such as genetic diversity (_π_ and _θ_) and divergence (D_xy_), lack any prominent signatures (**Fig. S31-33**). As evident from the elevated F_ST_, the major axis of genetic variation (assessed using local PCA) in Dup1 and Dup2 regions separates the BBC from non-black-bone chickens (**Fig. S34**). A phylogenetic tree of the SNPs from Dup1 and Dup2 regions also largely separates the BBC breeds from non-BBC breeds (**Fig. S35a** and **35b**). Our evaluation of the genetic differentiation (F_ST_) landscape between BBC breeds found reduced F_ST_ at Dup1 and Dup2 regions compared to the genomic background (see **Fig. S36**). However, comparing individual BBC breeds with non-black chicken breeds showed the opposite pattern with elevated F_ST_ at Dup1 and Dup2 regions (see **Fig. S37, 38**). Hence, the patterns of genetic differentiation (F_ST_), local PCA, and phylogenetic tree also support a common origin of the *Fm* locus in all BBC breeds.

### Isolation by distance pattern suggests dispersal between India and China

Our analysis discovered that the rearrangement junctions in all BBC are identical to the one in Kadaknath and strongly support a common origin for the *Fm* locus. An independent origin for the *Fm* locus would mean separate rearrangement events have created identical junction sequences. Given the lack of repeat sequences at the junctions, such independent origins seem highly unlikely and are not consistent with the genetic relatedness of BBC and non-BBC breeds at the *Fm* locus. Hence, the current distribution of BBC breeds across Southeast Asia needs an explanation. We hypothesized (see a schematic of the proposed scenario in **Fig. S39**) that the dispersal of BBC occurred after a common origin for the *Fm* locus, followed by recent crossbreeding with native and commercial breeds.

To test our hypothesis and delineate the dispersal route of different black-bone breeds, we first evaluated if a pattern of isolation by distance (IBD) is prevalent. The IBD plot shows a consistent increase in genetic distance with an increase in geographic distance (see **Fig. S40**, mantel’s r=0.64, p-value=0.0002). The IBD pattern persisted even after we repeated the analysis using ANGSD/variant-call-based estimates of nucleotide differentiation (F_ST_) after excluding various populations to avoid errors due to auto-correlation and biased estimates from isolated populations (see **Fig. S41-47**). Among the black-bone breeds, Tibetan black-bone (TBTC) and Sichuan black-bone (LCTMJ) chickens are genetically closest to Kadaknath (mean F_ST_∼0.17). The other black-bone breeds that are geographically more distant from Jhabua (LCEM, mean F_ST_=0.21; XBBC, mean F_ST_=0.22; LCMY, mean F_ST_=0.3 and YOSK, mean F_ST_=0.36) occur at increasing genetic distances from Kadaknath (see **Table S5**). The pattern of IBD spanning India and China suggests potentially human-mediated dispersal. We lack conclusive evidence to identify the direction of dispersal. However, analysis of our dataset, even after excluding alleles found in dbSNP[72] (**Fig. S48, 49**), found that Kadaknath has almost twice the number of private alleles than Chinese and Korean BBC breeds and suggests an India-to-China dispersal. More extensive fine-scale sampling may provide a definitive answer regarding the direction and timing of the dispersal.

### Genome-wide signatures of selection in Kadaknath chicken

We screened the genome for selection signatures to identify all Kadaknath-specific regions by comparing Kadaknath with Chinese BBC. We also compared Kadaknath and Chinese BBC (XBBC) with the Korean Yeonsan Ogye (YOSK) to identify population-specific signatures. Despite the sizeable geographic separation, the genome-wide mean F_ST_ between KADK and CHIN is 0.16, making it amenable to identifying selection signatures (see **Fig. 7a** and **Table S6**). A pairwise comparison between KADK and CHIN revealed 137 genic regions in the top 1% F_ST_ windows (see **Table S7** and **Fig. S50**-**82**). We further shortlisted candidates using the additional criteria that genetic diversity (_π_ and _θ_) and integrated haplotype score (iHS) should be strikingly different between KADK and CHIN. Two prominent signatures from these shortlisted regions indicative of selective sweeps in the KADK breed are evident on the 20^th^ chromosome in the vicinity of the Dup1 region. The first region (R1) of ∼300 Kb is ∼160 Kb before Dup1, and the second region (R2) of ∼70 Kb is ∼0.7 Mb before Dup1. In both regions, the genetic diversity (_π_ and _θ_) and Tajima’s D (_τ_) are strongly reduced in KADK compared to CHIN (see **Fig. 7b** and **Fig. S69**). The elevated iHS in KADK compared to CHIN and the positive value of extended haplotype homozygosity (XP-EHH) supports a selective sweep in KADK. This region also occurs in the top 10% genome-wide genetic divergence (D_xy_) windows (see **Fig. S83**).

**Fig. 7:**
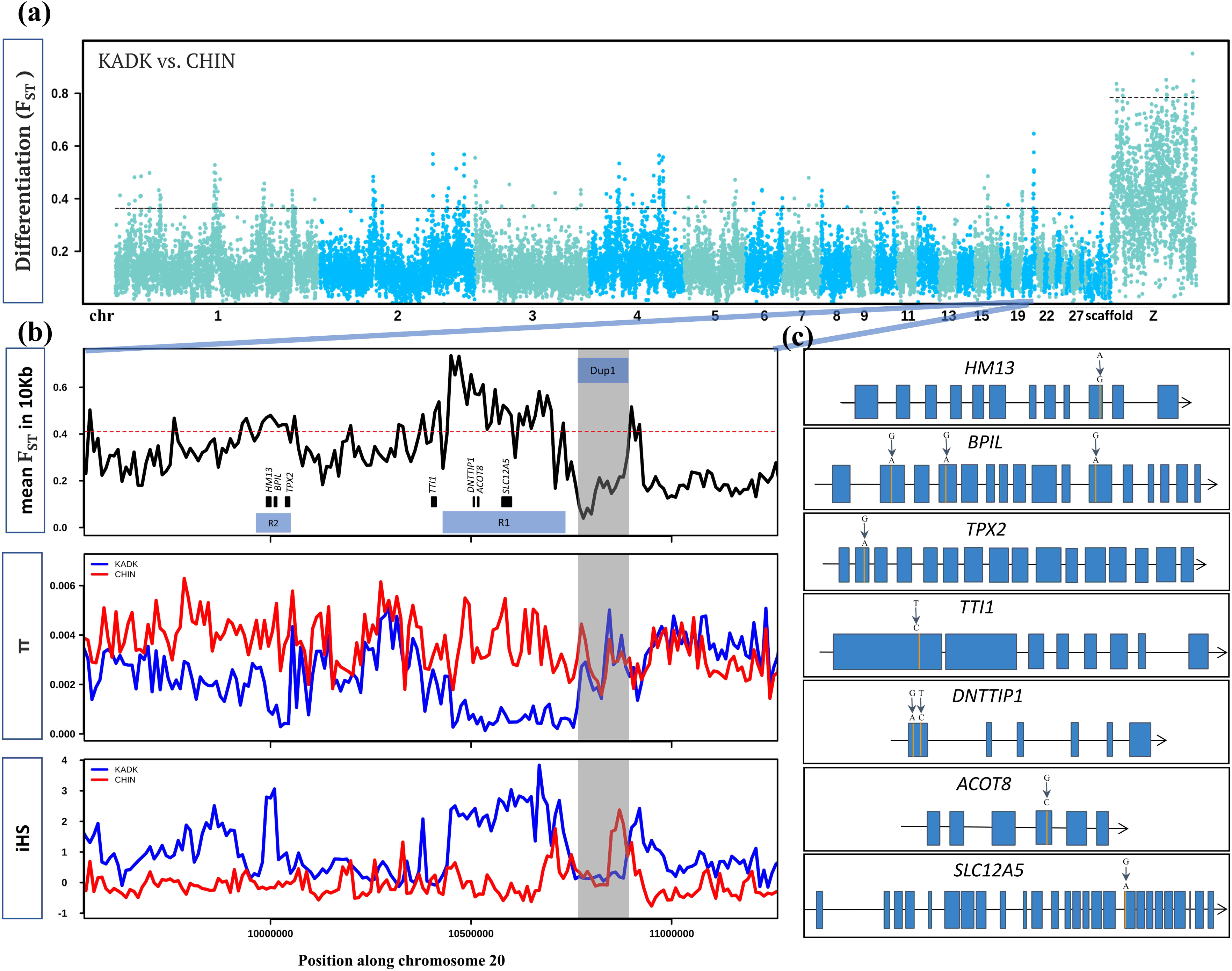
Within and between population comparison of KADK with CHIN: **(a)** Genome-wide landscape of pairwise genetic differentiation (F_ST_) between KADK and CHIN in 50Kb non-overlapping windows. The dark slate gray and deep sky blue colors represent the alternative chromosomes. The dotted horizontal black line marks the 99th percentile outlier of estimated F_ST_ for autosome and the Z chromosome, respectively. **(b)** The highest F_ST_ region with a prominent pattern occurs on chromosome 20. The panels from top to bottom show estimates of F_ST_ in 10Kb non-overlapping windows within population pairwise nucleotide diversity (_π_) and integrated haplotype score (iHS) with major allele as ancestral and minor allele as derived. The genes are represented as black boxes with names. The horizontal red dotted line marks the 99th percentile 10Kb F_ST_ outlier. The transparent grey color highlights the Dup1 region in all panes in the B panel, while R1 and R2 represent the selective sweep regions. **(c)** Non-synonymous changes (shown above the exon) within genes in the regions R1 and R2 are denoted by vertical orange lines. Blue boxes are the exons connected by black lines representing introns.

In the first region (R1), Deoxynucleotidyltransferase Terminal Interacting Protein 1 (*DNTTIP1*), Acyl-CoA Thioesterase 8 (*ACOT8*), and solute carrier family 12 member 5 (*SLC12A5*) genes contain non-synonymous changes. The TELO2-interacting protein one homolog (*TTI1*), which occurs near the first sweep region (R1), also has a non-synonymous change (**Fig. 7c** and **Table S8, 9**). The second sweep region (R2) contains three genes with non-synonymous changes: Histocompatibility Minor 13 (*HM13*), Bactericidal/permeability-increasing protein-like (*BPIL*), and Targeting protein for Xklp2 (*TPX2*). The most striking differences were in the *BPIL* gene, which has three fixed sites with non-synonymous changes. Of these changes, two alleles are unique to the KADK population (see **Fig. S84**). The first non-synonymous change (C**G**G->C**A**G, R->Q) occurs at position 69 in exon-2. The second non-synonymous change (**G**AG->**A**AG, E->K) is within the first BPI superfamily protein domain at position 159 in exon-4. The third and last non-synonymous change (C**G**C->C**A**C, R->H) is in the second domain at position 405 in exon-11. We also found that *HM13, SLC12A5,* and *DNTTIP1* have novel Kadaknath-specific changes.

The Korean Yeonsan Ogye (YOSK) breed is closer to the Chinese XBBC (mean F_ST_ = 0.26) than to Kadakanth (mean F_ST_=0.3). However, the genome-wide mean F_ST_ is still low enough to distinguish selection signatures. A comparison of YOSK with KADK confirmed that the R1 and R2 regions are specific to Kadaknath (see **Fig. S85** and **Table S6**). Surprisingly, our comparisons with YOSK identified a third region (R3) of ∼570 Kb, which is ∼2.16 Mb before Dup1 with a sweep specific to YOSK (see **Fig. S86** and **Table S9**). Local PCA of the *Fm* locus, R1, and R2 regions demonstrates that Flank1, R1, and R2 regions have changed only in Kadaknath and not in other BBC (**Fig. S87-89**). A careful examination of the entire chromosome 20 failed to identify any other sweep regions (see **Fig. S90**). Similar to the sweep in KADK, we found two genomic regions with signatures of a sweep in the CHIN population (see **Table S9**). The sweep region on Chr 4 contains the *PCDH7* gene, and the region on Chr 9 contains the *COL4A3* and *MFF* genes (see **Fig. S53, 58**). However, none of these genes have any non-synonymous fixed differences.

## Discussion

### All BBC breeds share a common origin of the *Fm* locus

As part of this study, we have generated the first whole-genome dataset of the Kadaknath breed spanning all three morphs. Hence, by comparing our data with public datasets, we can evaluate the relationship between all BBC breeds. Our comparison of the *Fm* locus junction region in genome sequencing data from BBC breeds conclusively establishes a common origin for the complex chromosomal rearrangement[14]. The following four lines of evidence support this conclusion: (1) The short-read coverage along the *Fm* locus, (2) the high sequence identity of the rearranged junctions across the BBC breeds, (3) the local and phylogenetic relationship between chicken breeds at the *Fm* locus, and (4) the patterns of pairwise genetic differentiation between BBC and non-BBC breeds.

### Spread of BBC across Southeast Asia

The earliest records of black-bone chicken have been found in the writings of Marco Polo, the Venetian explorer-writer who traveled through Asia [13]. The Compendium of Materia Medica or Bencao Gangmu, compiled and edited by Li Shizhen and published in the late 16th century, attributes various medicinal properties to BBC [73]. In Korea, black-bone chicken is thought to have had a royal connection, and “Dongui Bogam”, a traditional Korean medical encyclopedia compiled & edited by Heo Jun in 1613, records the medicinal use of BBC [23]. While an earlier study notes that “In 1635 AD, the finding of chickens with black meat (typical of fibromelanosis, FM mutation) in Mozambique suggested direct introductions from India.”[2] we could not find literature on how Kadaknath ended up in Jhabua, India.

Jhabua is close to the ancient port cities of Bharuch (260 Km) and Lothal (300 Km) on the west coast of India. Given the proximity to port cities and the prevalence of BBC only in Western India (and the spread of BBC to Africa), we suspected that Kadaknath may have traveled through a marine route. Ancient sea trade between India, Korea, and other parts of Asia are well documented [74]. Moreover, the movement of domesticated breeds during the period of colonialism was also facilitated by the common rule of large parts of Southeast Asia by various European powers [75].

Historical records are patchy, prone to error/obfuscation, and fail to provide conclusive information about the spread of BBC across Asia. Hence, we evaluated the relationship between BBC breeds and patterns of isolation by distance to infer the ancient dispersal route. Although each BBC breed has considerable genetic distinctiveness, population genetic analyses support the common heritage of all BBC breeds and reveal a trend of isolation by distance (IBD). While IBD patterns are well established in Chinese native chicken breeds[76], our dataset spans BBC breeds from India, China, Indonesia, and Korea. The sample size of some BBC breeds in our analysis is limited, and the sampling does not cover the entire geographic distribution of some breeds. However, our analysis achieves genome-wide pan-BBC breed sampling by including all the major breeds from Asia.

The BBC chicken from Tibet is genetically closest to the Kadaknath breed and suggests that the old Tibet-Nepal salt trade route or the maritime silk route may have facilitated the spread of BBC. Interestingly, the Tibetan BBC is genetically more similar to Kadaknath than some Chinese BBC breeds. Unfortunately, we lack BBC samples from Nepal. However, we believe BBC poultry in Nepal is Kadaknath chicken, recently imported from India. Hence, the ancestral stock of the BBC spread across Asia may not be currently available. Changes in trade routes and the introduction of commercial poultry breeds limit our ability to trace the historical prevalence of BBC breeds.

### Did black-bone chicken originate in India?

The lack of data from India and the considerable interest in BBC breeds in Europe, China, and Korea has meant that studies have focussed mainly on non-Indian BBC. Hence, Southern China and Tibet are considered the source of all BBC breeds[25, 77]. However, despite a traditionally restricted geographic distribution, the Kadaknath breed has nucleotide diversity comparable to Chinese BBC breeds and much higher than the Korean BBC. The number of private alleles identified in Kadakanth is also greater than in Chinese and Korean BBC breeds. Moreover, the mean genome-wide F_ST_ of 0.11 between Jhabua and Bhopal is comparable to the differentiation between some Chinese BBC breeds. The presence of three distinct morphs within Kadaknath chicken suggests phenotypic diversity derived either from commercial breeds or existing variations within Kadaknath. The extent of phenotypic diversity within BBC breeds from China, and Korea is not documented to allow a fair comparison. The high genetic diversity in Kadaknath supports the potential origin of all BBC in Jhabua, India. The export of black chicken from India to Africa in ∼1600 AD also supports that BBC was present in ancient India[2].

While we cannot conclude whether BBC had an Indian, Chinese, or more Southeast Asian origin, our data suggest that any of these sources are plausible. More widespread geographic sampling and analysis of allele-sharing patterns may provide a definitive answer regarding the origin of BBC. Unlike the initial domestication of the chicken, which may have occurred independently[2, 47] in several locations, the BBC has a single origin linked to the rearrangement at the *Fm* locus. Irrespective of the source of all BBC, we identify several Kadaknath-specific genetic changes. Hence, the genetic distinctiveness of the Kadaknath breed has long diverged from other BBC breeds and is a result of its unique heritage sustained by the Bhil and Bhilala tribal communities of Madhya Pradesh. The beliefs and practices of tribal communities may have contributed to the domestication and conservation of Kadaknath[78].

### Conclusive resolution of the complex chromosomal rearrangement

Earlier studies of the *Fm* locus have proposed three possible scenarios for the complex chromosomal rearrangement. Although the *\*Fm_2* scenario was favored based on crosses between BBC and non-BBC breeds, the correct scenario was not established by previous studies as the genome assembly of this region is challenging. We use a haplotype phasing approach relying upon published long-read datasets to conclusively resolve the correct arrangement to be *\*Fm_2* scenario at the *Fm* locus. In the *\*Fm_2* scenario, the distal region (Dup1+Int+Dup2+Flank2) is similar to the *\*N* arrangement found in non-BBC breeds. However, the proximal region (Flank1+Dup1+(inverted Dup2)) has a very different arrangement than the non-BBC one. In contrast to the *\*Fm_2* scenario, the proximal region would be similar to the *\*N* arrangement in *\*Fm_3* scenario. In the *\*Fm_1* scenario, only the first occurrence of Dup1 and last occurrence of Dup2 are similar to the *\*N* arrangement. Hence, in the *\*Fm_2* and *\*Fm_3* scenarios recombination with *\*N* arrangement may be easier (i.e., recombination is supressed only in the inverted ∼127Kb (Dup1) or 170Kb (Dup2) region) compared to *\* Fm_1* scenario (i.e., recombination is supressed in ∼709Kb (inverted (Dup2+Int+Dup1)) region). The ease of recombination with *\*N* arrangement could explain the prevalence of *\*Fm_2* scenario which requires two rearrangement events.

### Selective sweep near *Fm* locus may be a consequence of linkage

Our analyses identified two genomic regions (R1 and R2) in the vicinity of the *Fm* locus with prominent signatures of a selective sweep in the KADK population. The most promising candidate gene in this region that may have been the focus of selection is the *BPIL* gene which has accumulated three non-synonymous changes. The *BPIL* gene is part of the innate immune defense system, which binds and neutralizes lipopolysaccharides (LPS) from the outer membrane of Gram-negative bacteria[79–81]. In the chicken genome, five *BPI*-like genes occur near *BPI*[82], which help arrest bacterial growth, are prominent in phagocytosis by neutrophils, and work as a bactericidal protein[83, 84]. In chickens, defense against gram-negative bacteria may be especially important as they lack the complement *C9* gene required to effectively eliminate pathogens through the Membrane Attack Complex (MAC) formation[85]. Notably, another BBC breed from Korea, YOSK, has signatures of selection at the TLR4 gene involved in detecting Gram-negative bacteria[21]. Hence, various immune genes may have been selected in different chicken breeds to protect against gram-negative bacteria.

Several other genes with non-synonymous changes occur in R1 and R2 genomic regions. Although we cannot identify the functional consequences of these changes, the strong signature of selection and KADK-specific protein-coding alterations suggest phenotype-altering ability. The proximity of R1 and R2 regions to the *Fm* locus indicates that selective sweep may be due to close physical linkage with the *Fm* locus. Such co-selection of traits due to linkage and pleiotropy has occurred during animal domestication, including in chicken[6,86,87]. Hitchhiking of alleles can increase the frequency of both beneficial and mildly deleterious alleles [88–95]. Whether the changes in KADK are beneficial or deleterious will need to be investigated in future studies.

The genomic co-occurrence of agronomically important traits in domesticated plants and animals is also known[96–105]. The alleles causing phenotypic changes can occur in closely linked genes and will undergo selection for any of the phenotypes being favored by the breeding process. Notably, the R1 and R2 selective sweeps are found only in KADK, R3 in YOSK, and none in Chinese BBC breeds. Hence, the genetic variants and the selective sweep may represent recent events after the BBC breeds separated from each other. Comparing different BBC breeds provides snapshots of the selection process and the build-up of genetically linked co-selected allelic changes and may help understand the hitchhiking process during domestication.

Genome-wide data generated as part of this study and comparative analysis with other BBC breeds establishes the genetic uniqueness of Kadaknath that extends beyond the *Fm* locus. We also identify specific genes with signatures of selection that are likely to be responsible for the Kadaknath-specific phenotypes. Co-selection of genes that are in linkage, as shown in Kadaknath, is widespread in domestic species[6,86,96–99]. Such clustering of linked alleles under selection is favored in low recombination regions and near chromosomal rearrangements. Our work exemplifies the interaction of artificial selection and chromosomal rearrangement linked traits in domesticated species.

## Author contributions

Conceptualization: SSS, AS, NV

Methodology: SSS, AS

Investigation: SSS, AS, NV

Visualization: SSS, AS

Supervision: NV

Writing—original draft: SSS, AS

Writing—review & editing: SSS, AS, NV

## Data and materials availability

All primary sequencing data are available from the SRA under BioProject# PRJEB51457 and PRJEB58678. All **Supplementary information**, including other associated data and scripts used for analysis, are provided in an easy-to-browse format: https://github.com/ceglabsagarshinde/Kadaknath_Project.

## Ethics statement

The study was approved by the Institutional Ethics Committee (IEC) of the Indian Institute of Science Education and Research, Bhopal, vide reference number IISERB/IEC/Certificate/2018-11/03 dated 8^th^ June 2018.

## Contribution to field

Domestication of chicken occurred in Asia before spreading across the world. Large-scale population genomics has helped unravel the history of domestication and the genetic contribution from the wild relatives of chickens. Yet, different chicken breeds’ distinctiveness and adaptation to local environments have received limited attention, especially in India. Hence, population genomic datasets of indigenous breeds from India will help understand the spread of domesticated chicken and breed formation. Some indigenous breeds are experiencing a decline in numbers and need conservation. We focussed on one of the most distinctive indigenous chicken breeds, the Kadaknath chicken, which is well known for its black bone and meat. Commonly known as Kali masi or karaknath, the unique taste and use in traditional medicine have made this breed extremely popular across India.

Black-bone chicken breeds are found in several other countries and have fascinated breeders and geneticists alike. We compared the Kadaknath genome with other black-bone chicken breeds and found that all black-bone breeds share the same rearrangement junctions at the *Fm* locus, responsible for the intense black pigmentation throughout the body. Notably, we use public long-read high-coverage datasets to conclusively resolve the correct arrangement of the complex chromosomal rearrangement at the *Fm* locus.

