## Supplementary Figures for "Hitchhiking of bactericidal/permeability-increasing protein-like gene with the fibromelanosis locus in Kadaknath black-bone-chicken"

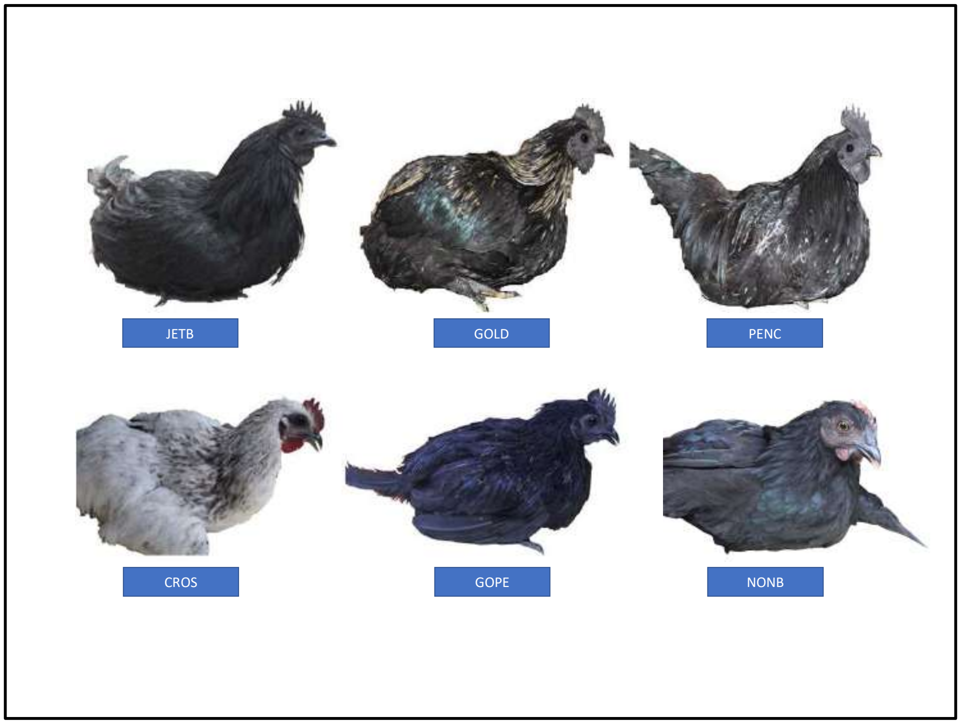


**Fig. S1.** The first row has photographs of the Kadaknath morphs Jetblack (JETB), Golden (GOLD), and Pencil (PENC). The second row has photographs of CROS, which has black internal organs while feathers are white with a red comb, Golden-Pencil-like phenotype (GOPE) and non-black individual (NONB) which have internal organs are non-black while the feathers are black with red comb color.

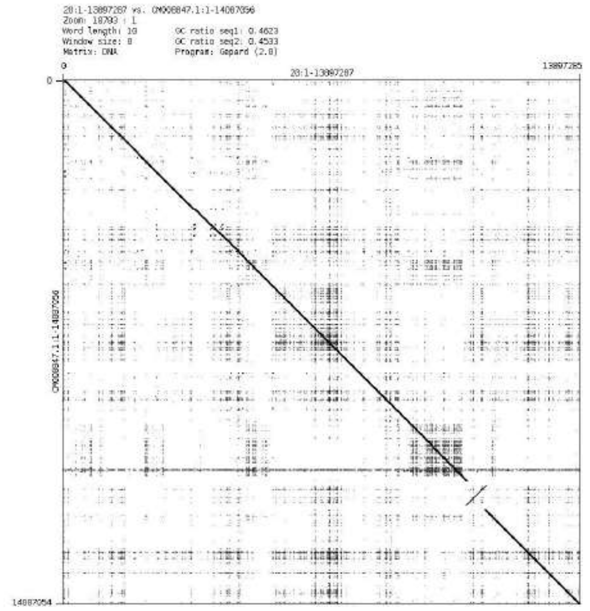


**Fig. S2.** Dot plot between chromosome 20 Galgal6 genome assembly with chromosome 20 (CM008847.1) Yeonsan ogye genome. The figure was generated using Gepard2.1. At *Fm* locus region in the dot plot the rearranged region can be seen.

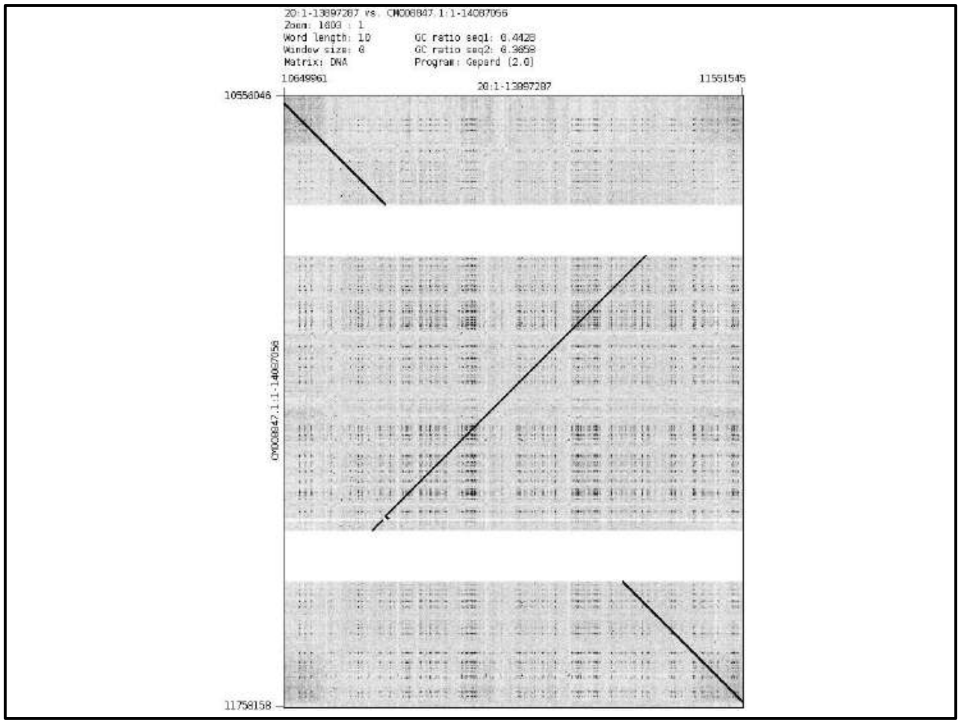


**Fig. S3.** Dot plot between chromosome 20 Galgal6 genome assembly with chromosome 20 (CM008847.1) Yeonsan ogye genome. The figure was generated using Gepard2.1. Zoomed view of dot plot at *Fm* region.

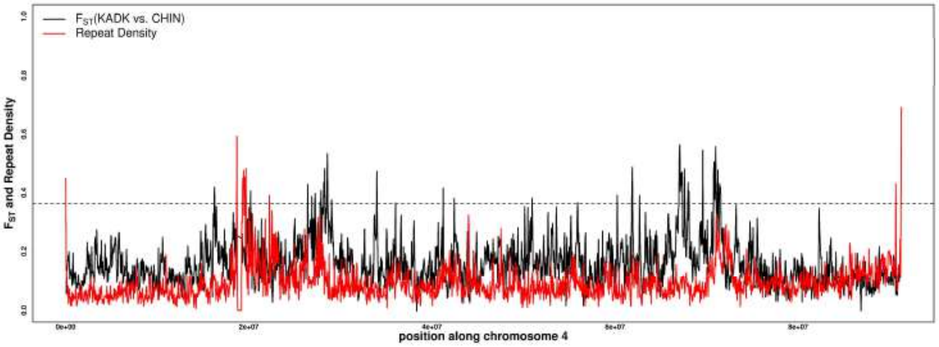


**Fig. S4.** Repeat density and $F_{ST}$ along with chromosome 4 are shown. A horizontal black dotted line represents the 99 percentile $F_{ST}$ threshold. The solid red line represents repeat density in Galgal6 genome assembly calculated in 50Kb windows, and the solid black line represents a pairwise $F_{ST}$ comparison between KADK and CHIN population using 50Kb windows.

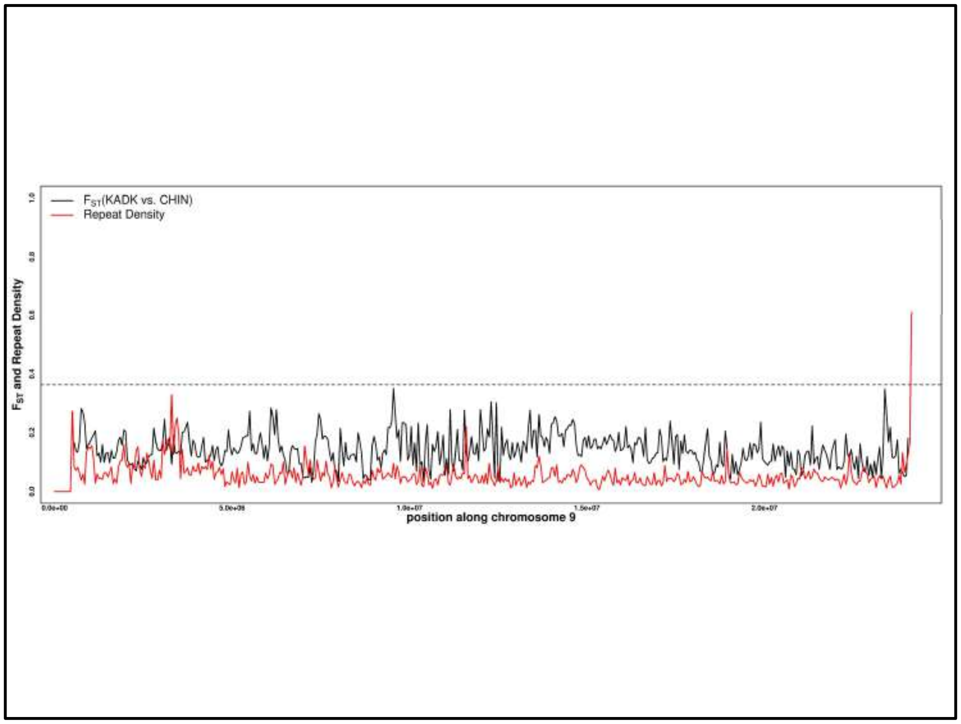


**Fig. S5.** Repeat density and $F_{ST}$ along with chromosome 9 are shown. A horizontal black dotted line represents the 99 percentile $F_{ST}$ threshold. The solid red line represents repeat density in Galgal6 genome assembly calculated in 50Kb windows, and the solid black line represents a pairwise $F_{ST}$ comparison between KADK and CHIN population using 50Kb windows.

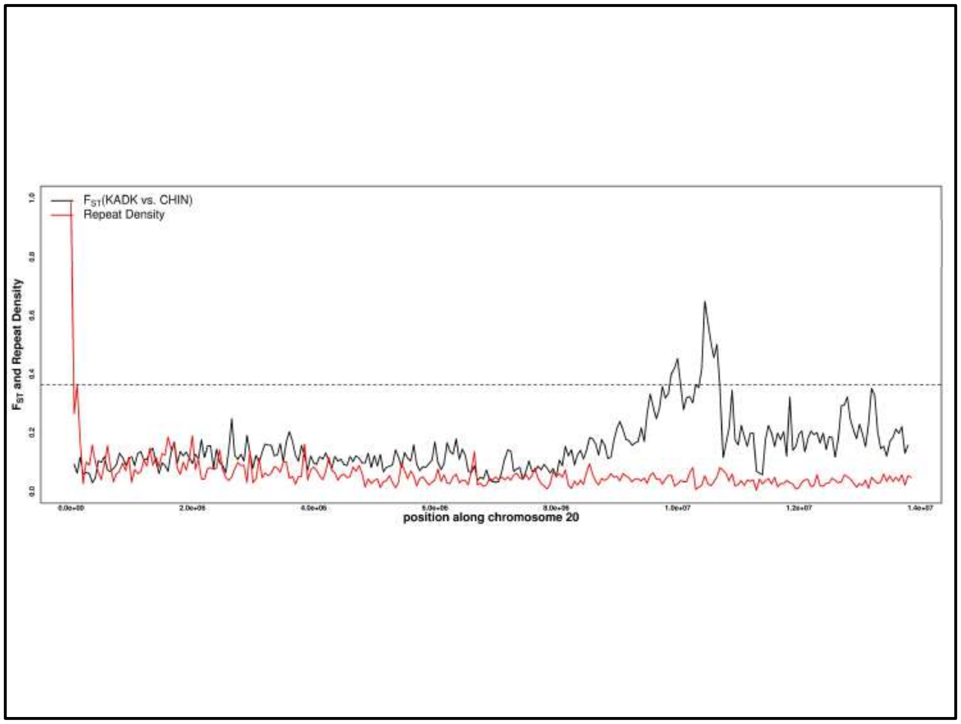


**Fig. S6.** Repeat density and $F_{ST}$ along with chromosome 20 are shown. A horizontal black dotted line represents the 99 percentile $F_{ST}$ threshold. The solid red line represents repeat density in Galgal6 genome assembly calculated in 50Kb windows, and the solid black line represents a pairwise $F_{ST}$ comparison between KADK and CHIN population using 50Kb windows.

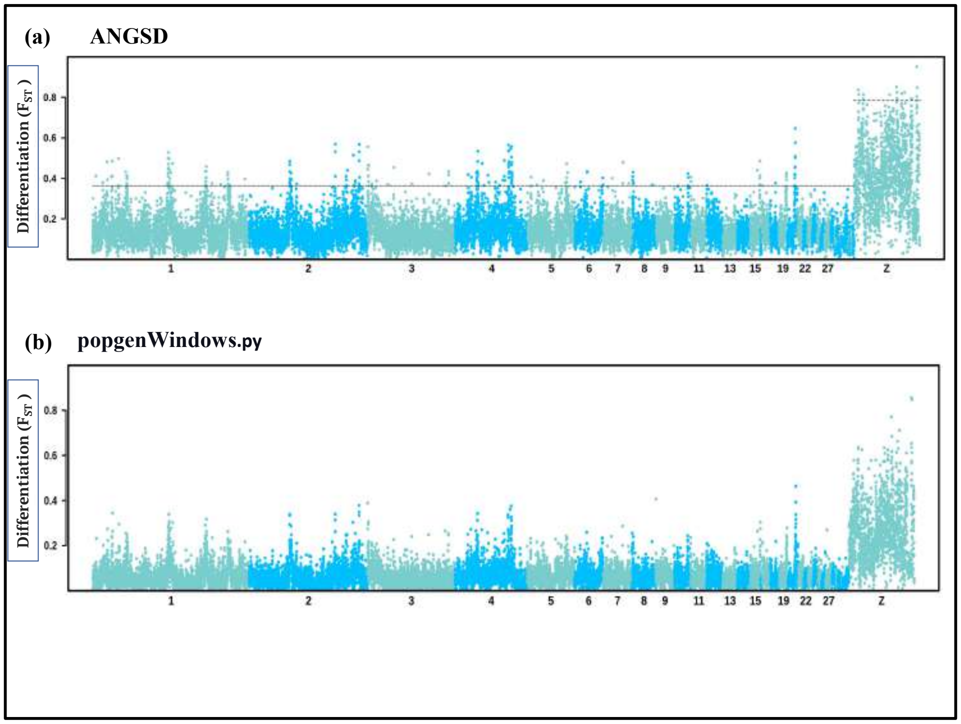


**Fig. S7.** Pairwise $F_{ST}$ between KADK and CHIN population calculated using **(a)** ANGSD **(b)** popgenWindows.py. The dark slate gray and deep sky blue colors represent the alternative chromosomes, dotted horizontal black line marks the 99th percentile outlier $F_{ST}$ estimated for autosome and Z chromosome.

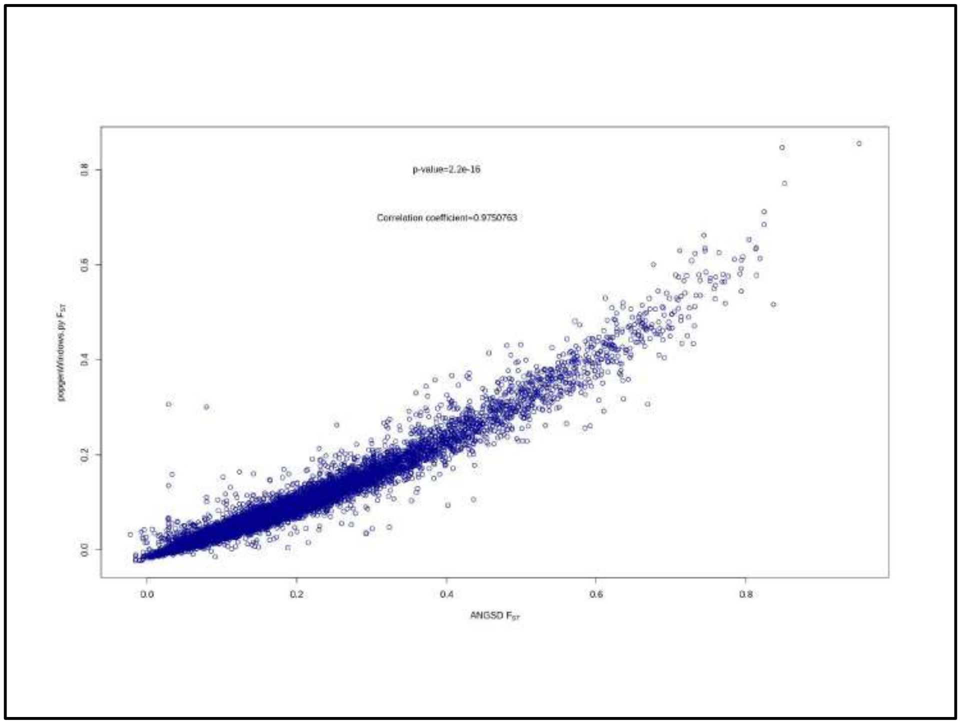


**Fig. S8.** Pearson correlation coefficient test on the $F_{ST}$ comparison from ANGSD and popgenWindows.py method. $F_{ST}$ from popgenWindows.py at Y-axis and $F_{ST}$ from ANGSD at X-axis has a significant positive correlation with a correlation coefficient= 0.975 shown in blue circles.

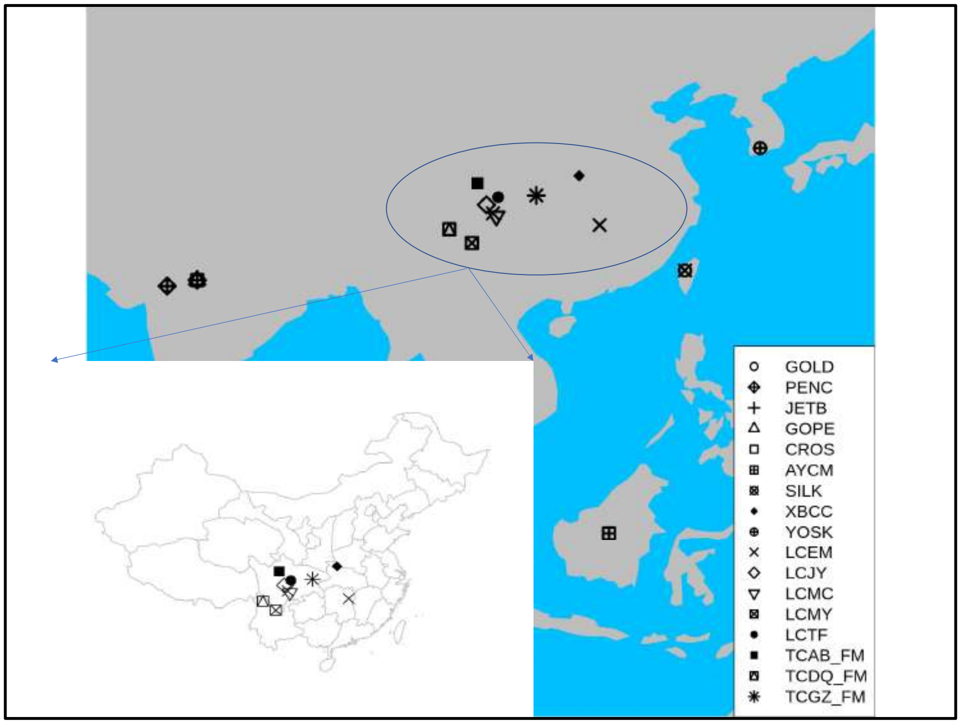


**Fig. S9.** Geographical locations of black-bone chicken breeds used in this study. The map was generated using rworldmap, map, and mapdata packages implemented in R. Different shapes represent different breeds. All breeds in oval shape circles belong to China, and the zoomed view of the China map represents the sample location of each breed.

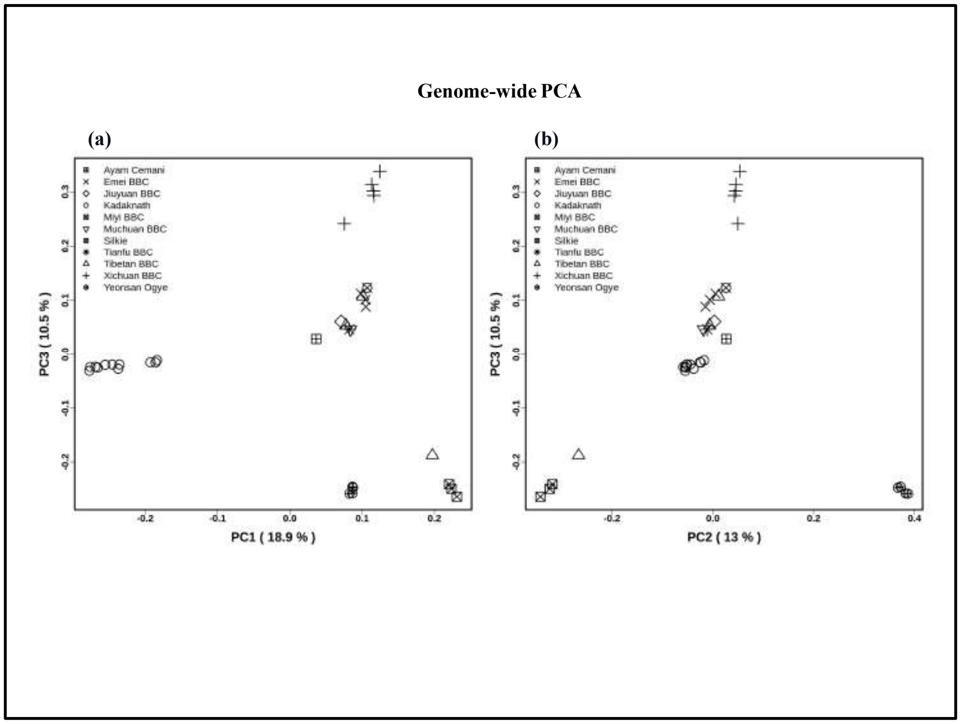


**Fig. S10.** Genome-wide principal component analysis (PCA) 34 black-bone chickens with the two principal groups **(a)** PC1 and PC3 and **(b)** PC2 and PC3 . Each breed is shown in black color with different shapes for each breed.

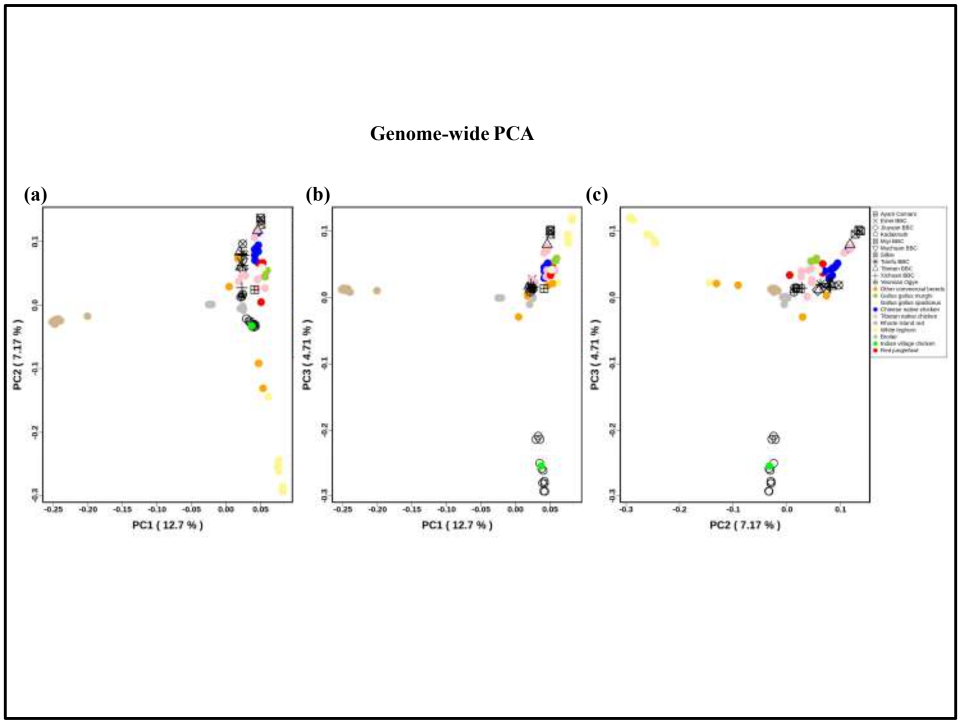


**Fig. S11.** Genome-wide PCA plot generated for 101 chicken individuals: **(a)** PC1 and PC2 explained 12.7% and 7.17 % variance, respectively. **(b)** PC1 and PC3 explained 12.7% and 4.71% variance, respectively. **(c)** PC2 and PC3 explained 7.17% and 4.71% variance, respectively. Non-black-bone breeds are shown by different colors. All black-bone breeds are shown in black color with different shapes for each breed

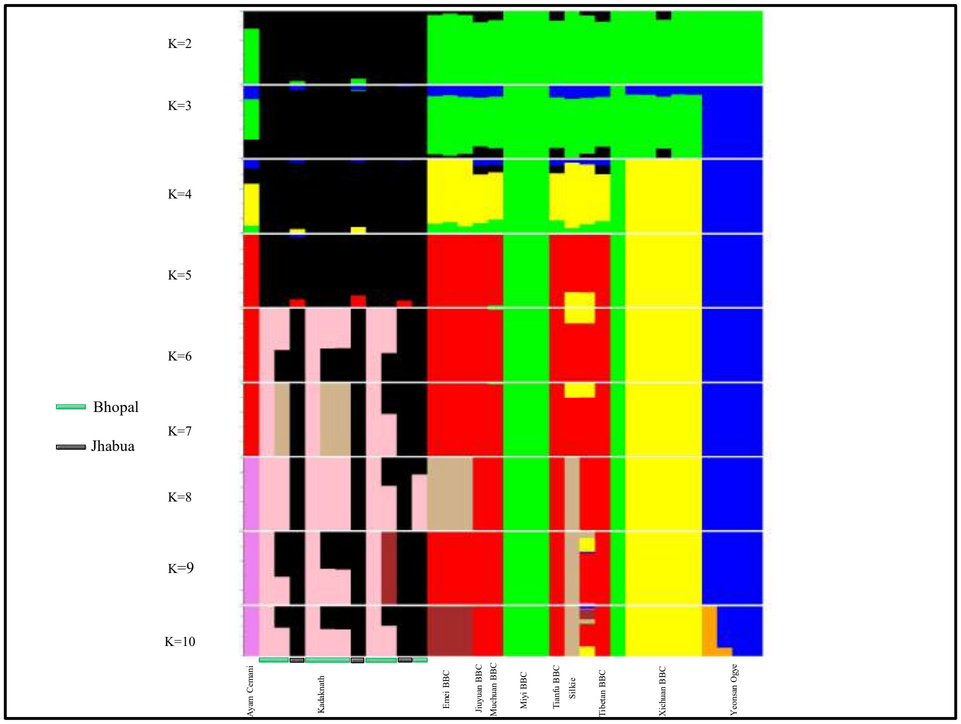


**Fig. S12.** Genome-wide admixture plot of 34 black-bone individuals using NGSadmix from K=2 to K=10. Within the Kadaknath, two sub-clusters (Bhopal and Jhabua) are shown in green and black rectangle boxes, respectively.

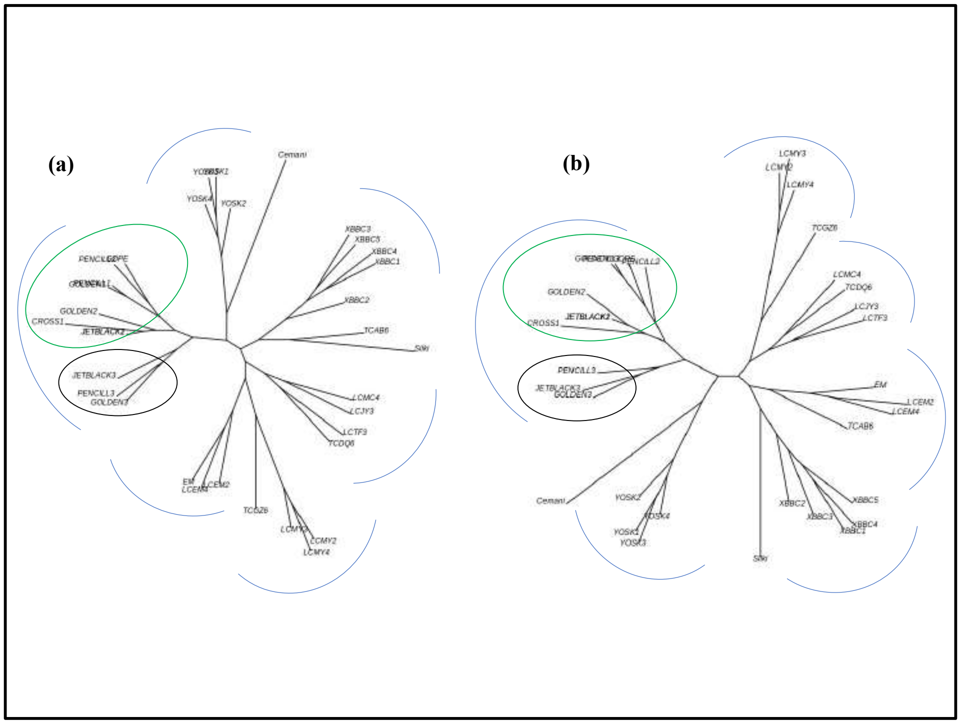


**Fig. S13.** Genome-wide phylogeny of 34 back-bone chicken inferred using the maximum likelihood based SNPhylo method. **(a)** From the bcftools SNP data **(b)** From the freebayes SNP data. Blue curve-shaped lines represent the respective breed individuals. The black color oval shape circle represents KADK individuals from Jhabua, while the green color oval shape circle represents KADK individuals from Bhopal.

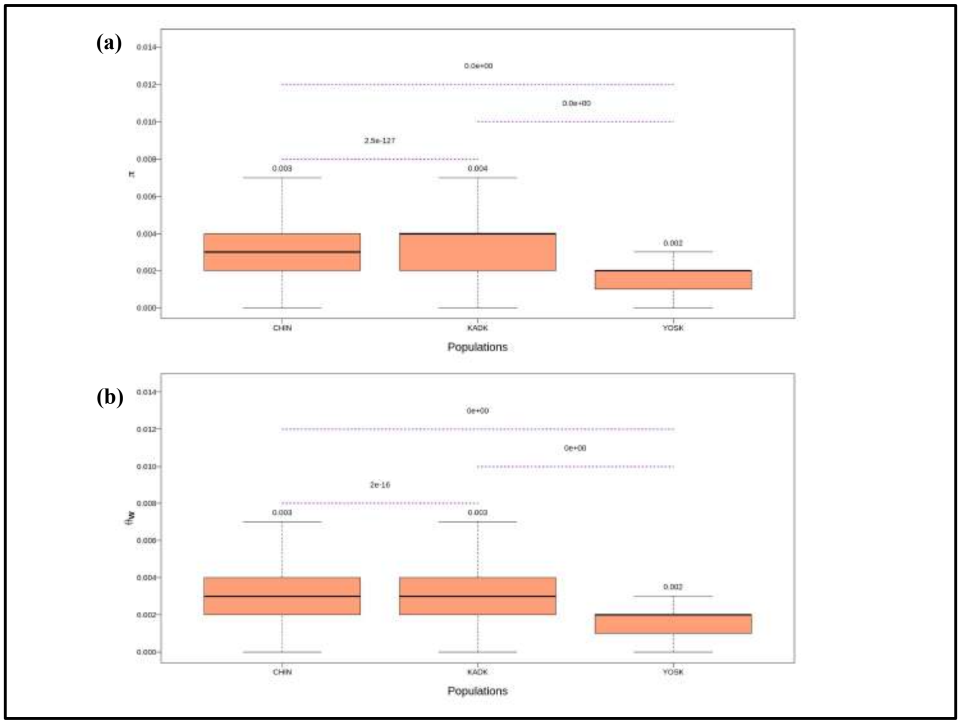


**Fig. S14.** **(a)** Nucleotide diversity (π) and **(b)** genetic diversity (θ) for three populations (CHIN, KADK, YOSK) shown using a Light salmon color boxplot with the mean of each population represented at the top of the box plot. We calculated the pairwise two-tailed Wilcox test representing the purple dotted lines between pairs and the significant p-values mentioned on the top of the dotted line.

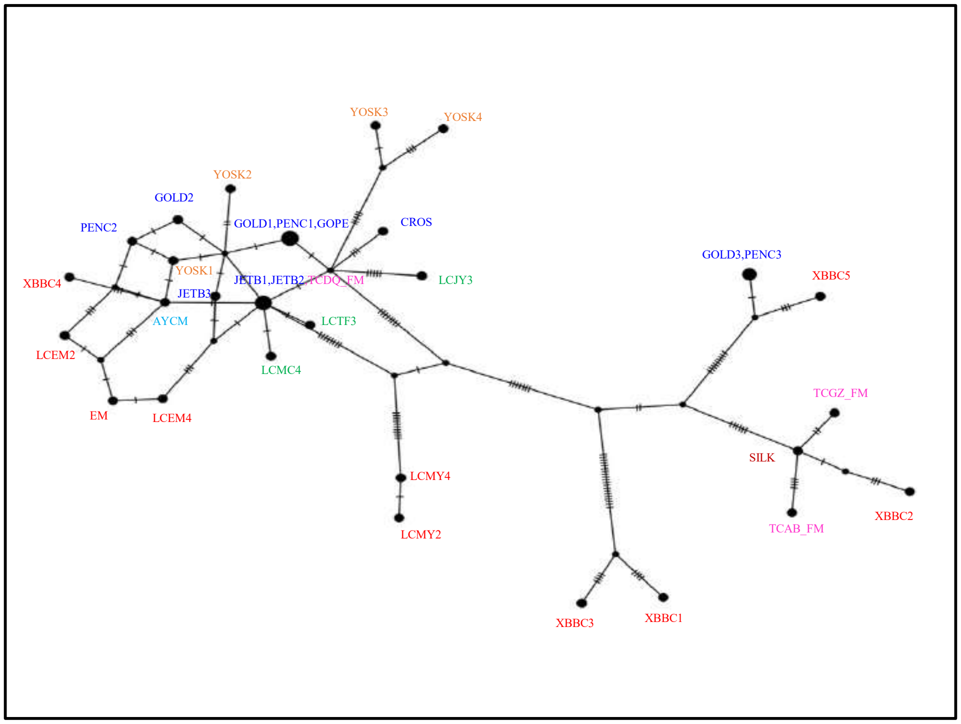


**Fig. S15.** PopART software is used for 33 black-bone chicken individuals for the whole mitochondrial network. Network constructed using the median-joining method.

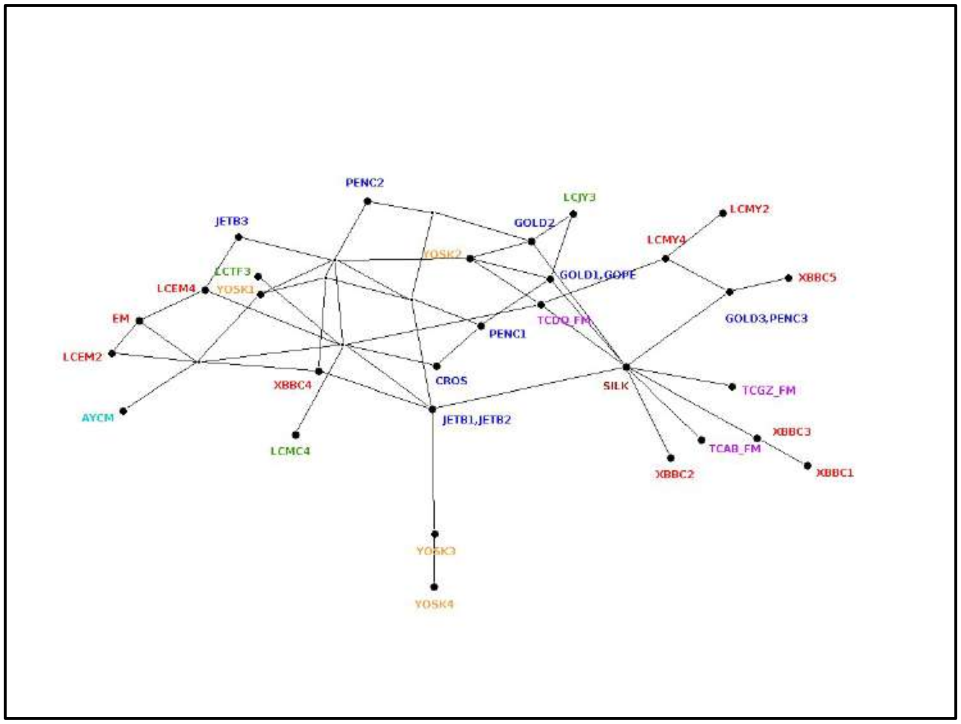


**Fig. S16.** SplitsTree software is used for 33 black-bone chicken individuals for the whole mitochondrial sequence. Network constructed using the median-joining method.

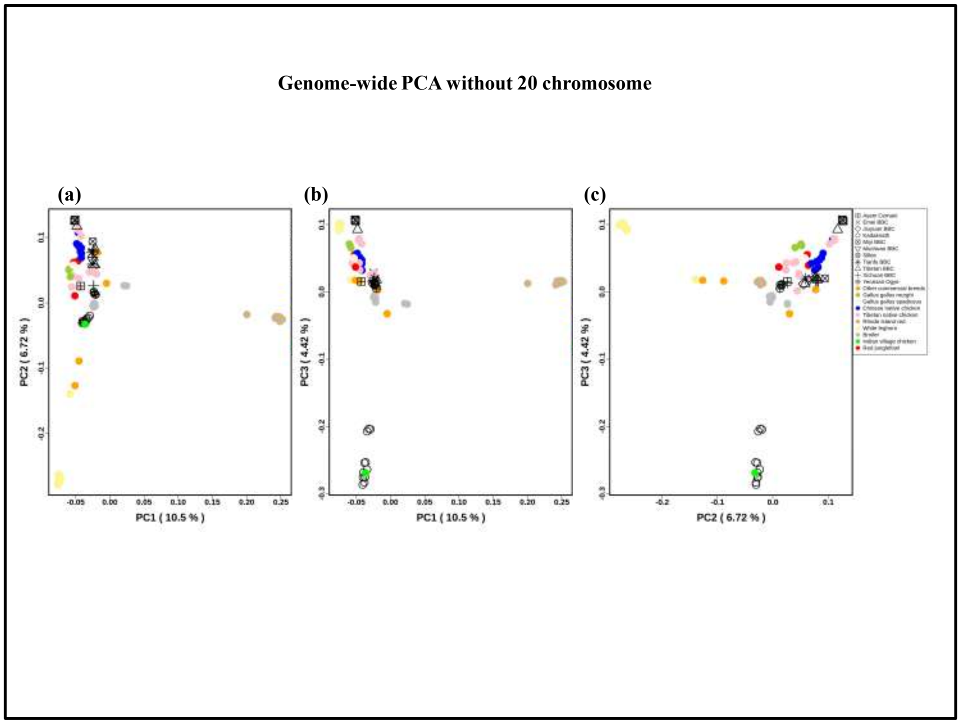


**Fig. S17.** Genome-wide PCA plot generated for 101 chicken individuals: **(a)** PC1 and PC2 explained 10.5% and 6.72 % variance, respectively. **(b)** PC1 and PC3 explained 10.5% and 4.42% variance, respectively. **(c)** PC2 and PC3 explained 6.72% and 4.42% variance, respectively. Non-black-bone breeds are shown by different colors. All black-bone breeds are shown in black color with different shapes for each breed

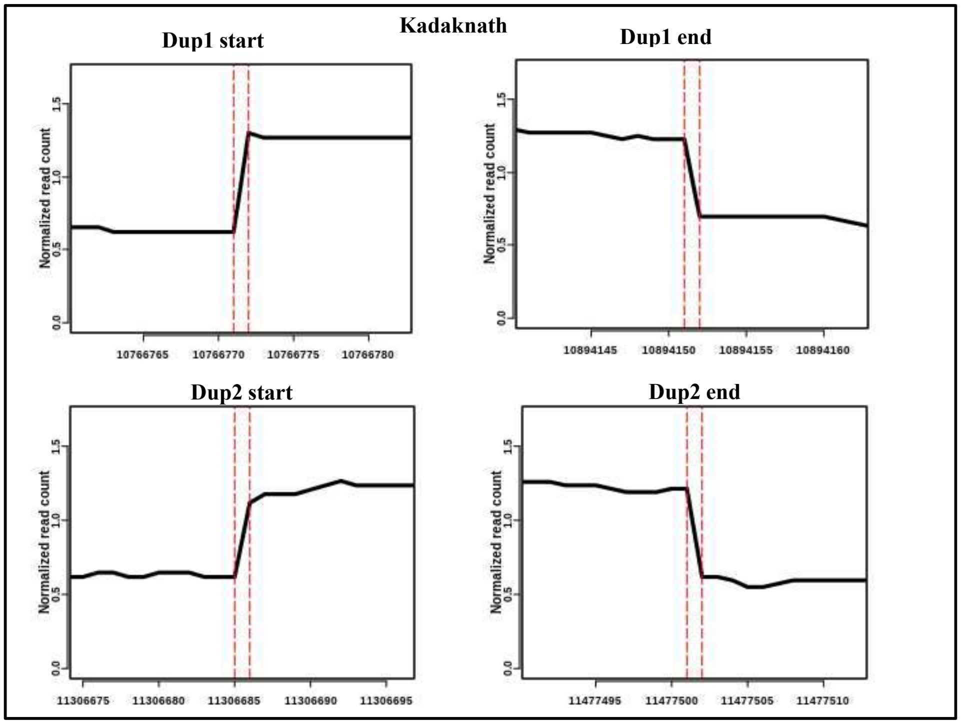


**Fig S18:** Normalized read coverage at Dup1 and Dup2 region: Normalized read coverage at the base-pair level junction in Kadaknath (ERR9560156) for Dup1 start, Dup1 end, Dup2 start, and Dup2 end shown in the figure. The vertical red dotted line represents the exact position of the start and end of Dup1 and Dup2.

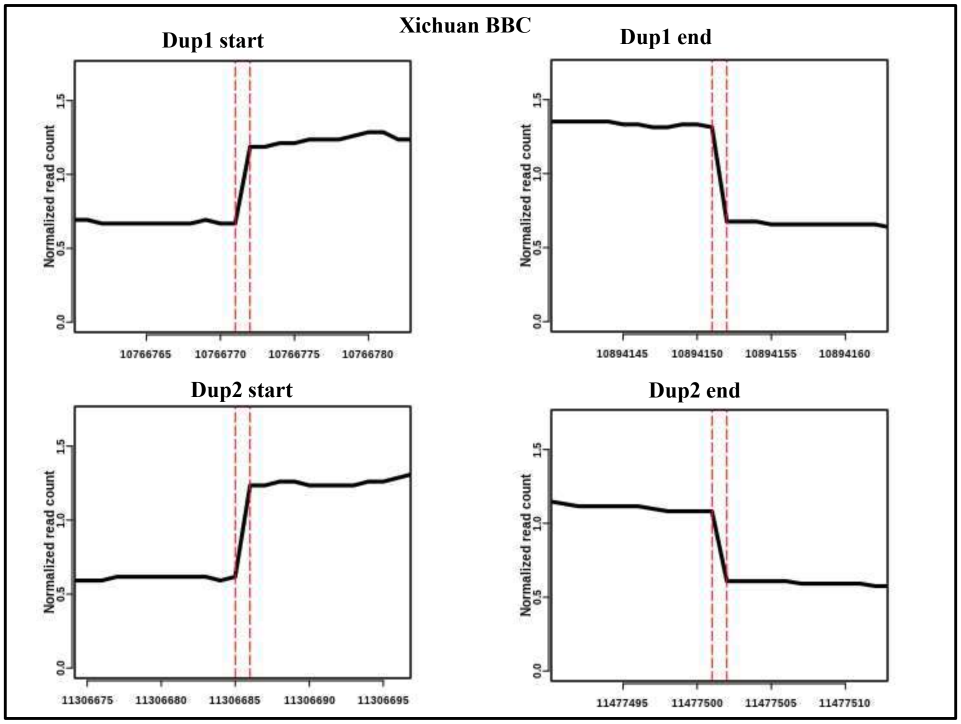


**Fig S19:** Normalized read coverage at Dup1 and Dup2 region: Normalized read coverage at the base-pair level junction in Xichuan BBC (SRR12103812) for Dup1 start, Dup1 end, Dup2 start, and Dup2 end shown in the figure. The vertical red dotted line represents the exact position of the start and end of Dup1 and Dup2.

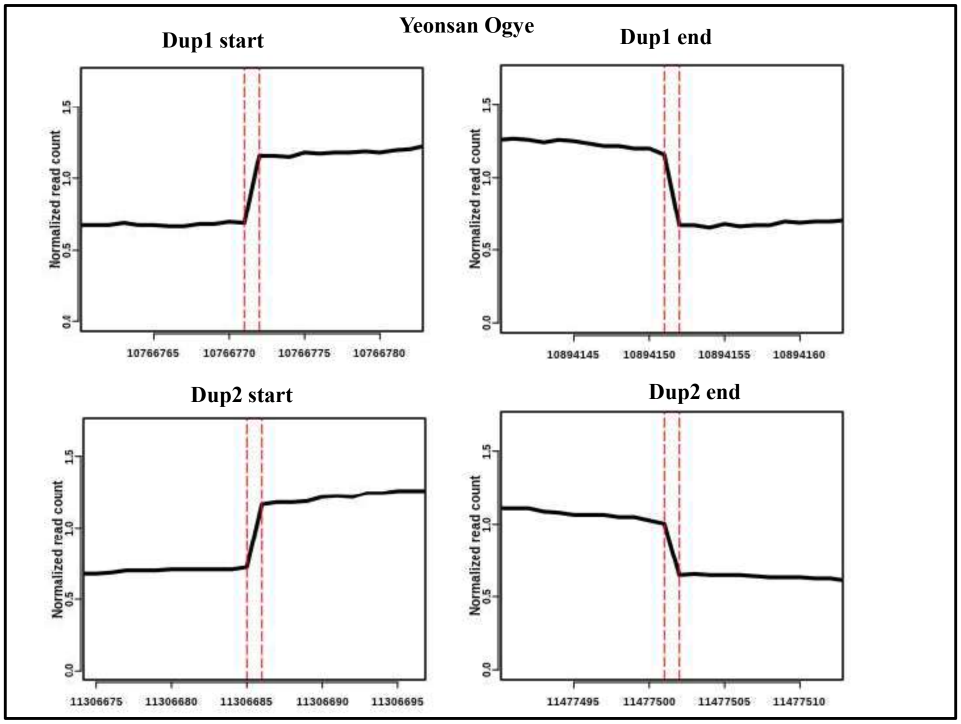


**Fig S20:** Normalized read coverage at Dup1 and Dup2 region: Normalized read coverage at the base-pair level junction in Yeonsan Ogye (SRR6189094, SRR6189095, SRR6189096, SRR6189098, SRR6189082, SRR6189084 and SRR6189087) for Dup1 start, Dup1 end, Dup2 start, and Dup2 end shown in the figure. The vertical red dotted line represents the exact position of the start and end of Dup1 and Dup2.

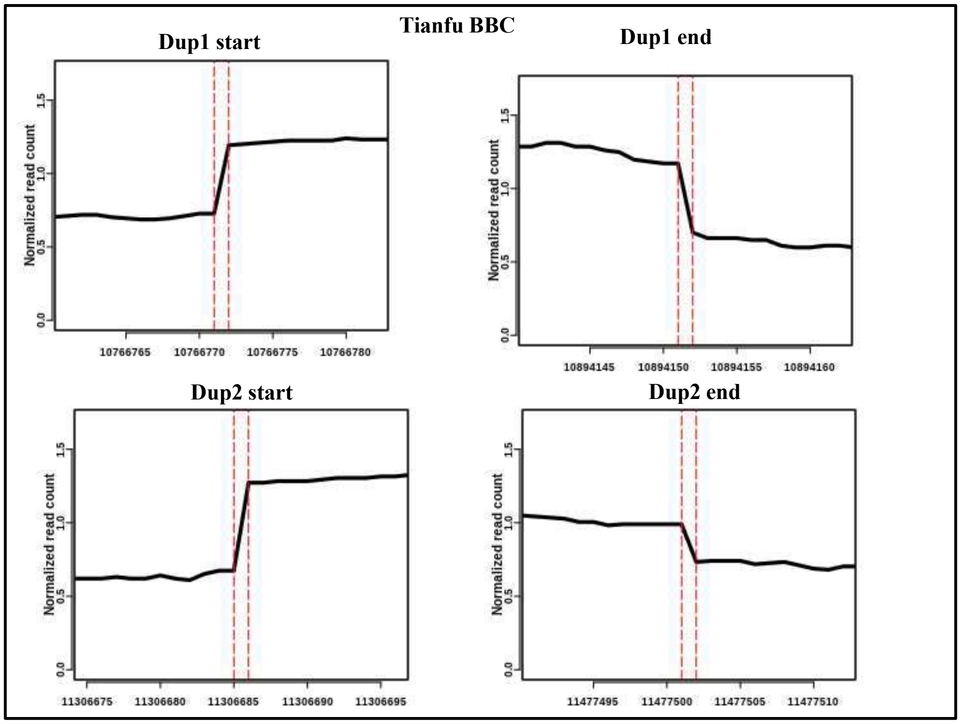


**Fig S21:** Normalized read coverage at Dup1 and Dup2 region: Normalized read coverage at the base-pair level junction in Tianfu BBC (SRR3041425) for Dup1 start, Dup1 end, Dup2 start, and Dup2 end shown in the figure. The vertical red dotted line represents the exact position of the start and end of Dup1 and Dup2.

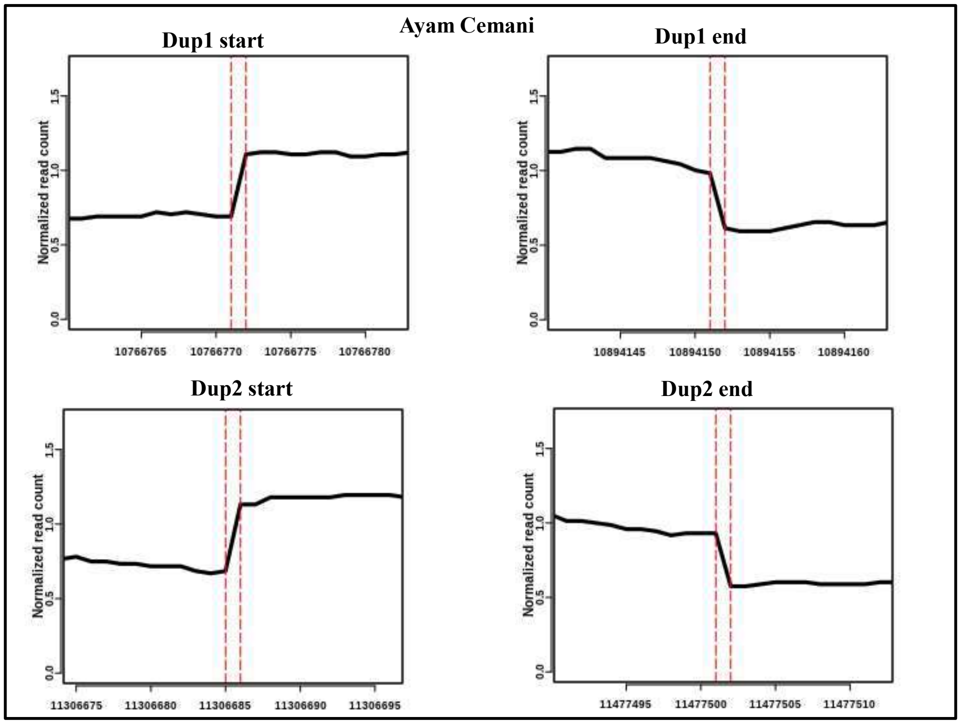


**Fig S22:** Normalized read coverage at Dup1 and Dup2 region: Normalized read coverage at the base-pair level junction in Ayam Cemani (SRR17916231) for **(A)** Dup1 start, Dup1 end, Dup2 start, and Dup2 end shown in the figure. The vertical red dotted line represents the exact position of the start and end of Dup1 and Dup2.

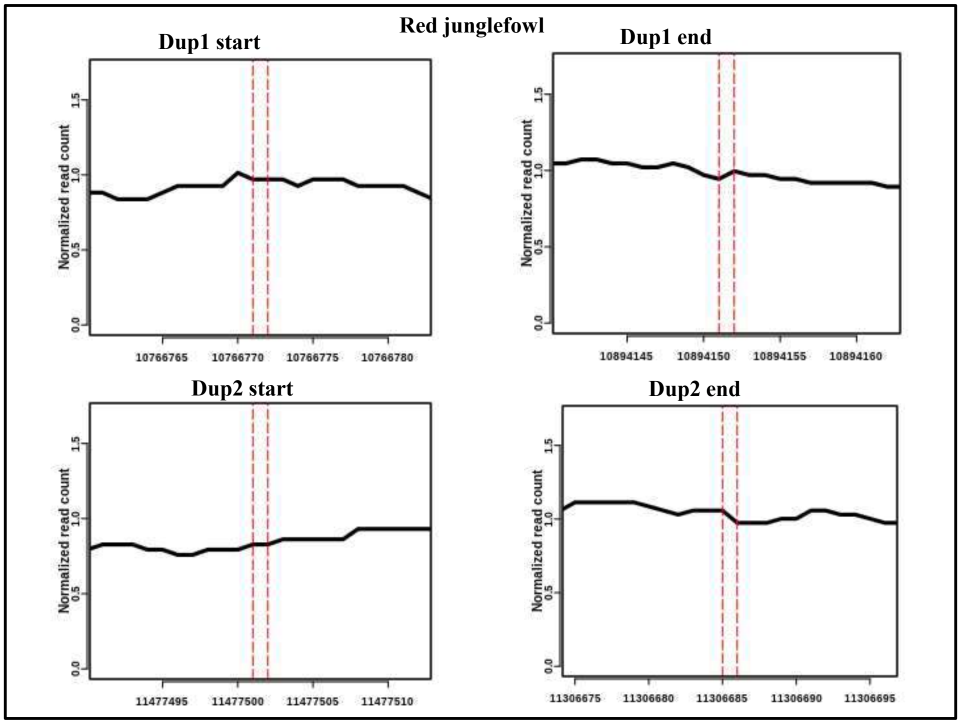


**Fig S23:** Normalized read coverage at Dup1 and Dup2 region: Normalized read coverage at the base-pair level junction in Red junglefowl (SRR3954707) for Dup1 start, Dup1 end, Dup2 start, and Dup2 end shown in the figure. The vertical red dotted line represents the exact position of the start and end of Dup1 and Dup2.

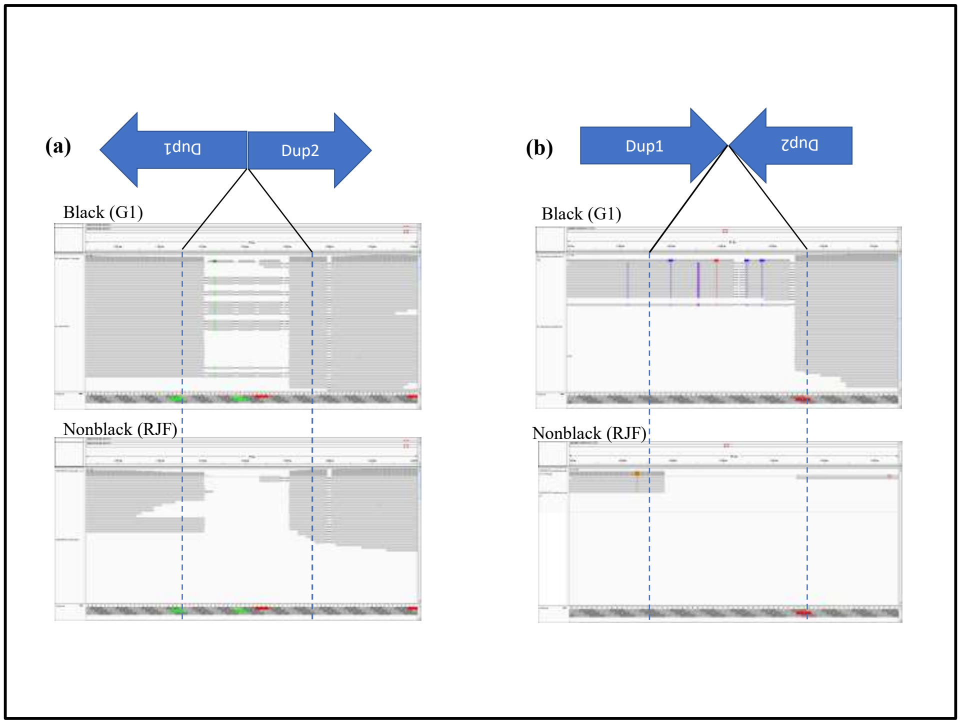


**Fig. S24**. Two rearranged junctions in black chicken: **(a)** inverted_Dup1 Dup2 junction where the blue arrows show the direction of rearrangement, the Dup1 is inverted and Dup2 without inversion. In this region, we mentioned the black and nonblack chicken. On the top panel, a black Kadaknath (Golden1) IGV screenshot of rearranged junction supported reads, while the bottom nonblack (Red junglefowl) chicken without any reads support. The blue dotted line represents the junction region. **(b)** dup1 dup2_inverted in the second junction where the arrows show rearrangement directions, dup1 is without inversion, whereas dup2 is inverted. The top panel defines a black chicken IGV screenshot of reads support to the junction. A bottom panel of nonblack chicken with no read support for the junction in the IGV screenshot.

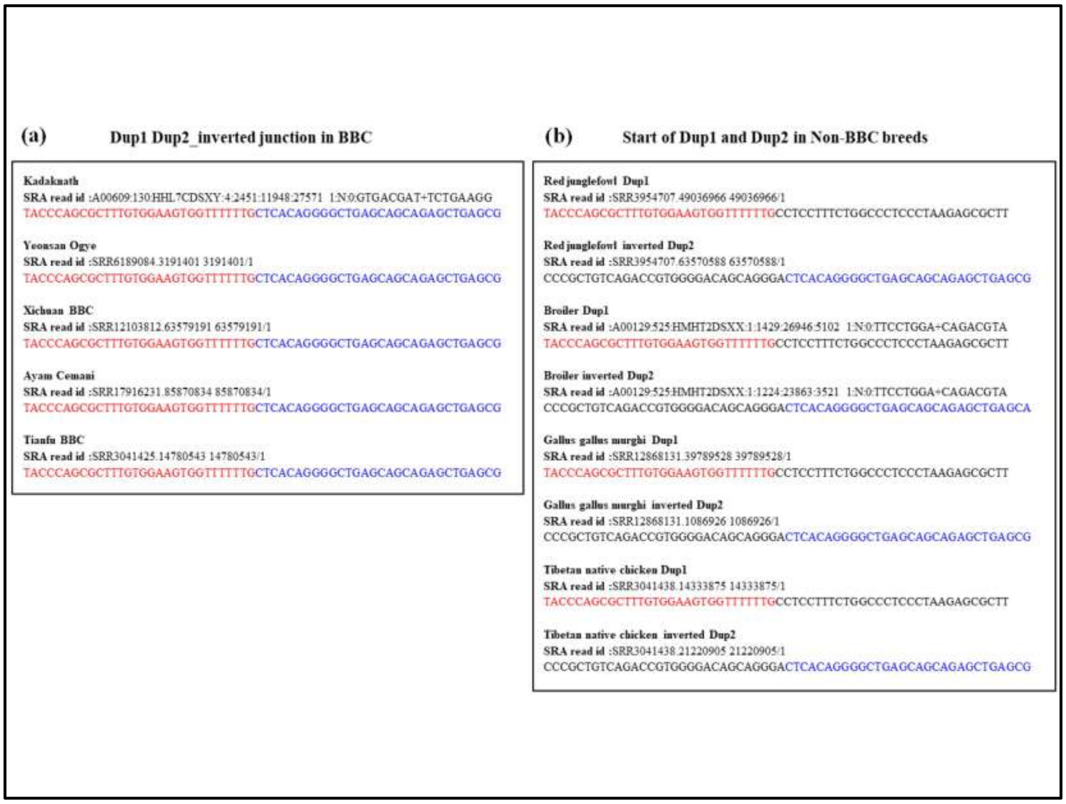


**Fig. S25:** **(a)** Dup1 Dup2 inverted junction in black-bone chicken breeds (b) Dup1 and Dup2 in non-BBC breeds. The reads containing the start of Dup1 and Dup2 have been shown along with their SRA id. The nucleotide sequences highlighted in red and blue represent Dup1 and Dup2, respectively, which is present as continuous sequence BBC breeds. Sequences in black color represent subsequent nucleotides.

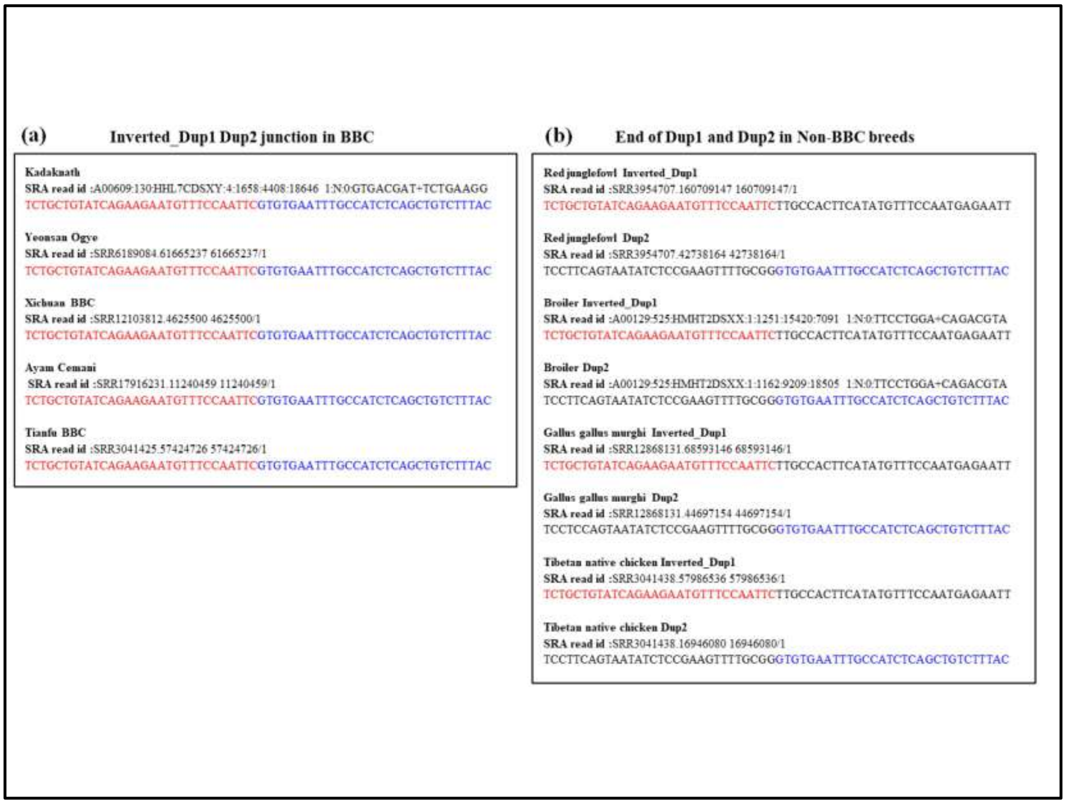


**Fig. S26:** **(a)** Inverted Dup1 Dup2 junction in black-bone chicken breeds **(b)** Dup1 and Dup2 in non-BBC breeds. The reads containing the end of Dup1 and Dup2 have been shown along with their SRA id. The nucleotide sequences highlighted in red and blue represent Dup1 and Dup2, respectively, which is present as continuous sequence BBC breeds. Sequences in black color represent subsequent nucleotides.

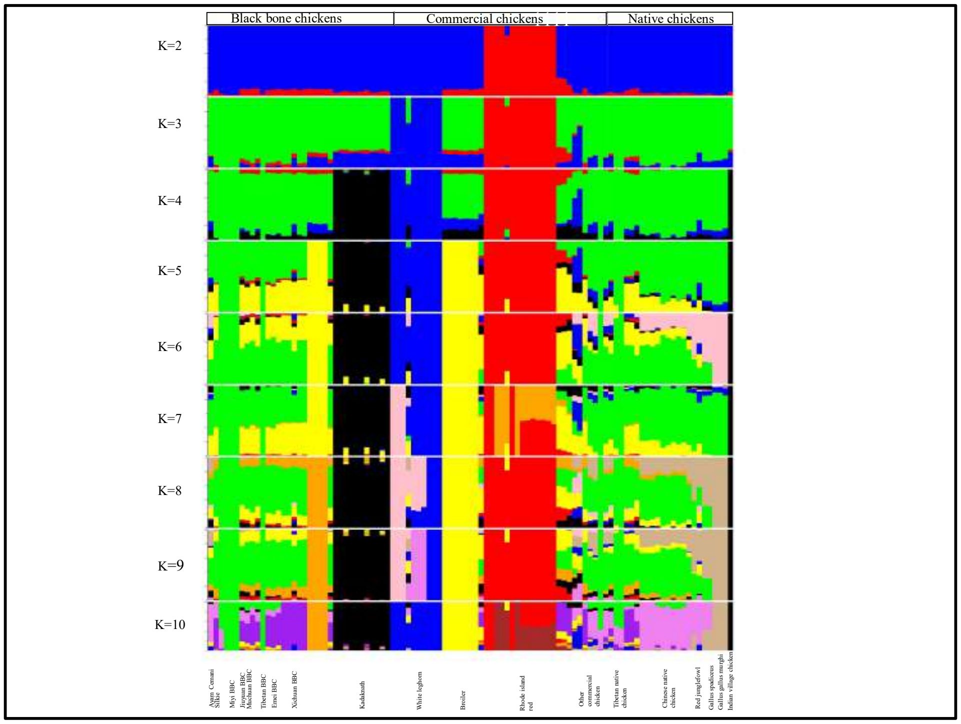


**Fig. S27.** Genome-wide population genetic structure and individual ancestry were estimated using NGSadmix for 101 chickens from different breeds based on best K=2 to K=10

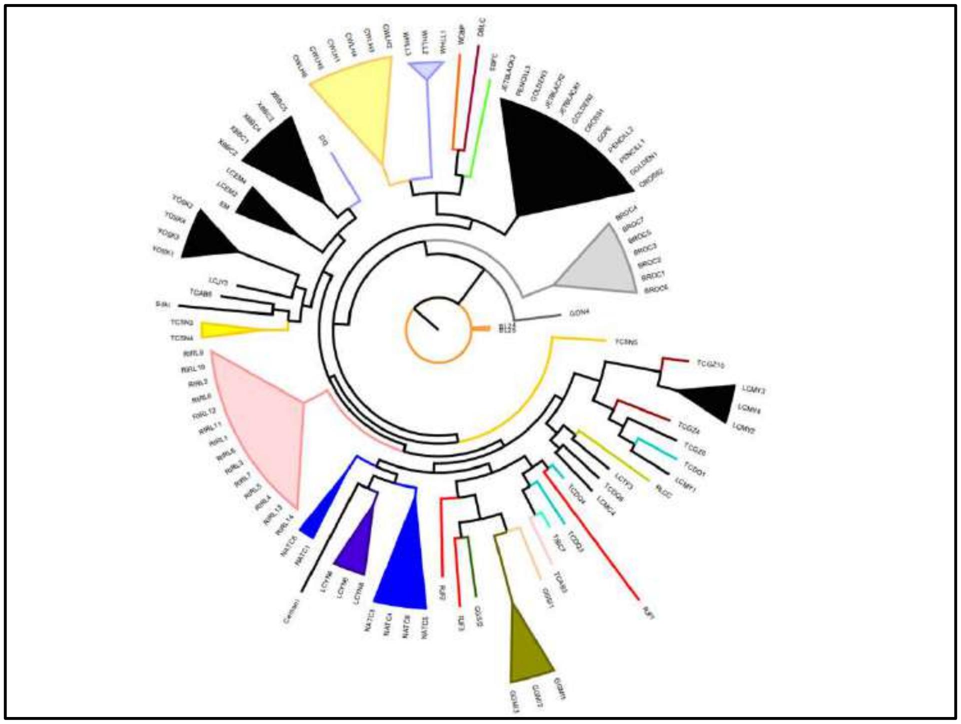

**Fig. S28.** Genome-wide phylogeny of 101 individuals using variant call from bcftools. The phylogeny is generated using SNPhylo and visualized in Figtree.

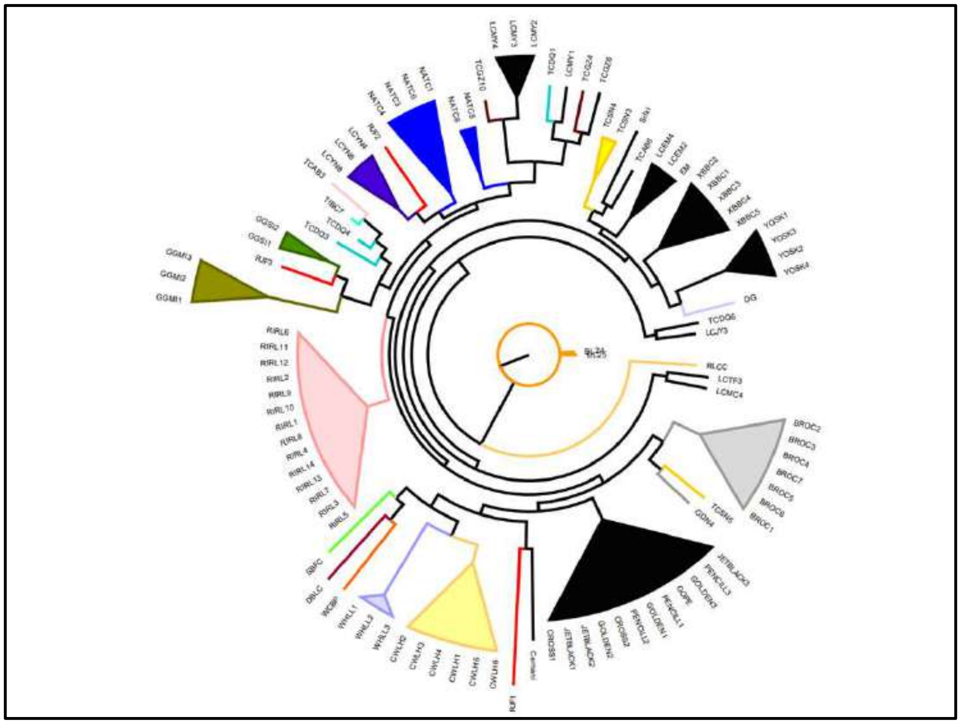

**Fig. S29.** Genome-wide phylogeny of 101 individuals using variant call from freebayes. The phylogeny is generated using SNPhylo and visualized in Figtree.

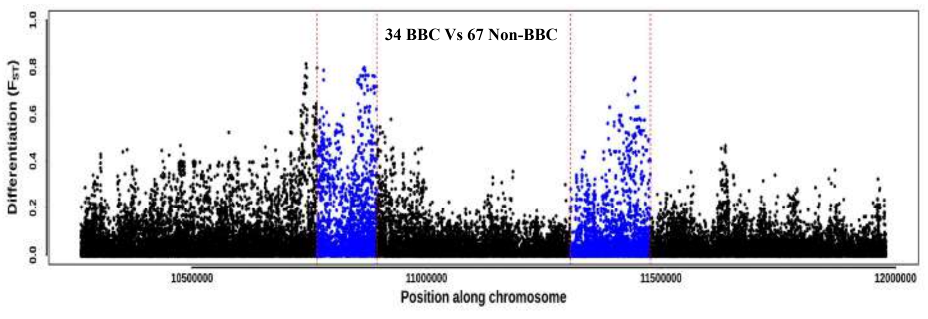


**Fig. S30.** $F_{ST}$ landscape of *Fm* locus between 34 black-bone chicken (BBC) and 67 non-black-bone chickens (non-BBC) is shown in a 1bp windows. Dup1 and Dup2 regions are represented in blue color dots, showing high $F_{ST}$ compared to the nearby regions.

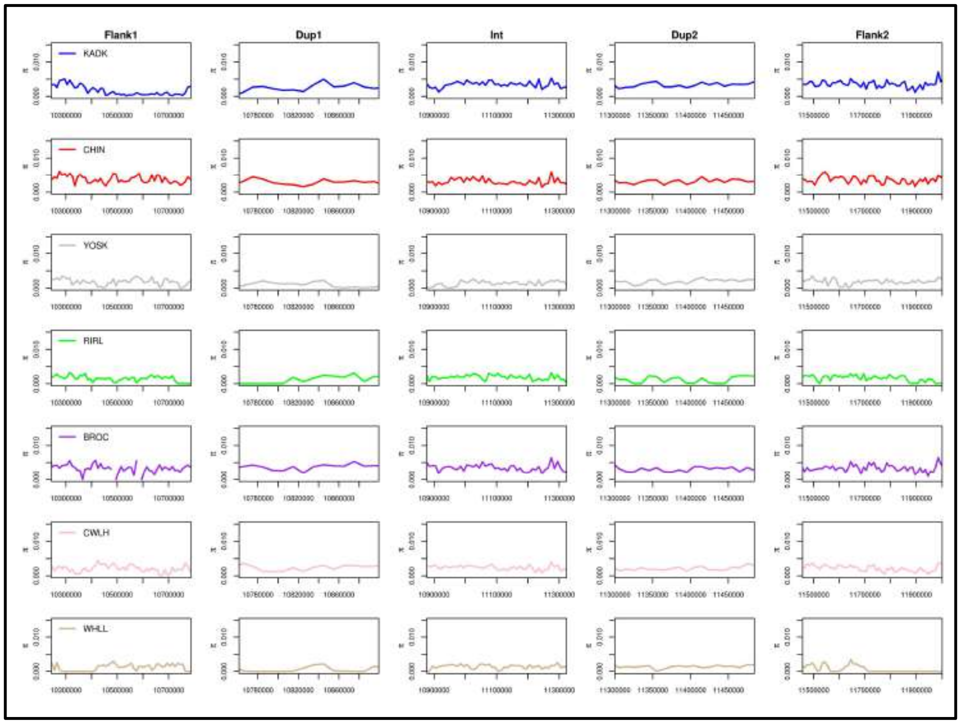


**Fig. S31.** Nucleotide diversity (**π**) at *Fm* locus regions (Flank1, Dup1, Int, Dup2, and Flank2) has been shown for each breed represented in different colors using 10Kb sliding windows.

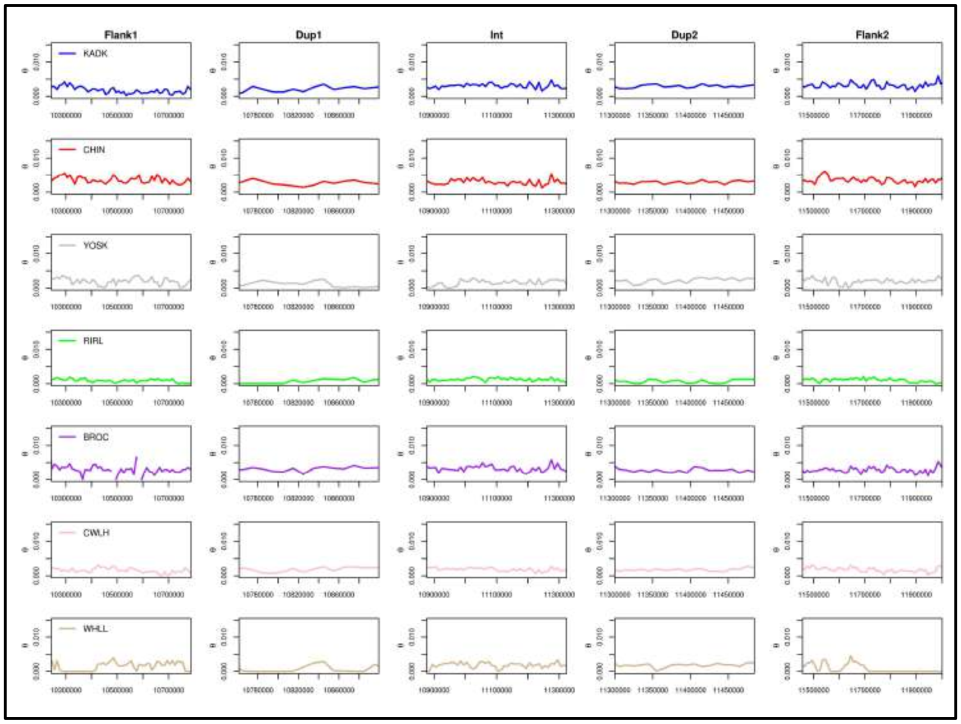


**Fig. S32.** Estimated Genetic diversity (**θ**) at *Fm* locus regions (Flank1, Dup1, Int, Dup2, and Flank2) has been shown for each breed represented in different colors using 10Kb sliding windows.

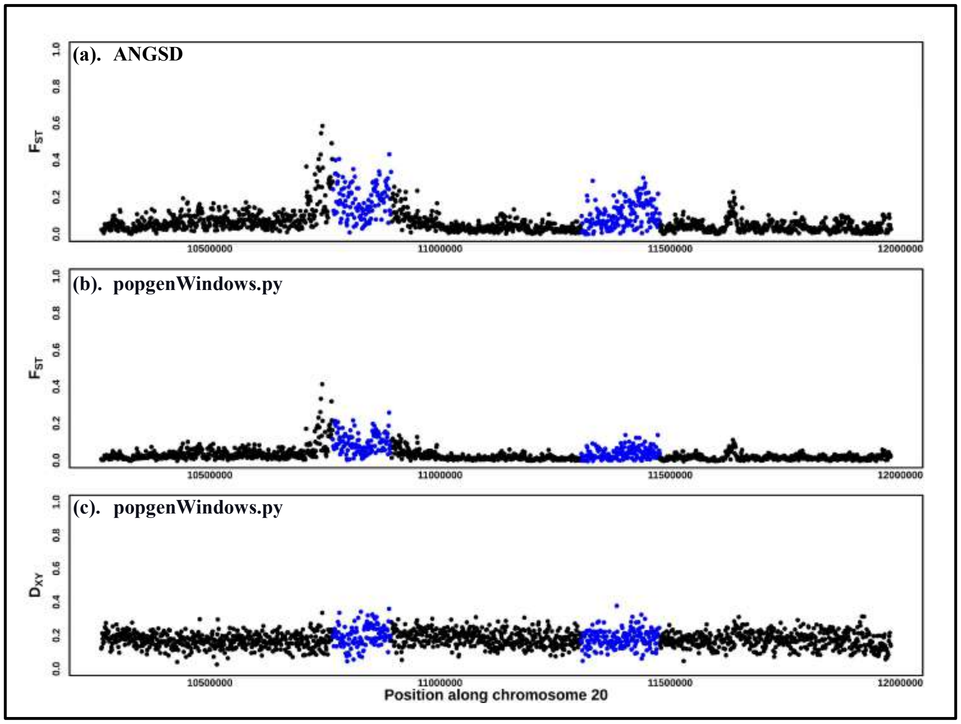


**Fig. S33.** Pairwise $F_{ST}$ estimation at *Fm* locus between 34 black-bone and 67 non-black-bone chickens is shown in a 1Kb window calculated using **(a)** ANGSD, **(b)** popgenWindows.py, and **(c)** Dxy using popgenWindows.py. Dup1 and Dup2 regions are represented in blue color dots, having high $F_{ST}$ compared to the nearby regions.

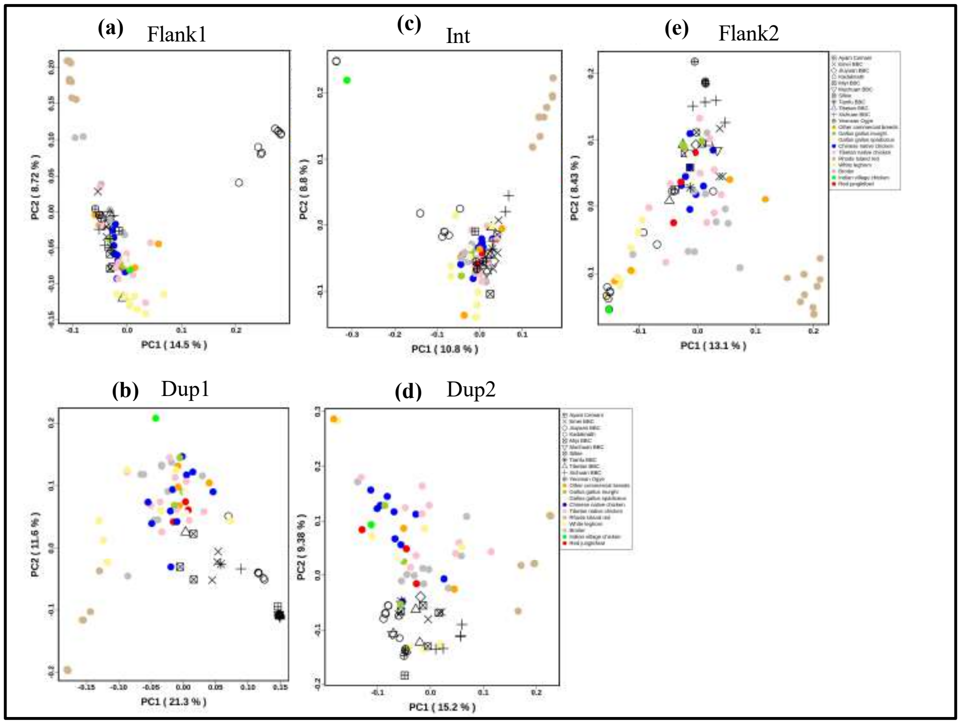


**Fig. S34.** Principal component analysis (PCA) of *Fm* region for 101 chicken individuals with the first two principal groups (PC1 and PC2). **(a)** Flank1 **(b)** Dup1 **(c)** Int **(d)** Dup2 **(e)** Flank2. Non-black-bone breeds are shown in different colors, while all black-bone breeds are shown in black with different shapes for each breed.

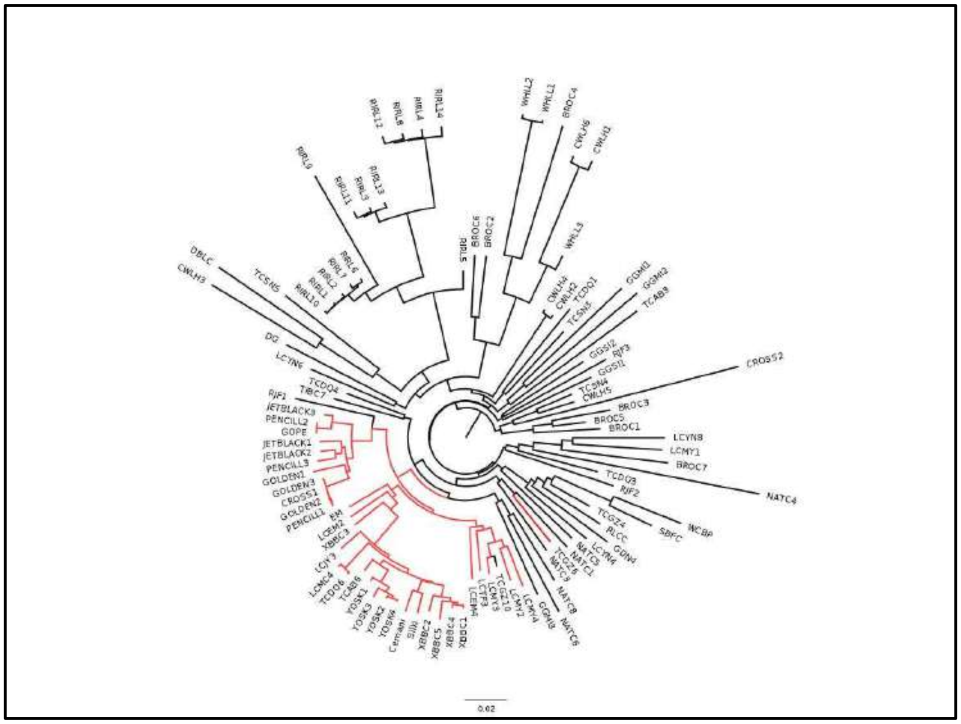

**Fig. S35.** Phylogenetic relationship of Dup1 and Dup2 region for 101 individuals using a variant call from bcftools has been shown. Red color nodes represent the black-bone chickens, and black color nodes represent the non-black-bone chickens.

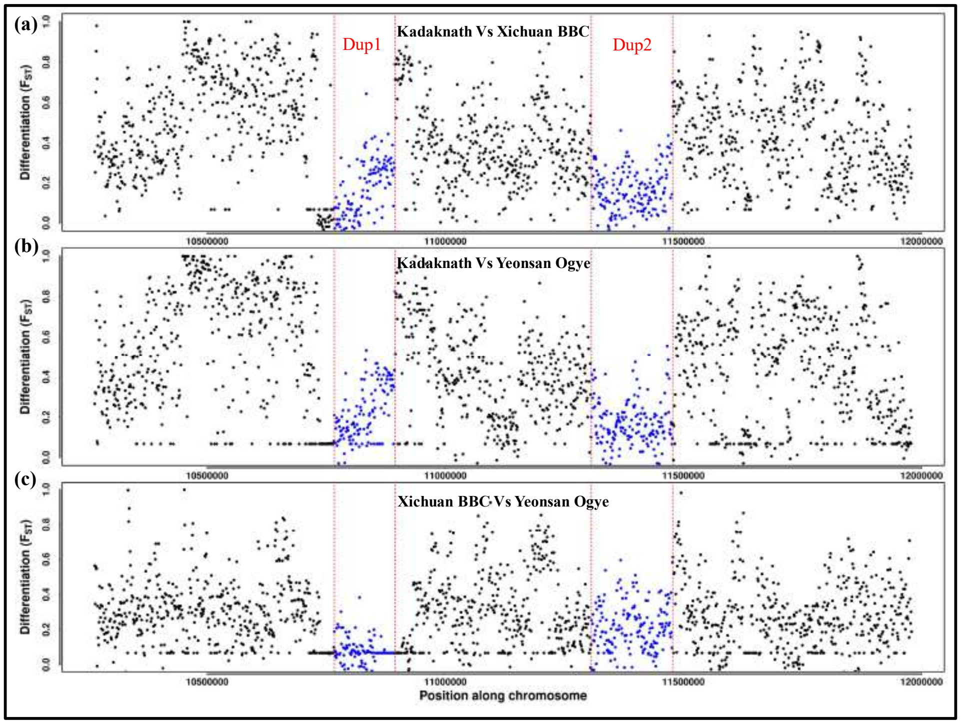


**Fig. S36:** Pairwise $F_{ST}$ comparisons between black-bone populations: $F_{ST}$ landscape of *Fm* locus between **(a)** Kadaknath and Xichuan BBC ,**(b)** Kadaknath BBC and Yeonsan Ogye and **(c)** Xichuan BBC and Yeonsan Ogye is shown in a 1Kb non-overlapping windows. Dup1 and Dup2 regions are represented in blue color dots.

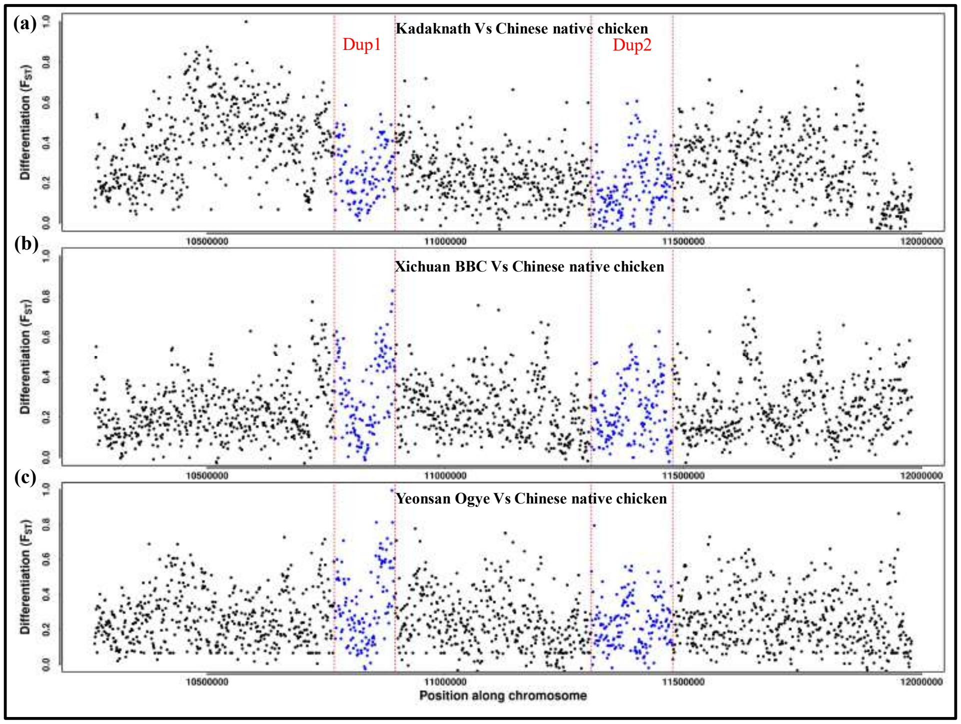


**Fig. S37:** Pairwise $F_{ST}$ comparisons between black-bone and non-black-bone populations: $F_{ST}$ landscape of *Fm* locus between **(a)** Kadaknath and Chinese native chicken,**(b)** Xichuan BBC and Chinese native chicken and **(c)** Yeonsan Ogye and Chinese native chicken is shown in a 1Kb non-overlapping windows. Dup1 and Dup2 regions are represented in blue color dots.

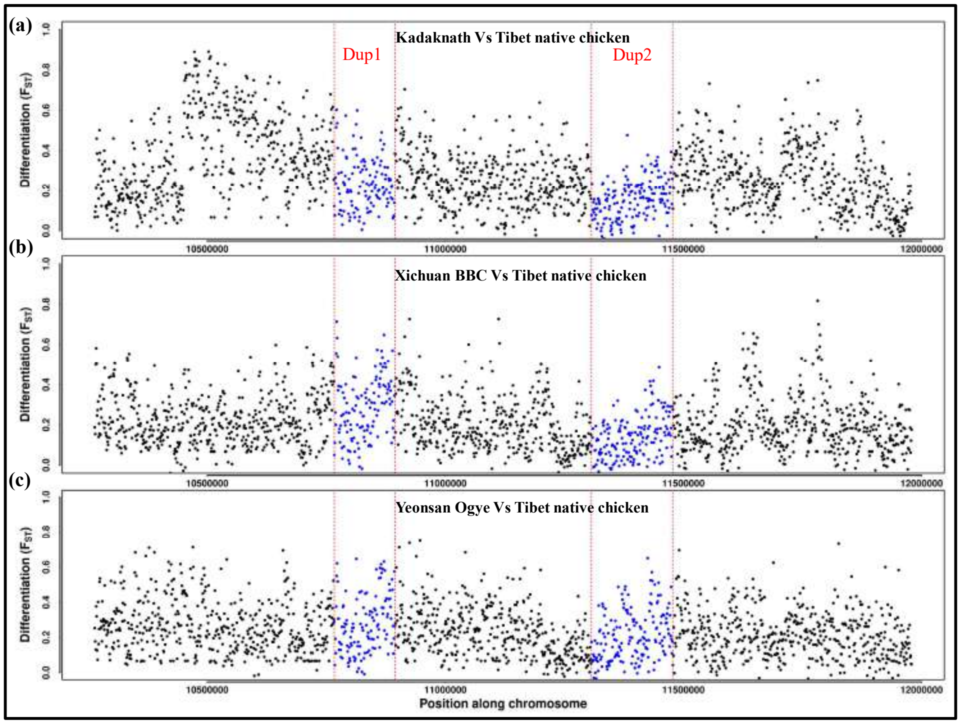


**Fig. S38:** Pairwise $F_{ST}$ comparisons between black-bone and non-black-bone population: $F_{ST}$ landscape of *Fm* locus between **(a)** Kadaknath and Tibet native chicken,**(b)** Xichuan BBC and Tibet native chicken and **(c)** Yeonsan Ogye and Tibet native chicken is shown in a 1Kb non-overlapping windows. Dup1 and Dup2 regions are represented in blue color dots.

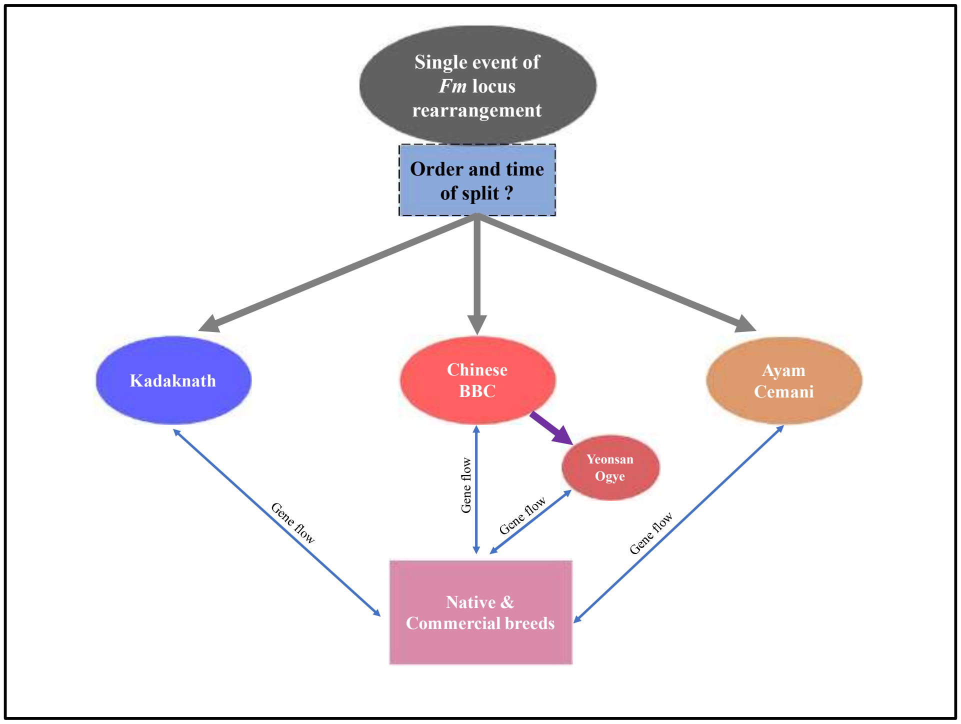


**Fig. S39:** The origin of black-bone phenotype(*Fm* locus rearrangement) and distribution of BBC breeds: The common origin of black-bone chicken through a single event at *Fm* locus rearrangement has been shown. The grey color arrows represent the distribution of different BBC breeds from a common ancestor. The purple color arrow represents the origin of Yeonsan Ogye from Chinese black-bone breeds. The two-sided blue color arrow represents the gene flow between different BBC breeds and various native and commercial breeds.

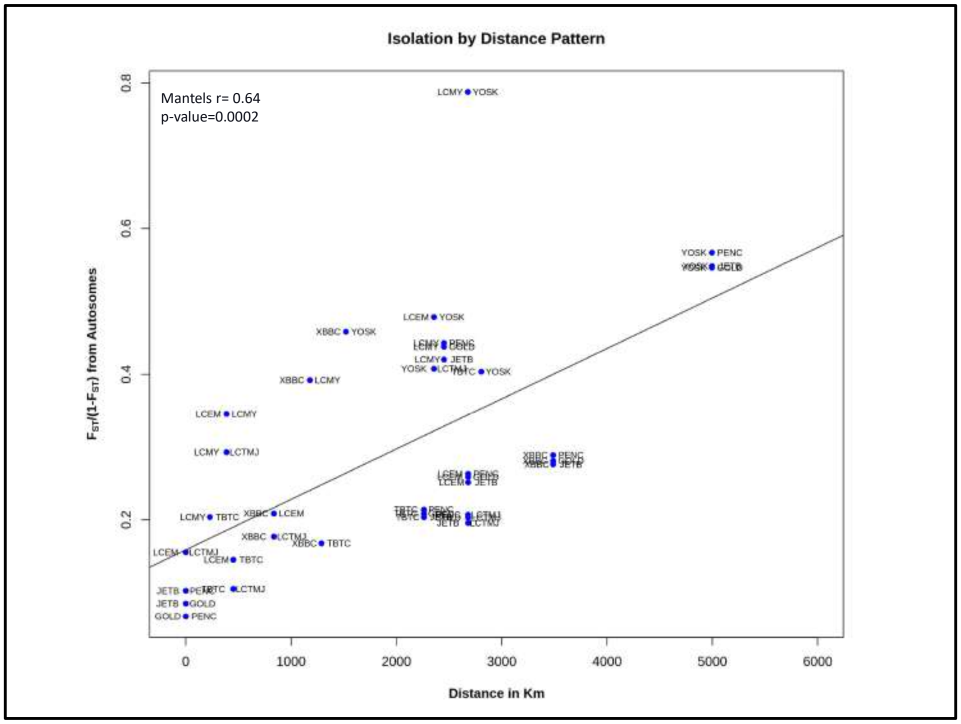


**Fig. S40.** Isolation by distance(IBD) pattern between 9 black-bone chicken populations: Pairwise mean $F_{ST}$ comparison between 9 black-bone chicken populations (JETB, PENC, GOLD, XBBC, LCMY, LCEM, TBTC, YOSK, and LCTMJ) using ANGSD is shown at Y-axis, while the geographical distance in km is shown at X-axis.

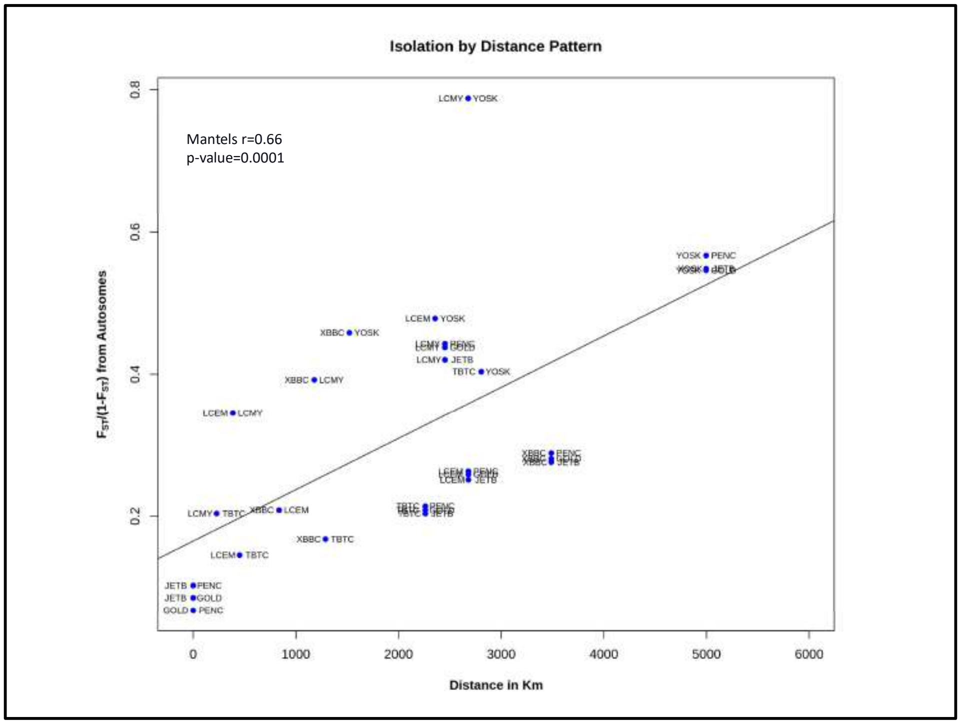


**Fig. S41.** Isolation by distance(IBD) pattern between 8 black-bone chicken populations: Pairwise mean $F_{ST}$ comparison between 8 black-bone chicken populations (JETB, PENC, GOLD, XBBC, LCMY, LCEM, TBTC, and YOSK) using ANGSD is shown at Y-axis, while the geographical distance in km is shown at X-axis.

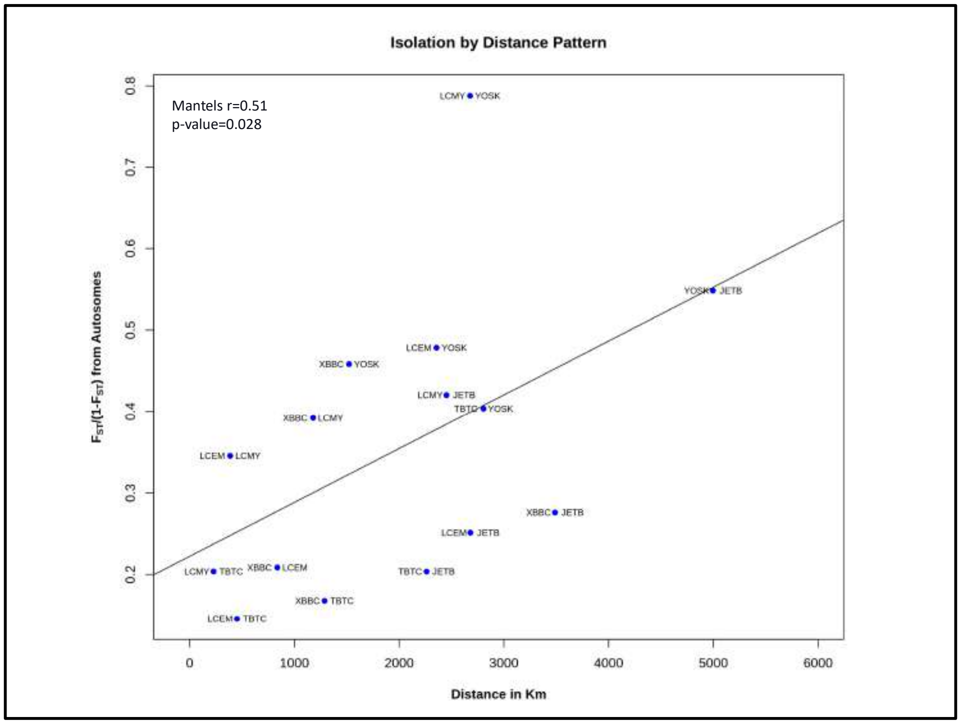


**Fig. S42.** Isolation by distance(IBD) pattern between 6 black-bone chicken populations: Pairwise mean $F_{ST}$ comparison between 6 black bone-chicken populations (JETB, XBBC, LCMY, LCEM, TBTC, and YOSK) using ANGSD is shown at Y-axis, while the geographical distance in km is shown at X-axis.

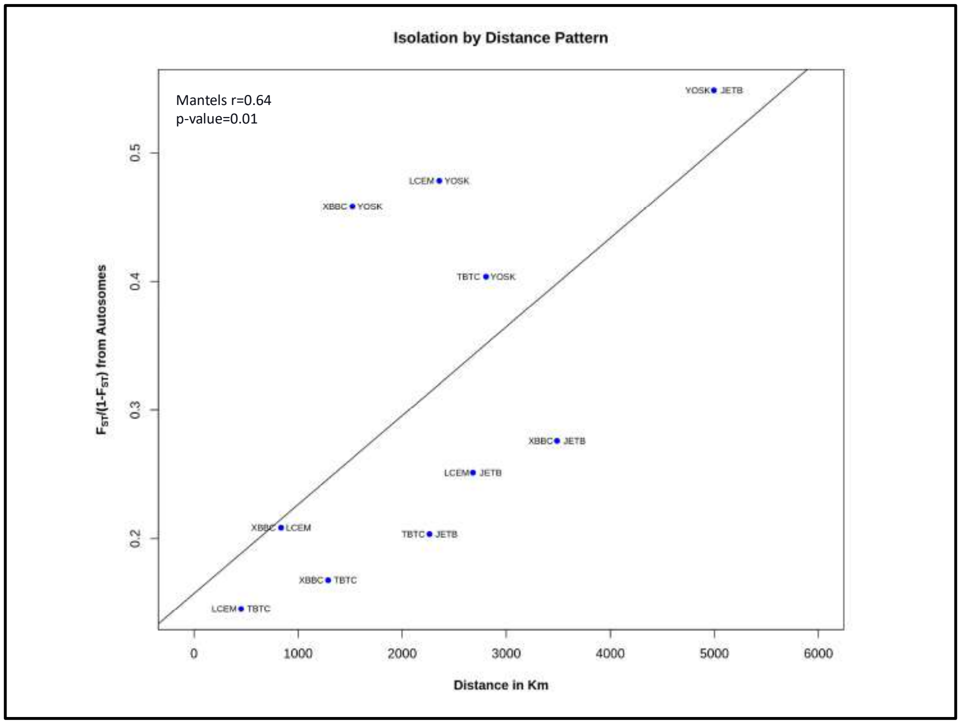


**Fig. S43.** Isolation by distance(IBD) pattern between 5 black-bone chicken populations: Pairwise mean $F_{ST}$ comparison between 5 black-bone chicken populations (JETB, XBBC, LCEM, TBTC, and YOSK) using ANGSD is shown at Y-axis, while geographical distance in km is shown at X-axis.

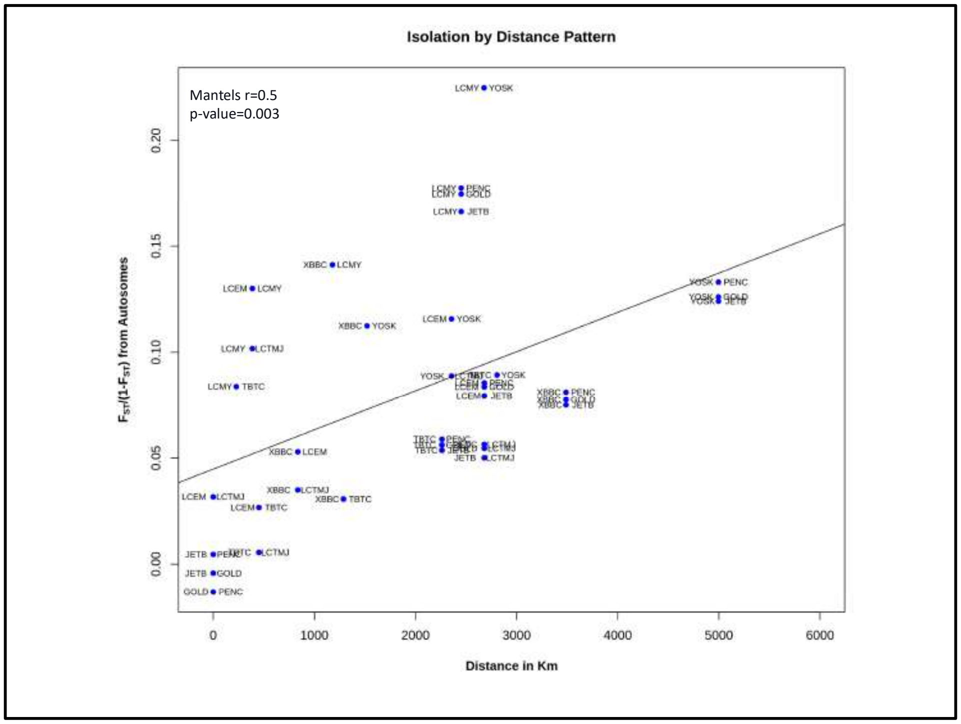


**Fig. S44.** Isolation by distance(IBD) pattern between 9 black-bone chicken populations : Pairwise mean $F_{ST}$ comparison between 9 black bone-chicken populations (JETB, PENC, GOLD, XBBC, LCMY, LCEM, TBTC, LCTMJ, and YOSK) using popgenWindows.py is shown at Y-axis, while the geographical distance in km is shown at X-axis.

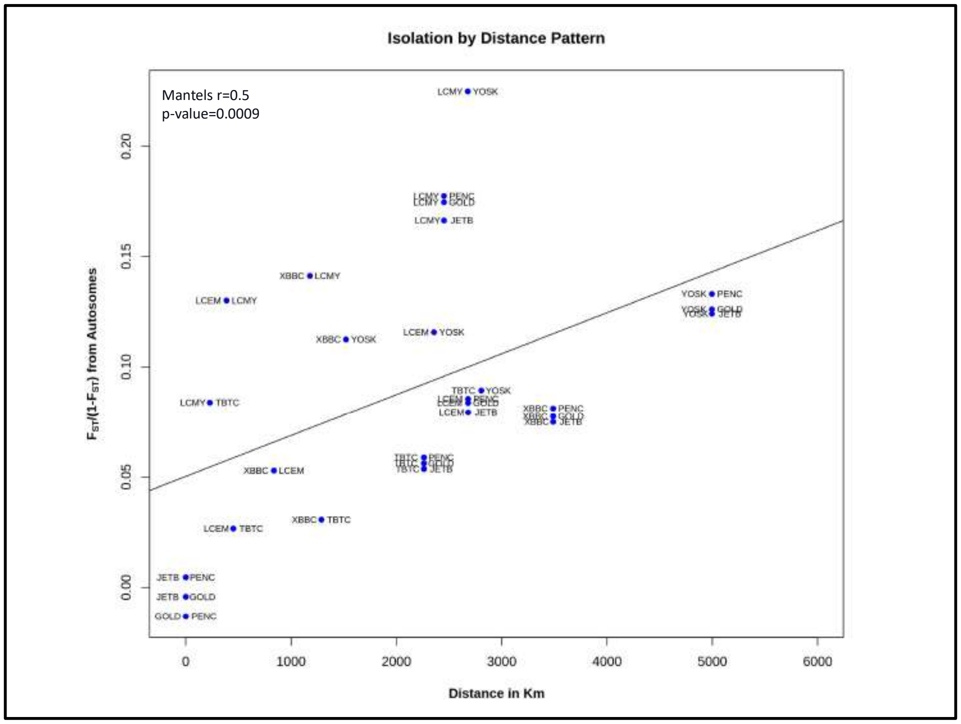


**Fig. S45.** Isolation by distance(IBD) pattern between 8 black-bone chicken populations : Pairwise mean $F_{ST}$ comparison between 8 black-bone chicken populations (JETB, PENC, GOLD, XBBC, LCMY, LCEM, TBTC, and YOSK) using popgenWindows.py is shown at Y axis, while the geographical distance in km is shown at X axis.

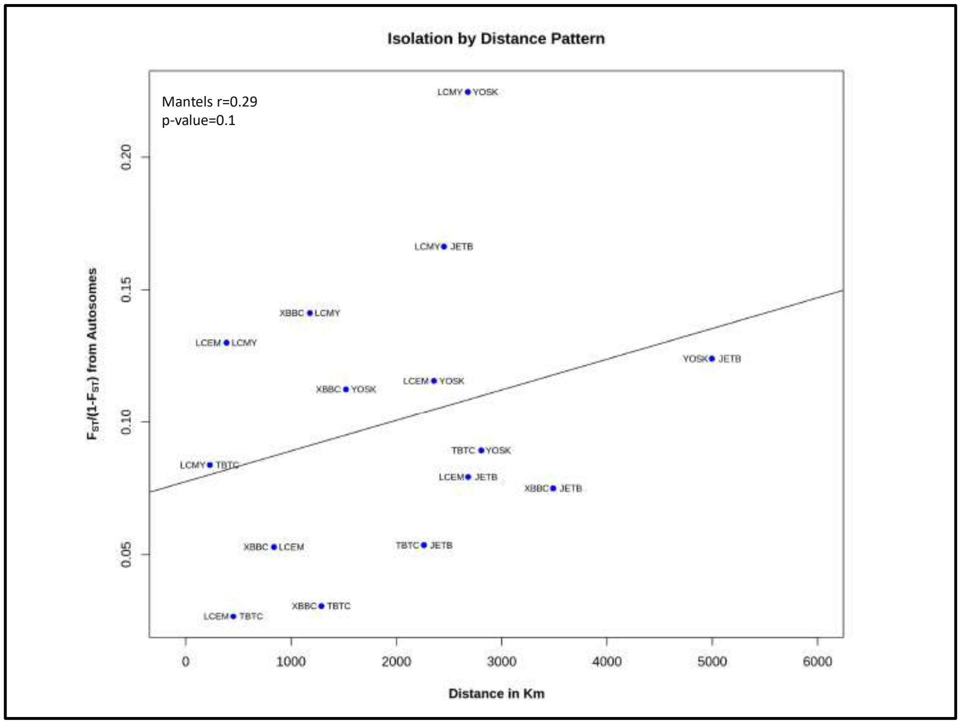


**Fig. S46.** Isolation by distance(IBD) pattern between 6 black-bone chicken populations : Pairwise mean $F_{ST}$ comparison between 6 black-bone chicken populations (JETB, XBBC, LCMY, LCEM, TBTC, and YOSK) using popgenWindows.py is shown at Y-axis, while the geographical distance in km is shown at X-axis.

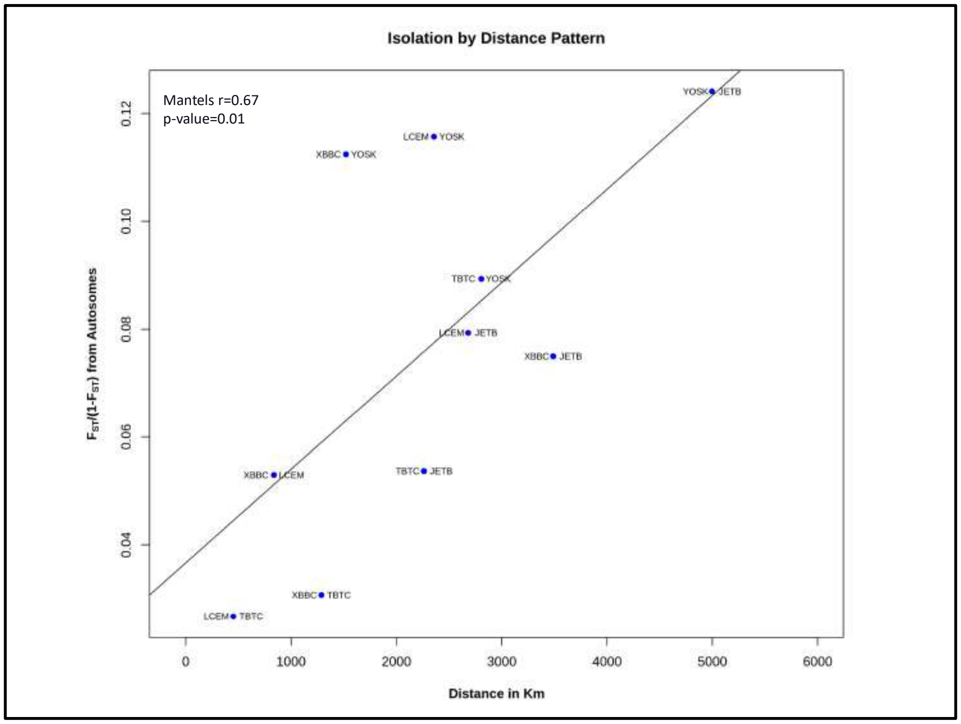


**Fig. S47.** Isolation by distance(IBD) pattern between 5 black-bone chicken populations: Pairwise mean $F_{ST}$ comparison between 5 black-bone chicken populations (JETB, XBBC, LCEM, TBTC, and YOSK) using popgenWindows.py is shown at Y-axis, while the geographical distance in km is shown at X-axis.

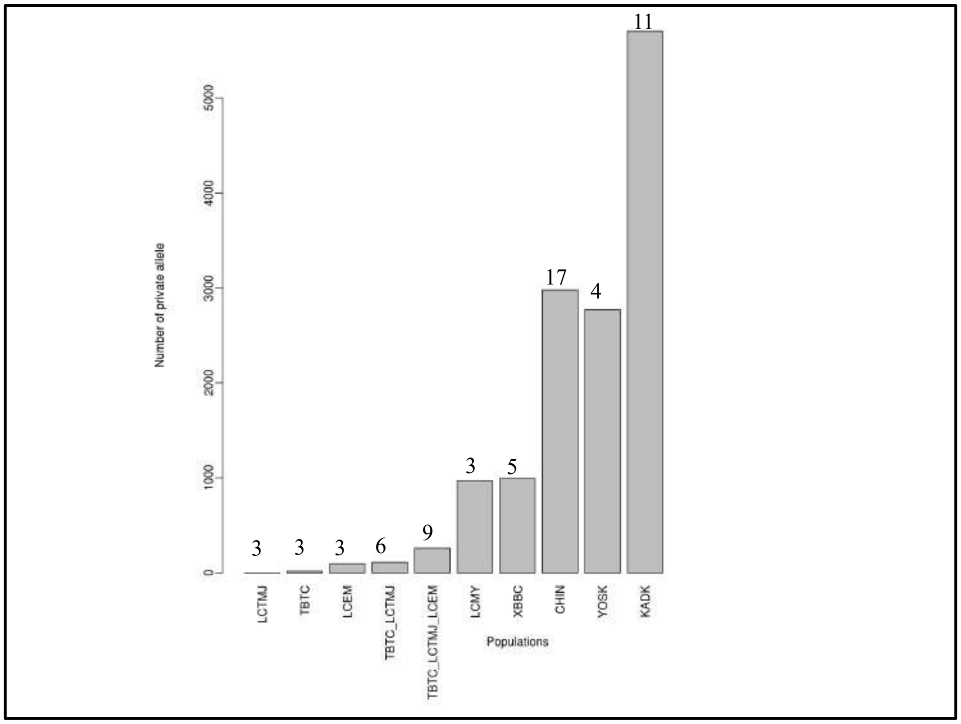


**Fig. S48.** Population-specific private allele count showed on Y-axis and X-axis represents the populations. The numbers of individuals in each population is mentioned at the top of each bar.

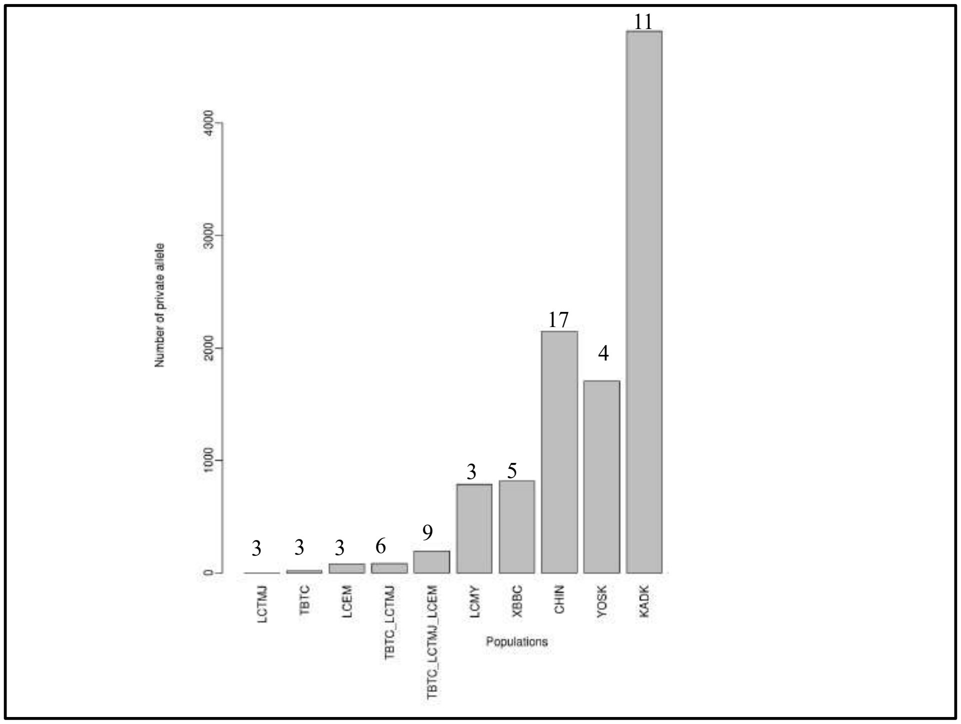


**Fig. S49.** Population-specific private allele count shown on Y-axis and X-axis represents the populations. The numbers of individuals in each population is mentioned at the top of each bar. The counts of population-specific private alleles are obtained after removing the common alleles from dbSNP.

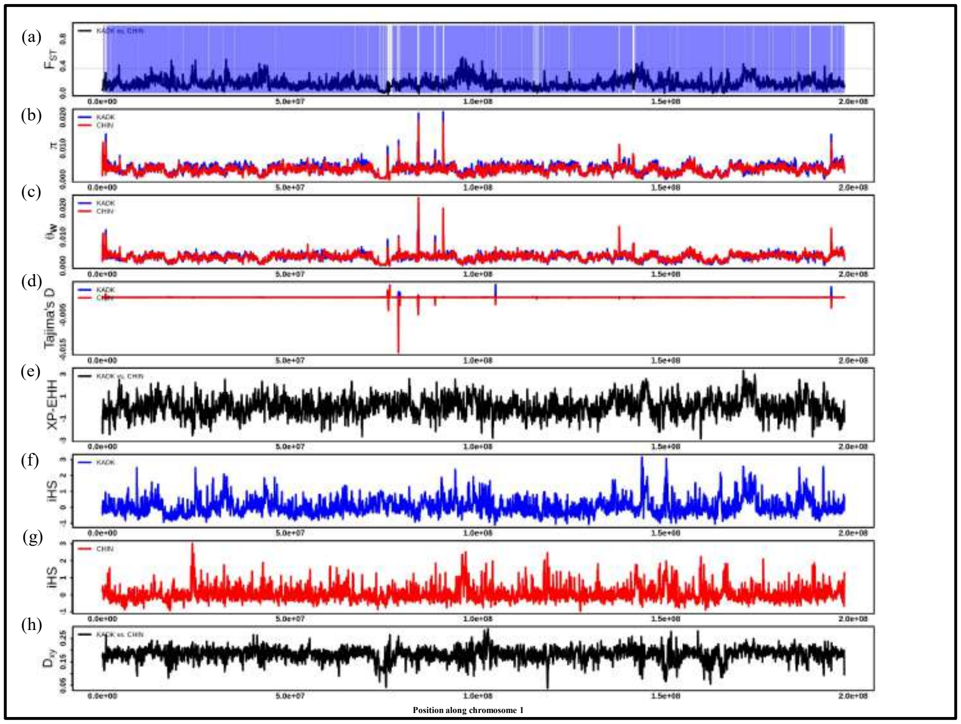


**Fig. S50. (a)** Pairwise $F_{ST}$ comparison between KADK and CHIN population along chromosome 1 using 50Kb windows. A horizontal black dotted line represents the 99 percentile $F_{ST}$ threshold. >80 percent callable region shown in transparent blue color while white color region represents <80 percent callable region. **b,c,d.** represents the $\pi$, Watterson theta, and Tajima`s D, respectively, where the solid blue line represents the KADK, and the solid red line represents the CHIN population. **(e)** Pairwise XP-EHH comparison between KADK and CHIN using 50Kb window. **f, g.** iHS results visualized in 50Kb window along the chromosome for KADK shown in solid blue color and CHIN shown in solid red color, respectively. **(h)** Dxy between KADK and CHIN.

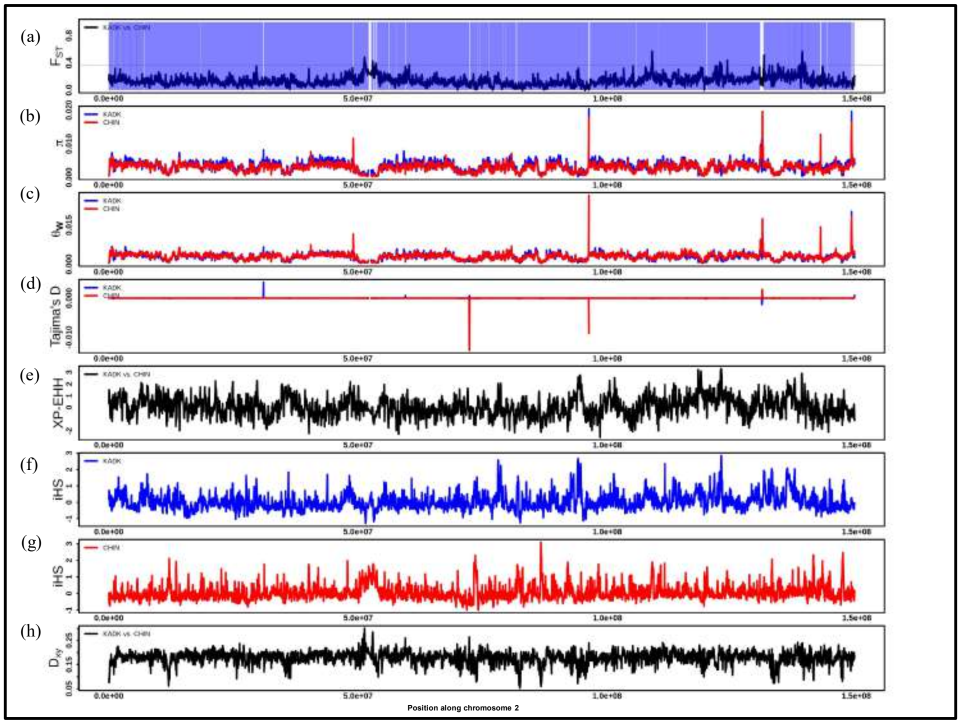


**Fig. S51. (a)** Pairwise $F_{ST}$ comparison between KADK and CHIN population along chromosome 2 using 50Kb windows. A horizontal black dotted line represents the 99 percentile $F_{ST}$ threshold. >80 percent callable region shown in transparent blue color while white color region represents <80 percent callable region. **b,c,d.** represents the $\pi$, Watterson theta, and Tajima`s D, respectively, where the solid blue line represents the KADK, and the solid red line represents the CHIN population. **(e)** Pairwise XP-EHH comparison between KADK and CHIN using 50Kb window. **f, g.** iHS results visualized in 50Kb window along the chromosome for KADK shown in solid blue color and CHIN shown in solid red color, respectively. **(h)** Dxy between KADK and CHIN.

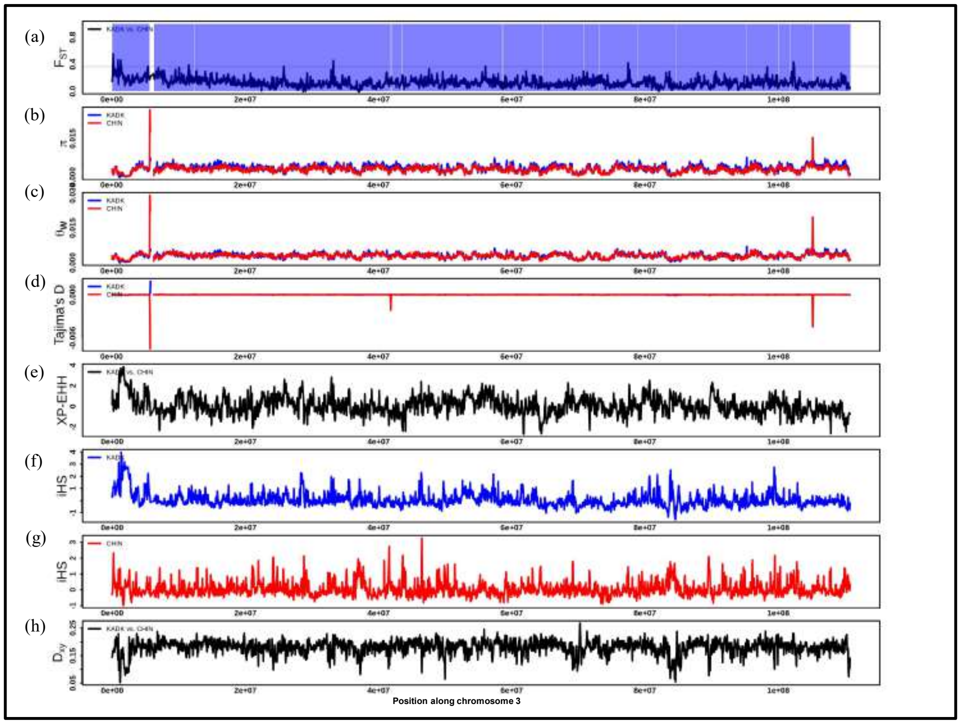


**Fig. S52. (a)** Pairwise $F_{ST}$ comparison between KADK and CHIN population along chromosome 3 using 50Kb windows. A horizontal black dotted line represents the 99 percentile $F_{ST}$ threshold. >80 percent callable region shown in transparent blue color while white color region represents <80 percent callable region. **b,c,d.** represents the $\pi$, Watterson theta, and Tajima`s D, respectively, where the solid blue line represents the KADK, and the solid red line represents the CHIN population. **(e)** Pairwise XP-EHH comparison between KADK and CHIN using 50Kb window. **f, g.** iHS results visualized in 50Kb window along the chromosome for KADK shown in solid blue color and CHIN shown in solid red color, respectively. **(h)** Dxy between KADK and CHIN.

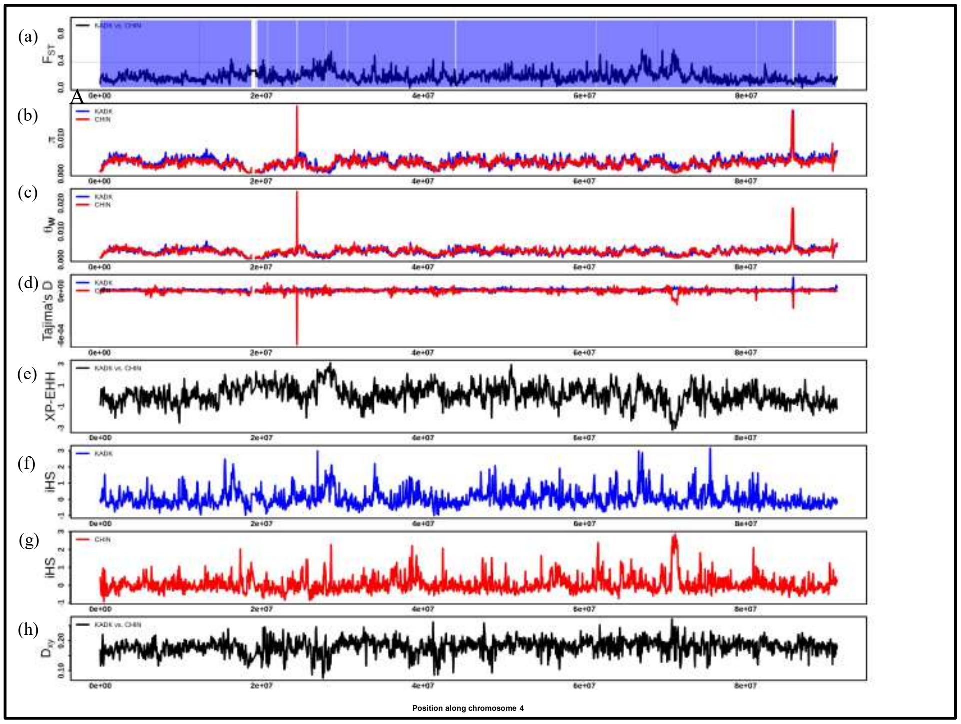


**Fig. S53. (a)** Pairwise $F_{ST}$ comparison between KADK and CHIN population along chromosome 4 using 50Kb windows. A horizontal black dotted line represents the 99 percentile $F_{ST}$ threshold. >80 percent callable region shown in transparent blue color while white color region represents <80 percent callable region. **b,c,d.** represents the π, Watterson theta, and Tajima`s D, respectively, where the solid blue line represents the KADK, and the solid red line represents the CHIN population. **(e)** Pairwise XP-EHH comparison between KADK and CHIN using 50Kb window. **f, g.** iHS results visualized in 50Kb window along the chromosome for KADK shown in solid blue color and CHIN shown in solid red color, respectively. **(h)** Dxy between KADK and CHIN.

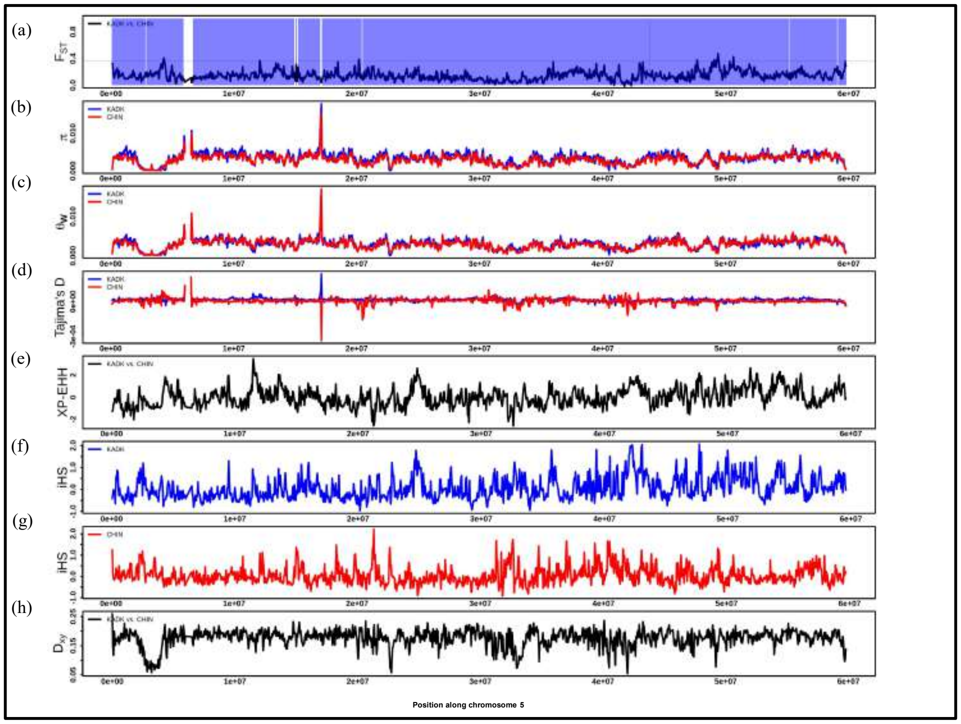


**Fig. S54. (a)** Pairwise $F_{ST}$ comparison between KADK and CHIN population along chromosome 5 using 50Kb windows. A horizontal black dotted line represents the 99 percentile $F_{ST}$ threshold. >80 percent callable region shown in transparent blue color while white color region represents <80 percent callable region. **b,c,d.** represents the π, Watterson theta, and Tajima`s D, respectively, where the solid blue line represents the KADK, and the solid red line represents the CHIN population. **(e)** Pairwise XP-EHH comparison between KADK and CHIN using 50Kb window. **f, g.** iHS results visualized in 50Kb window along the chromosome for KADK shown in solid blue color and CHIN shown in solid red color, respectively. **(h)** Dxy between KADK and CHIN.

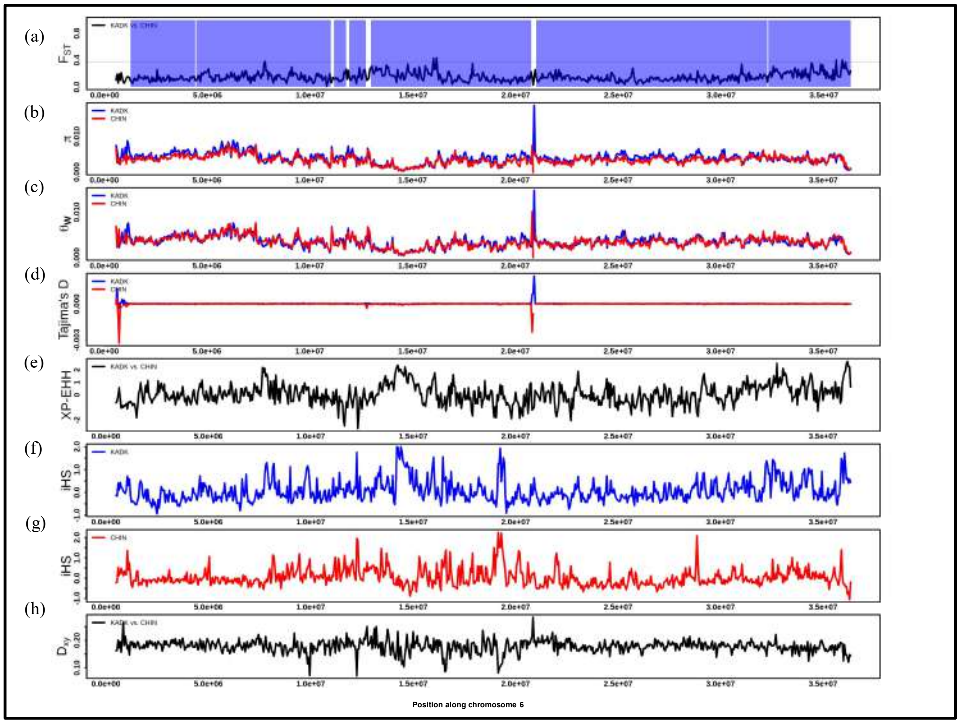


**Fig. S55. (a)** Pairwise $F_{ST}$ comparison between KADK and CHIN population along chromosome 6 using 50Kb windows. A horizontal black dotted line represents the 99 percentile $F_{ST}$ threshold. >80 percent callable region shown in transparent blue color while white color region represents <80 percent callable region. **b,c,d.** represents the $\pi$, Watterson theta, and Tajima`s D, respectively, where the solid blue line represents the KADK, and the solid red line represents the CHIN population. **(e)** Pairwise XP-EHH comparison between KADK and CHIN using 50Kb window. **f, g.** iHS results visualized in 50Kb window along the chromosome for KADK shown in solid blue color and CHIN shown in solid red color, respectively. **(h)** Dxy between KADK and CHIN.

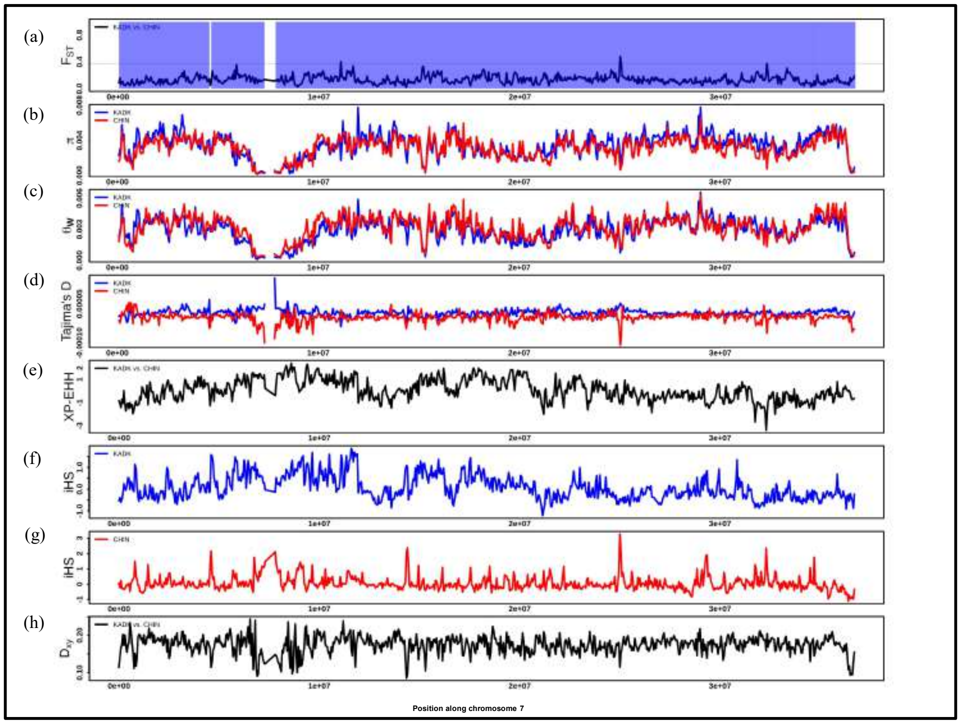


**Fig. S56. (a)** Pairwise $F_{ST}$ comparison between KADK and CHIN population along chromosome 7 using 50Kb windows. A horizontal black dotted line represents the 99 percentile $F_{ST}$ threshold. >80 percent callable region shown in transparent blue color while white color region represents <80 percent callable region. **b,c,d.** represents the $\pi$, Watterson theta, and Tajima`s D, respectively, where the solid blue line represents the KADK, and the solid red line represents the CHIN population. **(e)** Pairwise XP-EHH comparison between KADK and CHIN using 50Kb window. **f, g.** iHS results visualized in 50Kb window along the chromosome for KADK shown in solid blue color and CHIN shown in solid red color, respectively. **(h)** Dxy between KADK and CHIN.

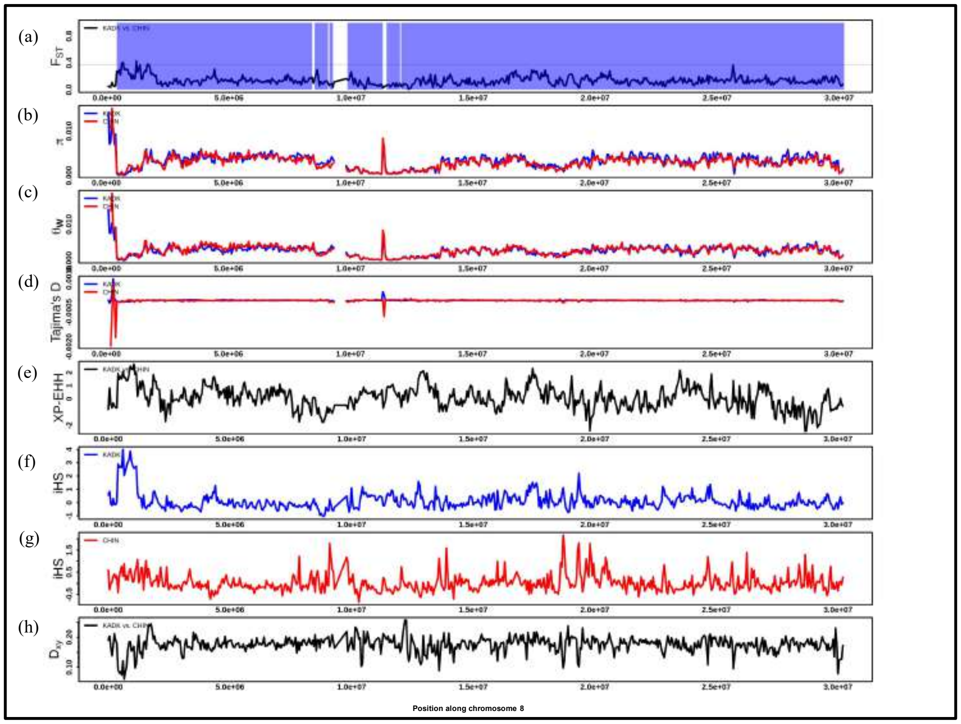

**Fig. S57. (a)** Pairwise $F_{ST}$ comparison between KADK and CHIN population along chromosome 8 using 50Kb windows. A horizontal black dotted line represents the 99 percentile $F_{ST}$ threshold. >80 percent callable region shown in transparent blue color while white color region represents <80 percent callable region. **b,c,d.** represents the $\pi$, Watterson theta, and Tajima`s D, respectively, where the solid blue line represents the KADK, and the solid red line represents the CHIN population. **(e)** Pairwise XP-EHH comparison between KADK and CHIN using 50Kb window. **f, g.** iHS results visualized in 50Kb window along the chromosome for KADK shown in solid blue color and CHIN shown in solid red color, respectively. **(h)** Dxy between KADK and CHIN.

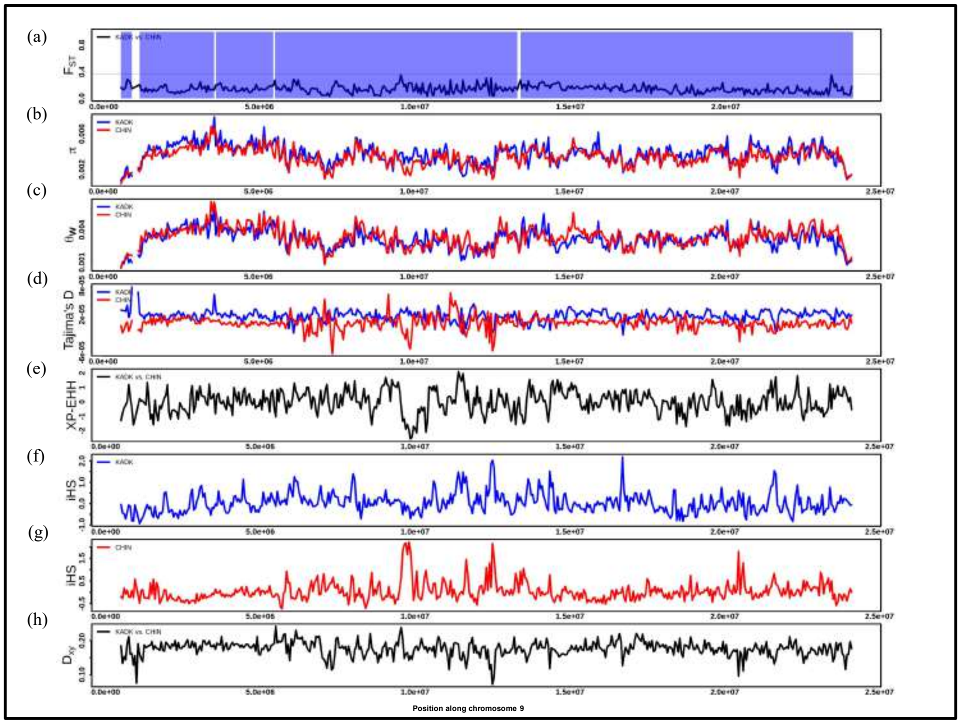

**Fig. S58. (a)** Pairwise $F_{ST}$ comparison between KADK and CHIN population along chromosome 9 using 50Kb windows. A horizontal black dotted line represents the 99 percentile $F_{ST}$ threshold. >80 percent callable region shown in transparent blue color while white color region represents <80 percent callable region. **b,c,d.** represents the $\pi$, Watterson theta, and Tajima`s D, respectively, where the solid blue line represents the KADK, and the solid red line represents the CHIN population. **(e)** Pairwise XP-EHH comparison between KADK and CHIN using 50Kb window. **f, g.** iHS results visualized in 50Kb window along the chromosome for KADK shown in solid blue color and CHIN shown in solid red color, respectively. **(h)** Dxy between KADK and CHIN.

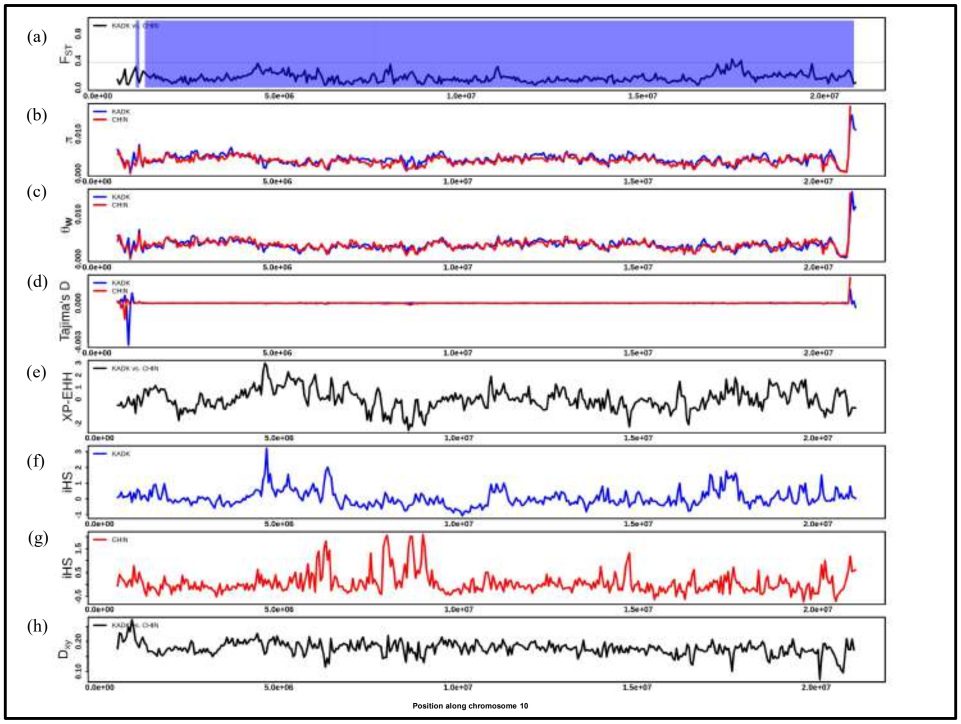


**Fig. S59. (a)** Pairwise $F_{ST}$ comparison between KADK and CHIN population along chromosome 10 using 50Kb windows. A horizontal black dotted line represents the 99 percentile $F_{ST}$ threshold. >80 percent callable region shown in transparent blue color while white color region represents <80 percent callable region. **b,c,d.** represents the $\pi$, Watterson theta, and Tajima`s D, respectively, where the solid blue line represents the KADK, and the solid red line represents the CHIN population. **(e)** Pairwise XP-EHH comparison between KADK and CHIN using 50Kb window. **f, g.** iHS results visualized in 50Kb window along the chromosome for KADK shown in solid blue color and CHIN shown in solid red color, respectively. **(h)** Dxy between KADK and CHIN.

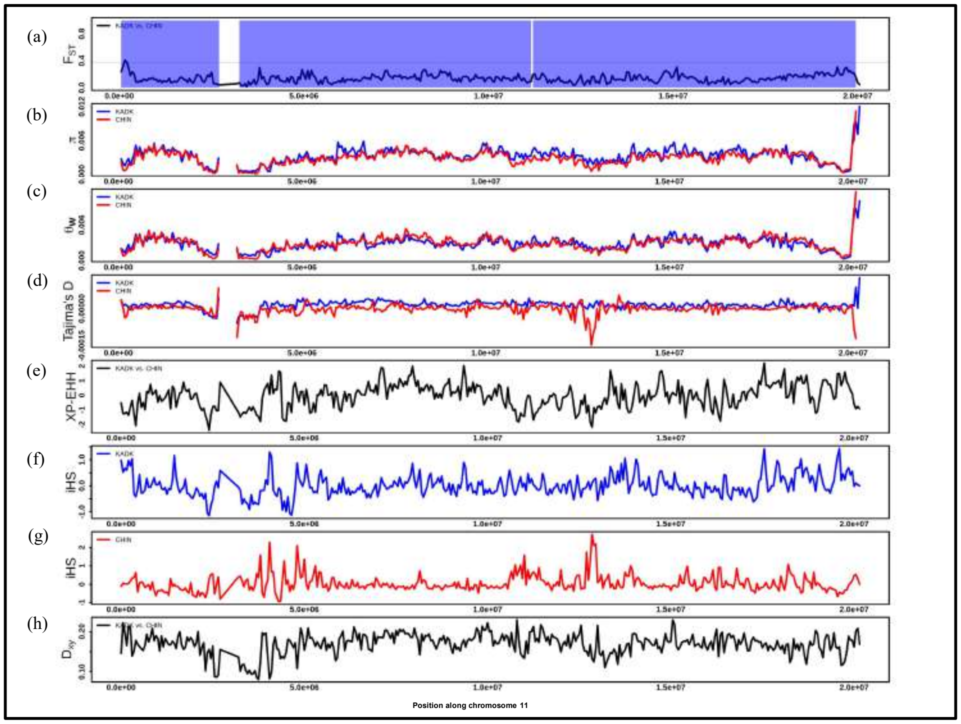

**Fig. S60. (a)** Pairwise $F_{ST}$ comparison between KADK and CHIN population along chromosome 11 using 50Kb windows. A horizontal black dotted line represents the 99 percentile $F_{ST}$ threshold. >80 percent callable region shown in transparent blue color while white color region represents <80 percent callable region. **b,c,d.** represents the $\pi$, Watterson theta, and Tajima`s D, respectively, where the solid blue line represents the KADK, and the solid red line represents the CHIN population. **(e)** Pairwise XP-EHH comparison between KADK and CHIN using 50Kb window. **f, g.** iHS results visualized in 50Kb window along the chromosome for KADK shown in solid blue color and CHIN shown in solid red color, respectively. **(h)** Dxy between KADK and CHIN.

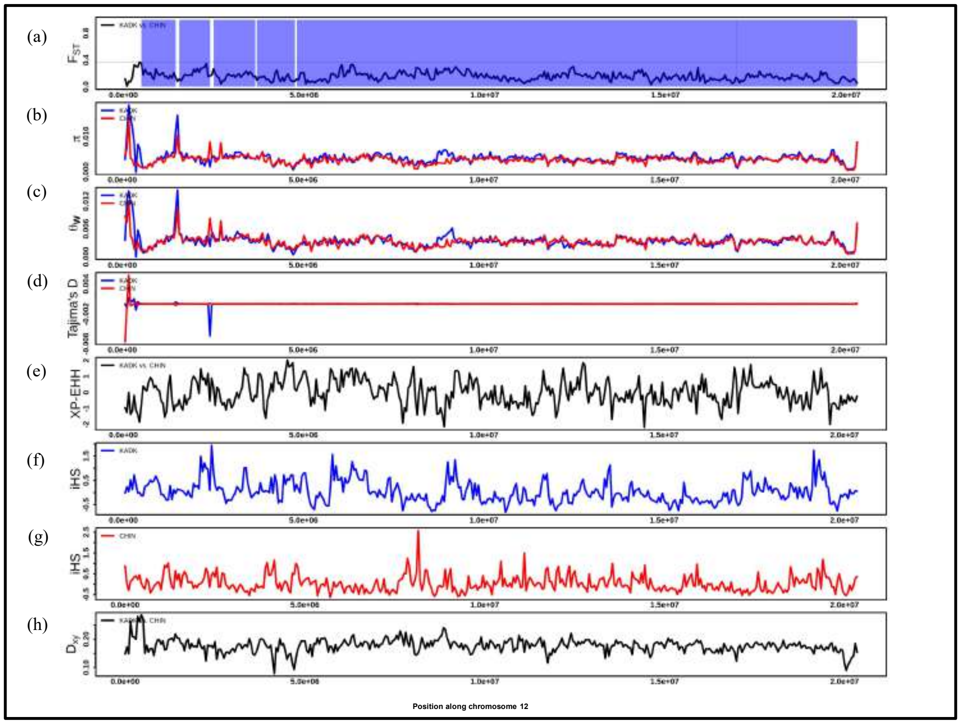


**Fig. S61. (a)** Pairwise $F_{ST}$ comparison between KADK and CHIN population along chromosome 12 using 50Kb windows. A horizontal black dotted line represents the 99 percentile $F_{ST}$ threshold. >80 percent callable region shown in transparent blue color while white color region represents <80 percent callable region. **b,c,d.** represents the $\pi$, Watterson theta, and Tajima`s D, respectively, where the solid blue line represents the KADK, and the solid red line represents the CHIN population. **(e)** Pairwise XP-EHH comparison between KADK and CHIN using 50Kb window. **f, g.** iHS results visualized in 50Kb window along the chromosome for KADK shown in solid blue color and CHIN shown in solid red color, respectively. **(h)** Dxy between KADK and CHIN.

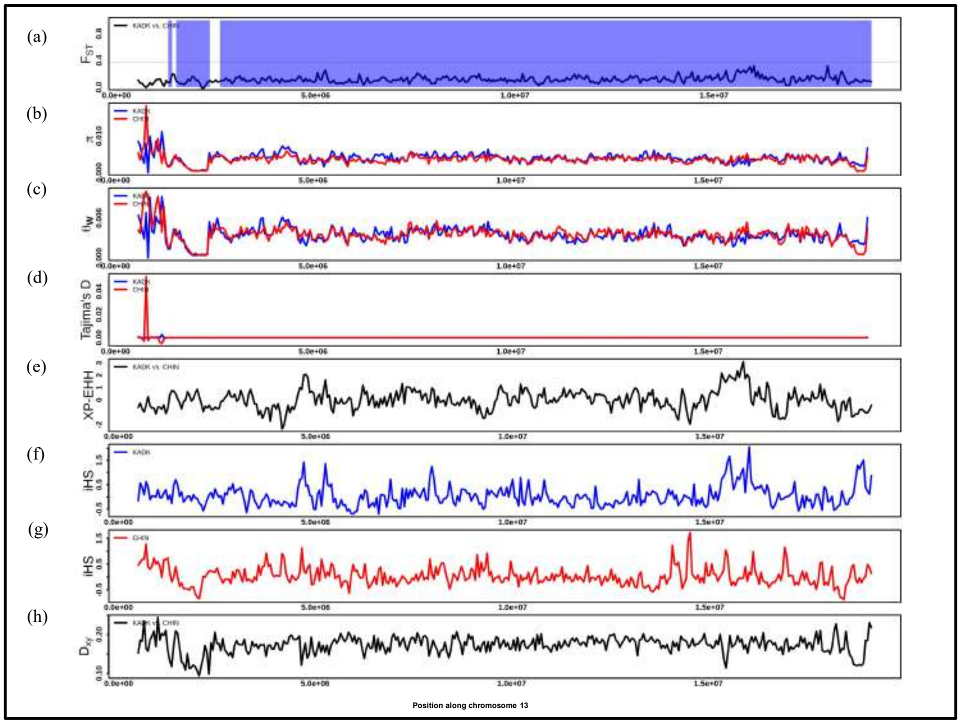


**Fig. S62. (a)** Pairwise $F_{ST}$ comparison between KADK and CHIN population along chromosome 13 using 50Kb windows. A horizontal black dotted line represents the 99 percentile $F_{ST}$ threshold. >80 percent callable region shown in transparent blue color while white color region represents <80 percent callable region. **b,c,d.** represents the $\pi$, Watterson theta, and Tajima`s D, respectively, where the solid blue line represents the KADK, and the solid red line represents the CHIN population. **(e)** Pairwise XP-EHH comparison between KADK and CHIN using 50Kb window. **f, g.** iHS results visualized in 50Kb window along the chromosome for KADK shown in solid blue color and CHIN shown in solid red color, respectively. **(h)** Dxy between KADK and CHIN.

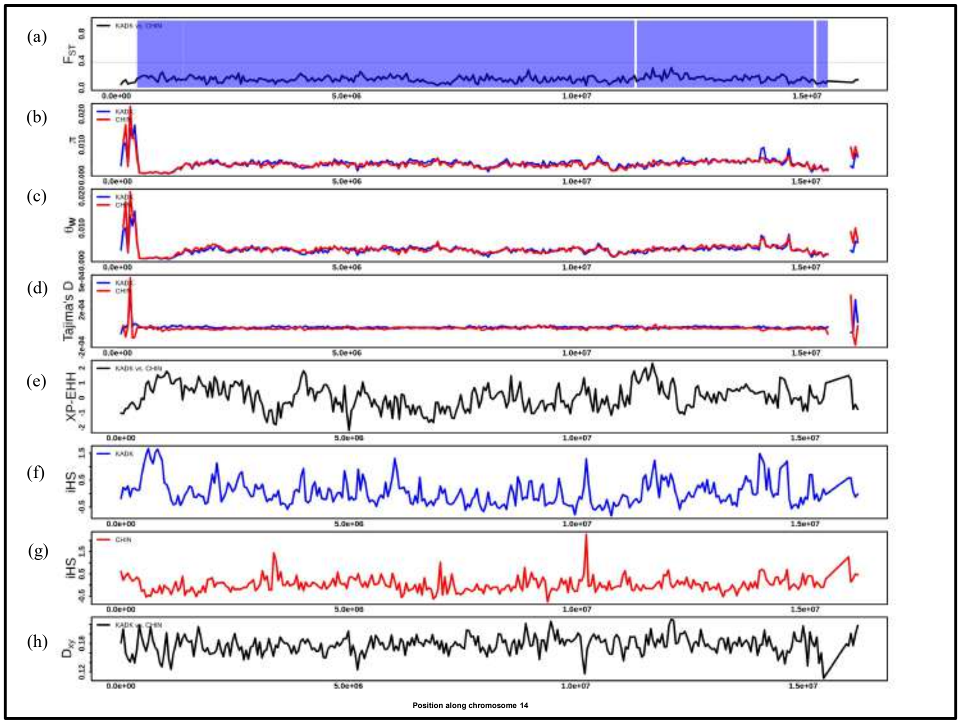


**Fig. S63. (a)** Pairwise $F_{ST}$ comparison between KADK and CHIN population along chromosome 14 using 50Kb windows. A horizontal black dotted line represents the 99 percentile $F_{ST}$ threshold. >80 percent callable region shown in transparent blue color while white color region represents <80 percent callable region. **b,c,d.** represents the $\pi$, Watterson theta, and Tajima`s D, respectively, where the solid blue line represents the KADK, and the solid red line represents the CHIN population. **(e)** Pairwise XP-EHH comparison between KADK and CHIN using 50Kb window. **f, g.** iHS results visualized in 50Kb window along the chromosome for KADK shown in solid blue color and CHIN shown in solid red color, respectively. **(h)** Dxy between KADK and CHIN.

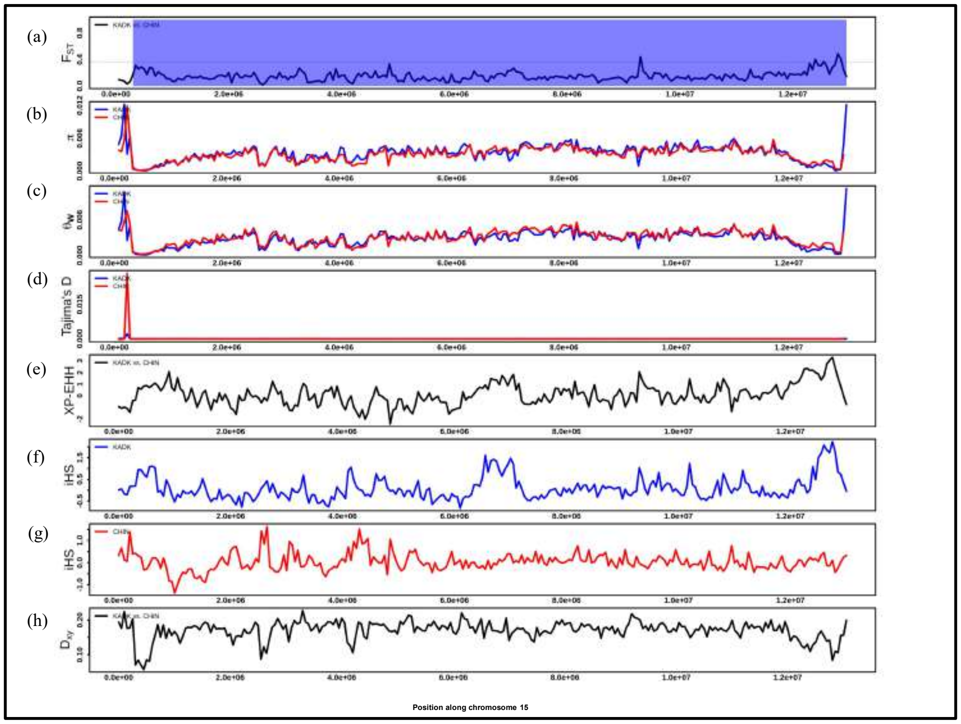


**Fig. S64. (a)** Pairwise $F_{ST}$ comparison between KADK and CHIN population along chromosome 15 using 50Kb windows. A horizontal black dotted line represents the 99 percentile $F_{ST}$ threshold. >80 percent callable region shown in transparent blue color while white color region represents <80 percent callable region. **b,c,d.** represents the $\pi$, Watterson theta, and Tajima`s D, respectively, where the solid blue line represents the KADK, and the solid red line represents the CHIN population. **(e)** Pairwise XP-EHH comparison between KADK and CHIN using 50Kb window. **f, g.** iHS results visualized in 50Kb window along the chromosome for KADK shown in solid blue color and CHIN shown in solid red color, respectively. **(h)** Dxy between KADK and CHIN.

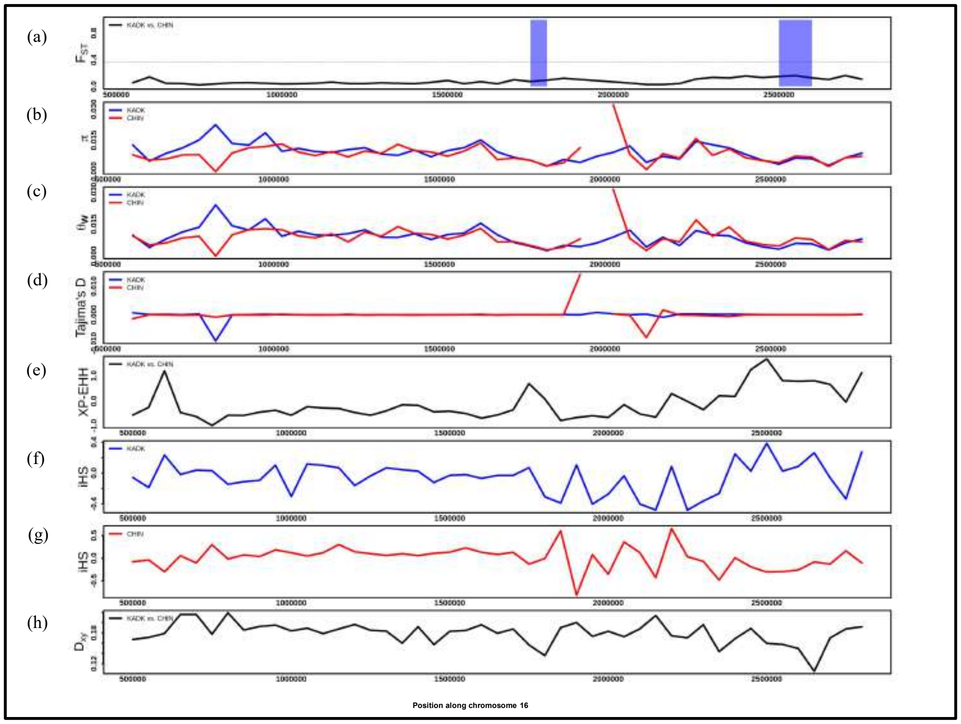


**Fig. S65. (a)** Pairwise $F_{ST}$ comparison between KADK and CHIN population along chromosome 16 using 50Kb windows. A horizontal black dotted line represents the 99 percentile $F_{ST}$ threshold. >80 percent callable region shown in transparent blue color while white color region represents <80 percent callable region. **b,c,d.** represents the π, Watterson theta, and Tajima`s D, respectively, where the solid blue line represents the KADK, and the solid red line represents the CHIN population. **(e)** Pairwise XP-EHH comparison between KADK and CHIN using 50Kb window. **f, g.** iHS results visualized in 50Kb window along the chromosome for KADK shown in solid blue color and CHIN shown in solid red color, respectively. **(h)** Dxy between KADK and CHIN.

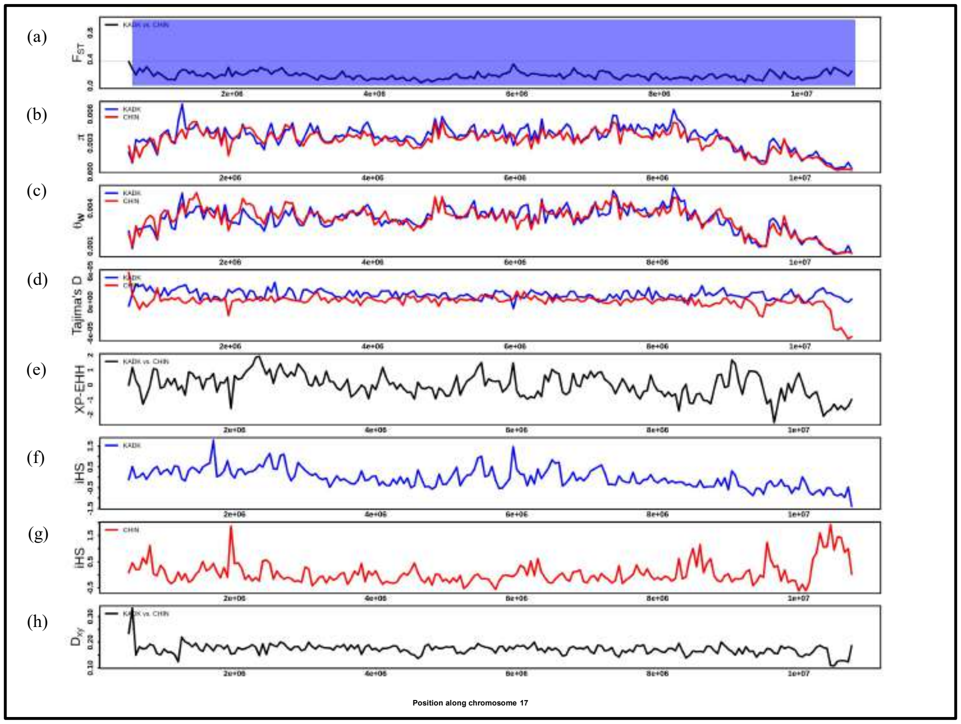


**Fig. S66. (a)** Pairwise $F_{ST}$ comparison between KADK and CHIN population along chromosome 17 using 50Kb windows. A horizontal black dotted line represents the 99 percentile $F_{ST}$ threshold. >80 percent callable region shown in transparent blue color while white color region represents <80 percent callable region. **b,c,d.** represents the $\pi$, Watterson theta, and Tajima`s D, respectively, where the solid blue line represents the KADK, and the solid red line represents the CHIN population. **(e)** Pairwise XP-EHH comparison between KADK and CHIN using 50Kb window. **f, g.** iHS results visualized in 50Kb window along the chromosome for KADK shown in solid blue color and CHIN shown in solid red color, respectively. **(h)** Dxy between KADK and CHIN.

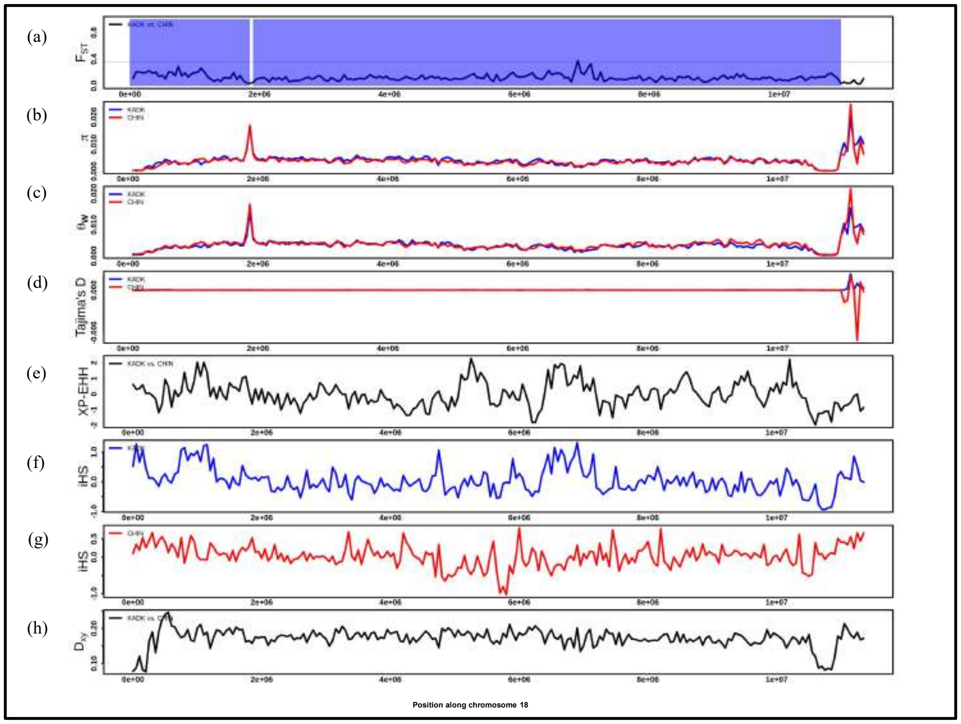


**Fig. S67. (a)** Pairwise $F_{ST}$ comparison between KADK and CHIN population along chromosome 18 using 50Kb windows. A horizontal black dotted line represents the 99 percentile $F_{ST}$ threshold. >80 percent callable region shown in transparent blue color while white color region represents <80 percent callable region. **b,c,d.** represents the $\pi$, Watterson theta, and Tajima`s D, respectively, where the solid blue line represents the KADK, and the solid red line represents the CHIN population. **(e)** Pairwise XP-EHH comparison between KADK and CHIN using 50Kb window. **f, g.** iHS results visualized in 50Kb window along the chromosome for KADK shown in solid blue color and CHIN shown in solid red color, respectively. **(h)** Dxy between KADK and CHIN.

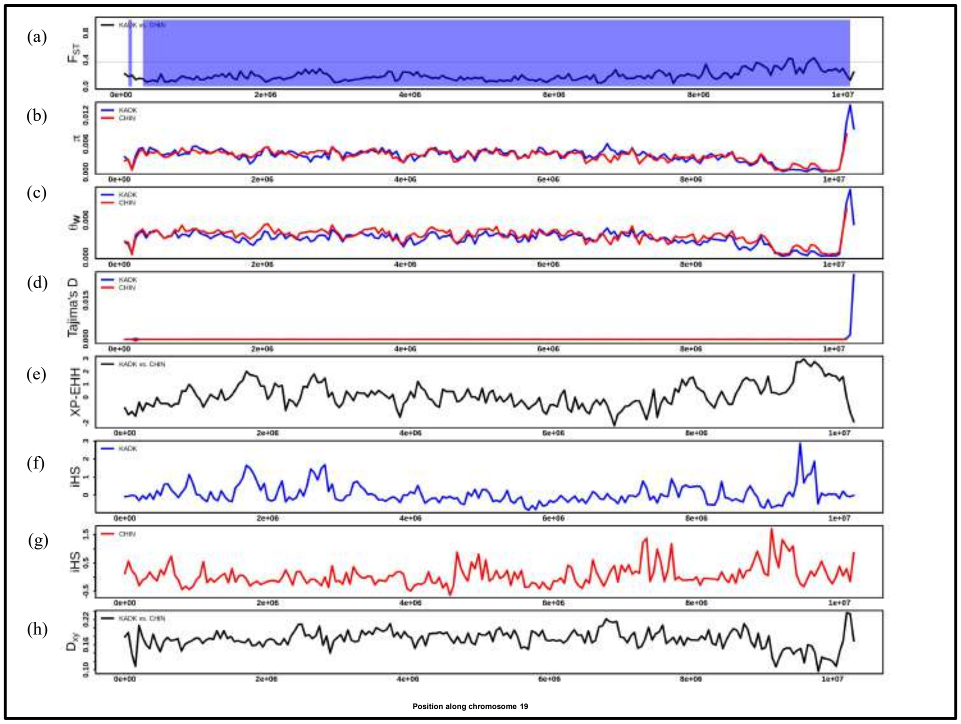


**Fig. S68. (a)** Pairwise $F_{ST}$ comparison between KADK and CHIN population along chromosome 19 using 50Kb windows. A horizontal black dotted line represents the 99 percentile $F_{ST}$ threshold. >80 percent callable region shown in transparent blue color while white color region represents <80 percent callable region. **b,c,d.** represents the $\pi$, Watterson theta, and Tajima`s D, respectively, where the solid blue line represents the KADK, and the solid red line represents the CHIN population. **(e)** Pairwise XP-EHH comparison between KADK and CHIN using 50Kb window. **f, g.** iHS results visualized in 50Kb window along the chromosome for KADK shown in solid blue color and CHIN shown in solid red color, respectively. **(h)** Dxy between KADK and CHIN.

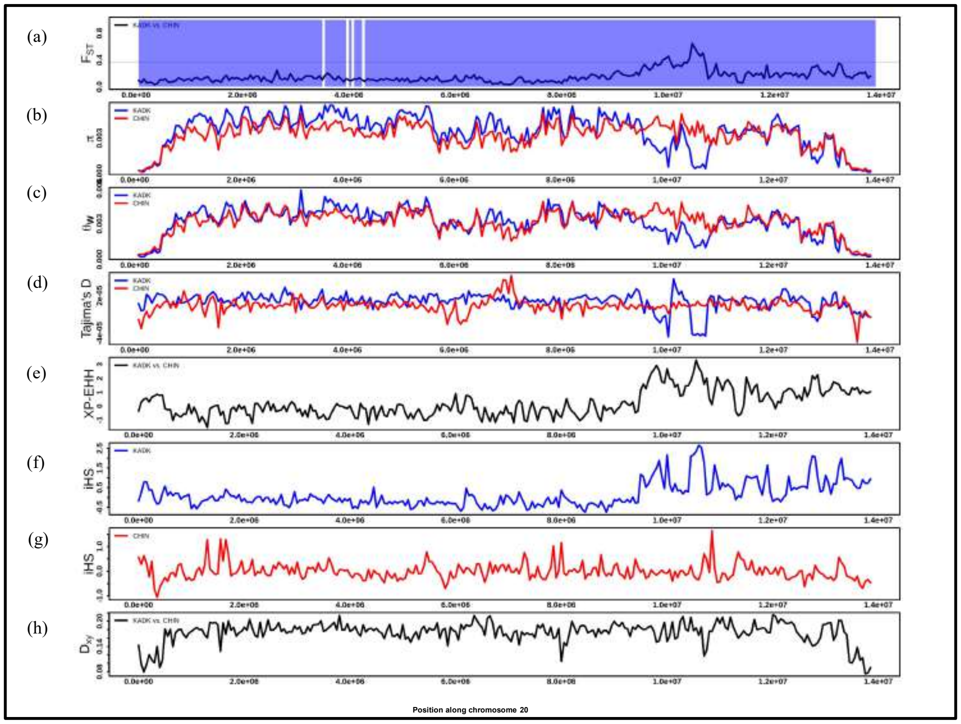


**Fig. S69. (a)** Pairwise $F_{ST}$ comparison between KADK and CHIN population along chromosome 20 using 50Kb windows. A horizontal black dotted line represents the 99 percentile $F_{ST}$ threshold. >80 percent callable region shown in transparent blue color while white color region represents <80 percent callable region. **b,c,d.** represents the $\pi$, Watterson theta, and Tajima`s D, respectively, where the solid blue line represents the KADK, and the solid red line represents the CHIN population. **(e)** Pairwise XP-EHH comparison between KADK and CHIN using 50Kb window. **f, g.** iHS results visualized in 50Kb window along the chromosome for KADK shown in solid blue color and CHIN shown in solid red color, respectively. **(h)** Dxy between KADK and CHIN.

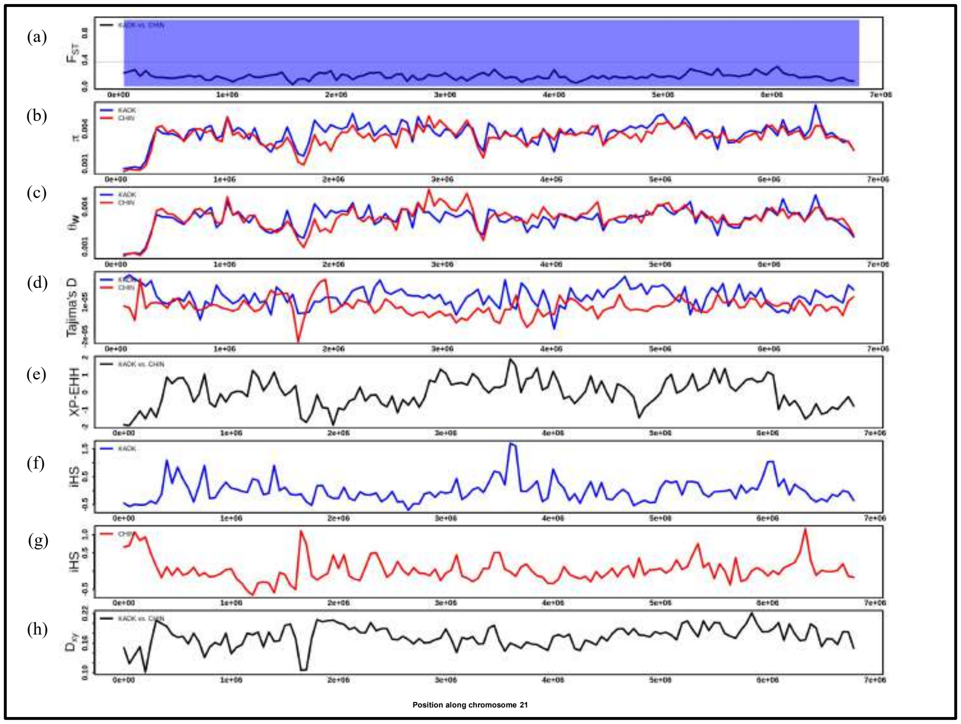

**Fig. S70. (a)** Pairwise $F_{ST}$ comparison between KADK and CHIN population along chromosome 21 using 50Kb windows. A horizontal black dotted line represents the 99 percentile $F_{ST}$ threshold. >80 percent callable region shown in transparent blue color while white color region represents <80 percent callable region. **b,c,d.** represents the $\pi$, Watterson theta, and Tajima`s D, respectively, where the solid blue line represents the KADK, and the solid red line represents the CHIN population. **(e)** Pairwise XP-EHH comparison between KADK and CHIN using 50Kb window. **f, g.** iHS results visualized in 50Kb window along the chromosome for KADK shown in solid blue color and CHIN shown in solid red color, respectively. **(h)** Dxy between KADK and CHIN.

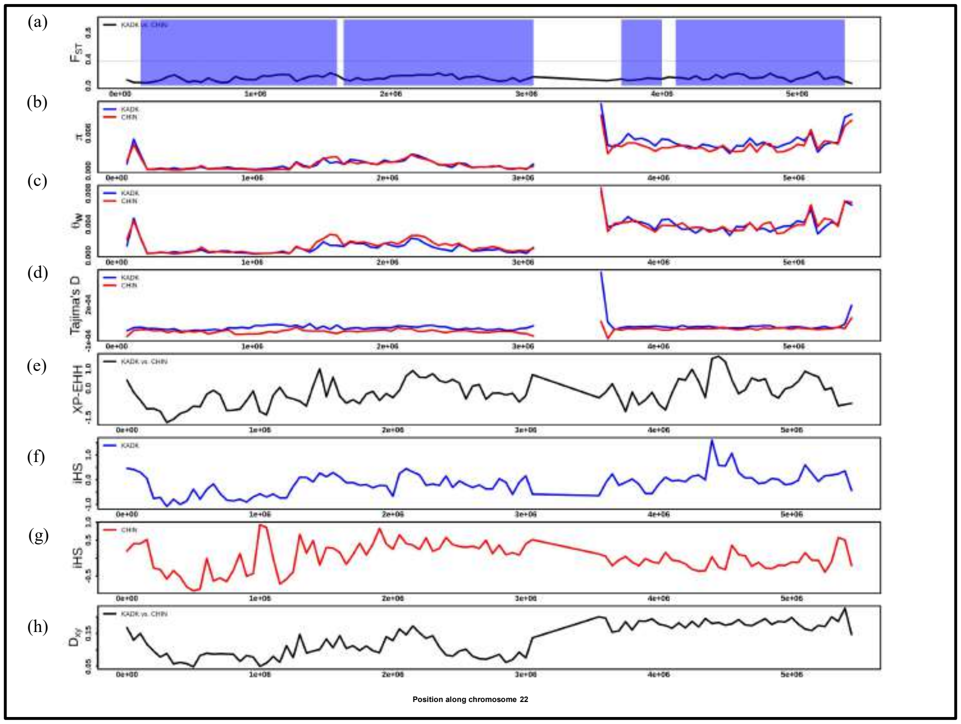


**Fig. S71. (a)** Pairwise $F_{ST}$ comparison between KADK and CHIN population along chromosome 22 using 50Kb windows. A horizontal black dotted line represents the 99 percentile $F_{ST}$ threshold. >80 percent callable region shown in transparent blue color while white color region represents <80 percent callable region. **b,c,d.** represents the $\pi$, Watterson theta, and Tajima`s D, respectively, where the solid blue line represents the KADK, and the solid red line represents the CHIN population. **(e)** Pairwise XP-EHH comparison between KADK and CHIN using 50Kb window. **f, g.** iHS results visualized in 50Kb window along the chromosome for KADK shown in solid blue color and CHIN shown in solid red color, respectively. **(h)** Dxy between KADK and CHIN.

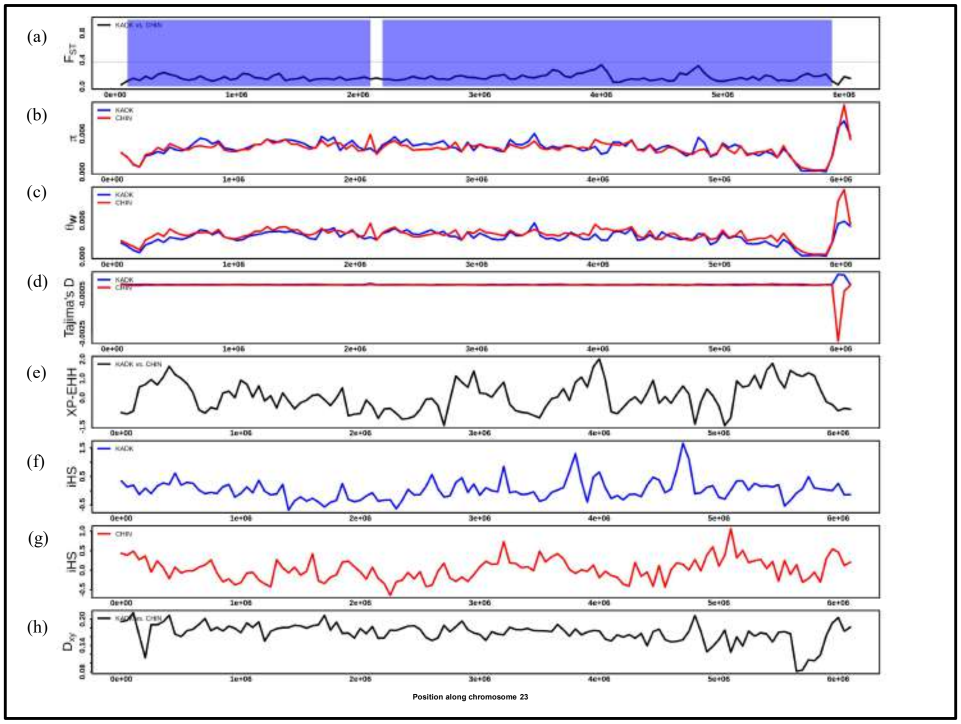

**Fig. S72. (a)** Pairwise $F_{ST}$ comparison between KADK and CHIN population along chromosome 23 using 50Kb windows. A horizontal black dotted line represents the 99 percentile $F_{ST}$ threshold. >80 percent callable region shown in transparent blue color while white color region represents <80 percent callable region. **b,c,d.** represents the $\pi$, Watterson theta, and Tajima`s D, respectively, where the solid blue line represents the KADK, and the solid red line represents the CHIN population. **(e)** Pairwise XP-EHH comparison between KADK and CHIN using 50Kb window. **f, g.** iHS results visualized in 50Kb window along the chromosome for KADK shown in solid blue color and CHIN shown in solid red color, respectively. **(h)** Dxy between KADK and CHIN.

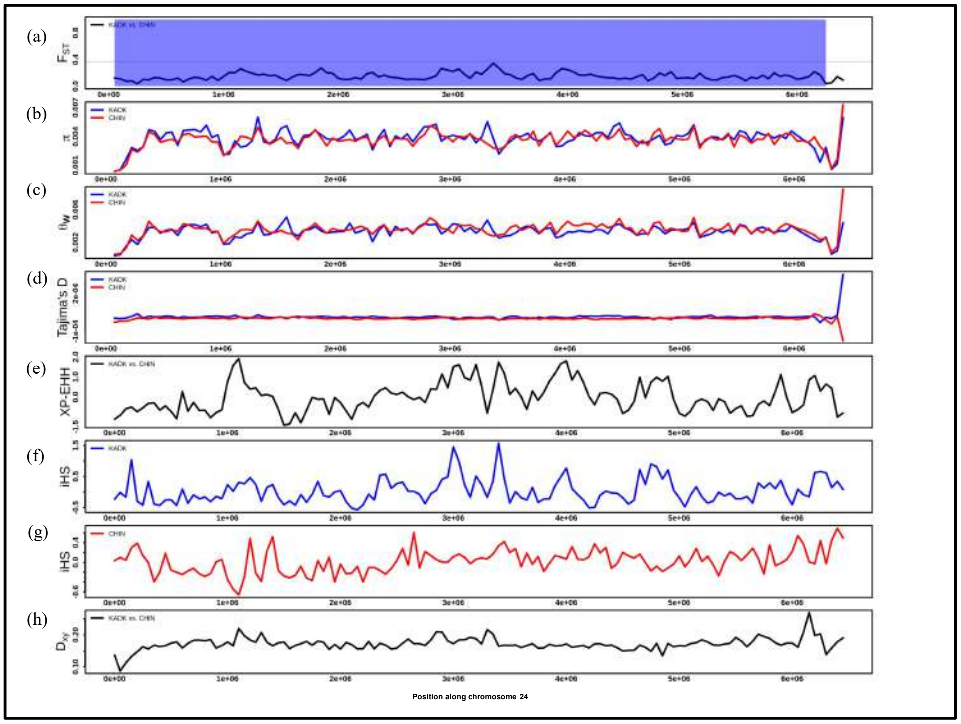


**Fig. S73. (a)** Pairwise $F_{ST}$ comparison between KADK and CHIN population along chromosome 24 using 50Kb windows. A horizontal black dotted line represents the 99 percentile $F_{ST}$ threshold. >80 percent callable region shown in transparent blue color while white color region represents <80 percent callable region. **b,c,d.** represents the $\pi$, Watterson theta, and Tajima`s D, respectively, where the solid blue line represents the KADK, and the solid red line represents the CHIN population. **(e)** Pairwise XP-EHH comparison between KADK and CHIN using 50Kb window. **f, g.** iHS results visualized in 50Kb window along the chromosome for KADK shown in solid blue color and CHIN shown in solid red color, respectively. **(h)** Dxy between KADK and CHIN.

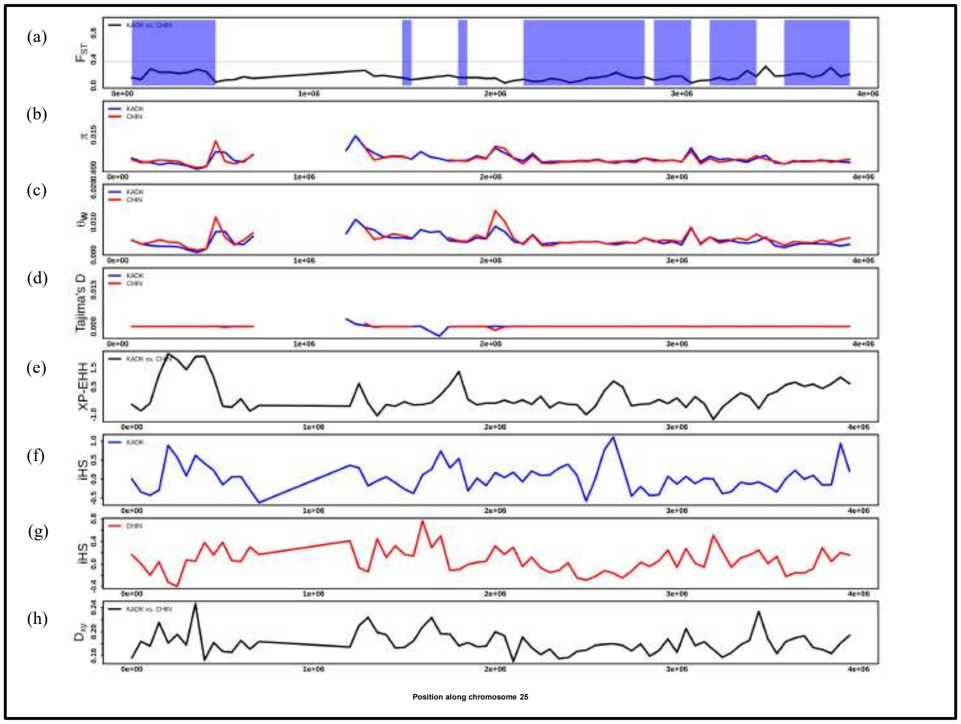


**Fig. S74. (a)** Pairwise $F_{ST}$ comparison between KADK and CHIN population along chromosome 25 using 50Kb windows. A horizontal black dotted line represents the 99 percentile $F_{ST}$ threshold. >80 percent callable region shown in transparent blue color while white color region represents <80 percent callable region. **b,c,d.** represents the $\pi$, Watterson theta, and Tajima`s D, respectively, where the solid blue line represents the KADK, and the solid red line represents the CHIN population. **(e)** Pairwise XP-EHH comparison between KADK and CHIN using 50Kb window. **f, g.** iHS results visualized in 50Kb window along the chromosome for KADK shown in solid blue color and CHIN shown in solid red color, respectively. **(h)** Dxy between KADK and CHIN.

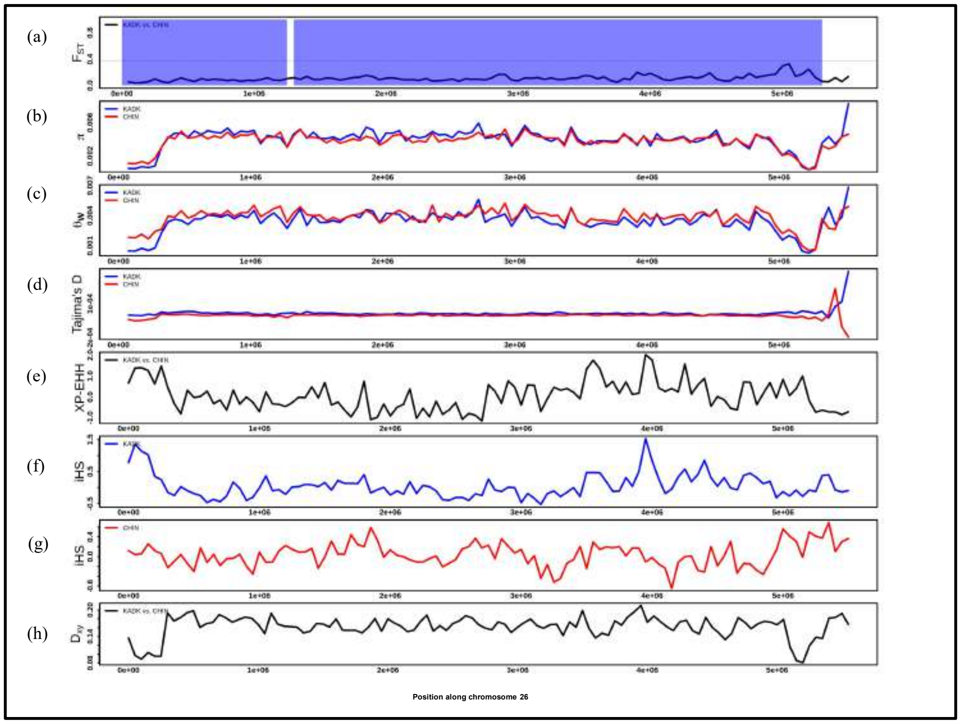


**Fig. S75. (a)** Pairwise $F_{ST}$ comparison between KADK and CHIN population along chromosome 26 using 50Kb windows. A horizontal black dotted line represents the 99 percentile $F_{ST}$ threshold. >80 percent callable region shown in transparent blue color while white color region represents <80 percent callable region. **b,c,d.** represents the $\pi$, Watterson theta, and Tajima`s D, respectively, where the solid blue line represents the KADK, and the solid red line represents the CHIN population. **(e)** Pairwise XP-EHH comparison between KADK and CHIN using 50Kb window. **f, g.** iHS results visualized in 50Kb window along the chromosome for KADK shown in solid blue color and CHIN shown in solid red color, respectively. **(h)** Dxy between KADK and CHIN.

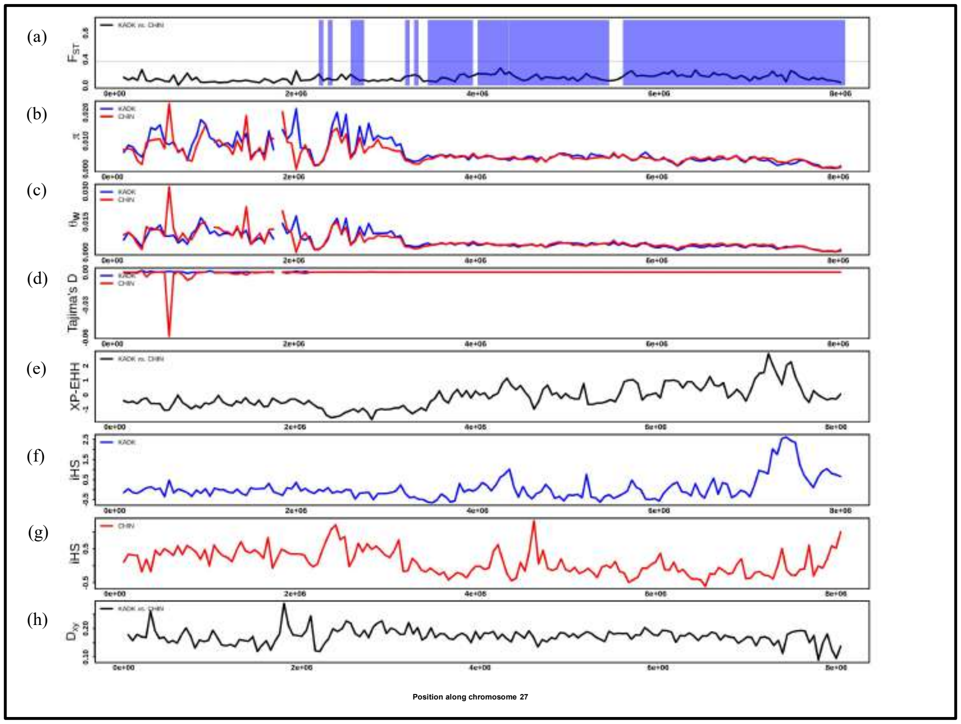

**Fig. S76. (a)** Pairwise $F_{ST}$ comparison between KADK and CHIN population along chromosome 27 using 50Kb windows. A horizontal black dotted line represents the 99 percentile $F_{ST}$ threshold. >80 percent callable region shown in transparent blue color while white color region represents <80 percent callable region. **b,c,d.** represents the $\pi$, Watterson theta, and Tajima`s D, respectively, where the solid blue line represents the KADK, and the solid red line represents the CHIN population. **(e)** Pairwise XP-EHH comparison between KADK and CHIN using 50Kb window. **f, g.** iHS results visualized in 50Kb window along the chromosome for KADK shown in solid blue color and CHIN shown in solid red color, respectively. **(h)** Dxy between KADK and CHIN.

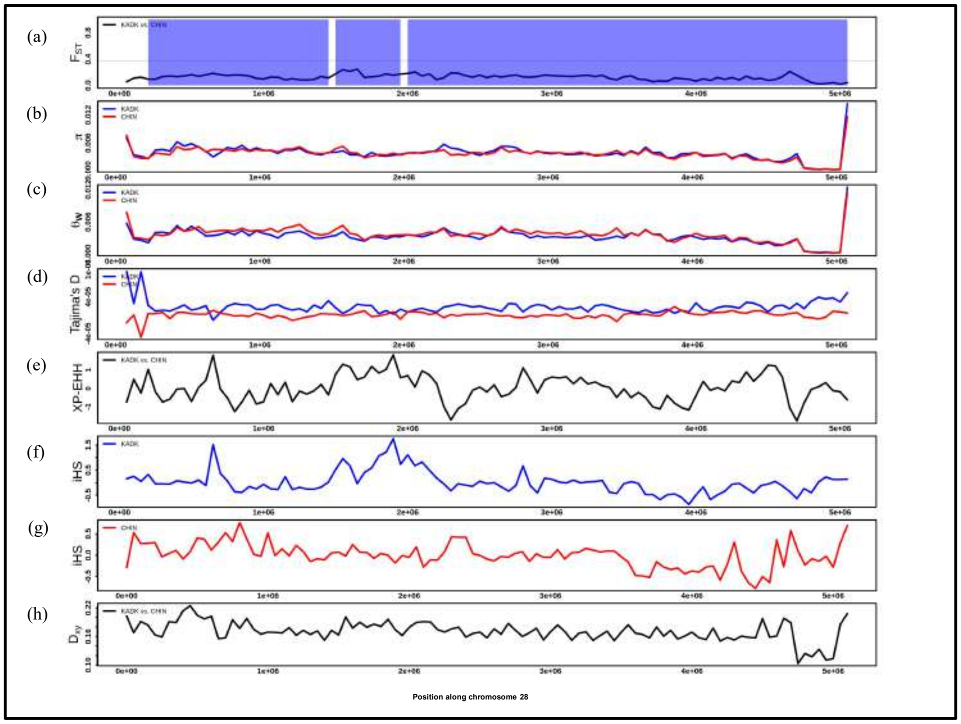


**Fig. S77. (a)** Pairwise $F_{ST}$ comparison between KADK and CHIN population along chromosome 28 using 50Kb windows. A horizontal black dotted line represents the 99 percentile $F_{ST}$ threshold. >80 percent callable region shown in transparent blue color while white color region represents <80 percent callable region. **b,c,d.** represents the π, Watterson theta, and Tajima`s D, respectively, where the solid blue line represents the KADK, and the solid red line represents the CHIN population. **(e)** Pairwise XP-EHH comparison between KADK and CHIN using 50Kb window. **f, g.** iHS results visualized in 50Kb window along the chromosome for KADK shown in solid blue color and CHIN shown in solid red color, respectively. **(h)** Dxy between KADK and CHIN.

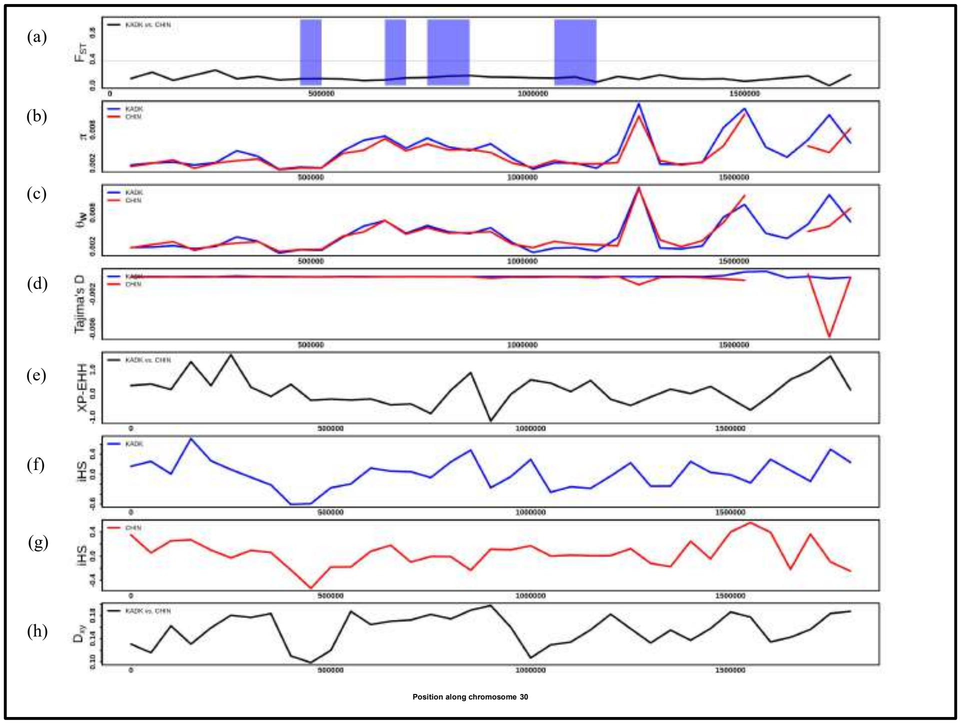


**Fig. S78. (a)** Pairwise $F_{ST}$ comparison between KADK and CHIN population along chromosome 30 using 50Kb windows. A horizontal black dotted line represents the 99 percentile $F_{ST}$ threshold. >80 percent callable region shown in transparent blue color while white color region represents <80 percent callable region. **b,c,d.** represents the π, Watterson theta, and Tajima`s D, respectively, where the solid blue line represents the KADK, and the solid red line represents the CHIN population. **(e)** Pairwise XP-EHH comparison between KADK and CHIN using 50Kb window. **f, g.** iHS results visualized in 50Kb window along the chromosome for KADK shown in solid blue color and CHIN shown in solid red color, respectively. **(h)** Dxy between KADK and CHIN.

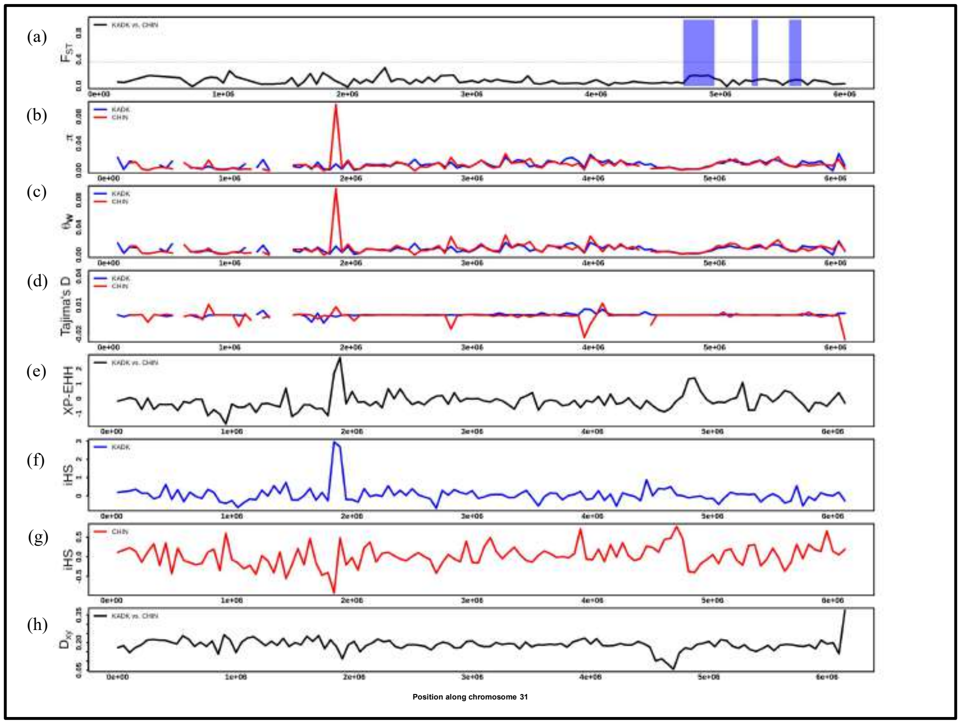


**Fig. S79. (a)** Pairwise $F_{ST}$ comparison between KADK and CHIN population along chromosome 31 using 50Kb windows. A horizontal black dotted line represents the 99 percentile $F_{ST}$ threshold. >80 percent callable region shown in transparent blue color while white color region represents <80 percent callable region. **b,c,d.** represents the π, Watterson theta, and Tajima`s D, respectively, where the solid blue line represents the KADK, and the solid red line represents the CHIN population. **(e)** Pairwise XP-EHH comparison between KADK and CHIN using 50Kb window. **f, g.** iHS results visualized in 50Kb window along the chromosome for KADK shown in solid blue color and CHIN shown in solid red color, respectively. **(h)** Dxy between KADK and CHIN.

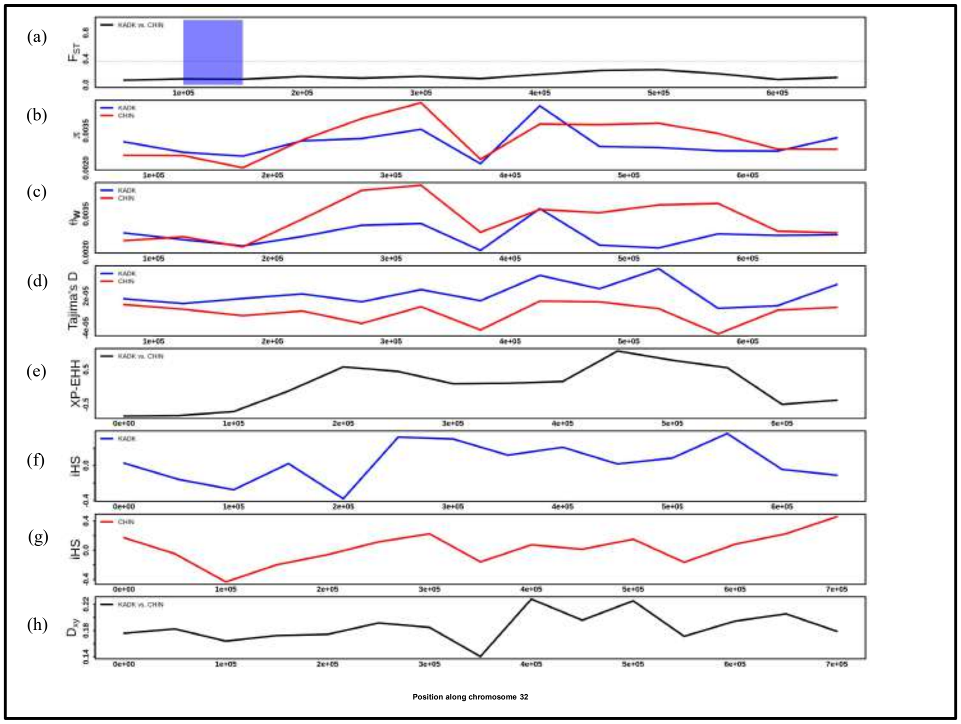


**Fig. S80. (a)** Pairwise $F_{ST}$ comparison between KADK and CHIN population along chromosome 32 using 50Kb windows. A horizontal black dotted line represents the 99 percentile $F_{ST}$ threshold. >80 percent callable region shown in transparent blue color while white color region represents <80 percent callable region. **b,c,d.** represents the $\pi$, Watterson theta, and Tajima`s D, respectively, where the solid blue line represents the KADK, and the solid red line represents the CHIN population. **(e)** Pairwise XP-EHH comparison between KADK and CHIN using 50Kb window. **f, g.** iHS results visualized in 50Kb window along the chromosome for KADK shown in solid blue color and CHIN shown in solid red color, respectively. **(h)** Dxy between KADK and CHIN.

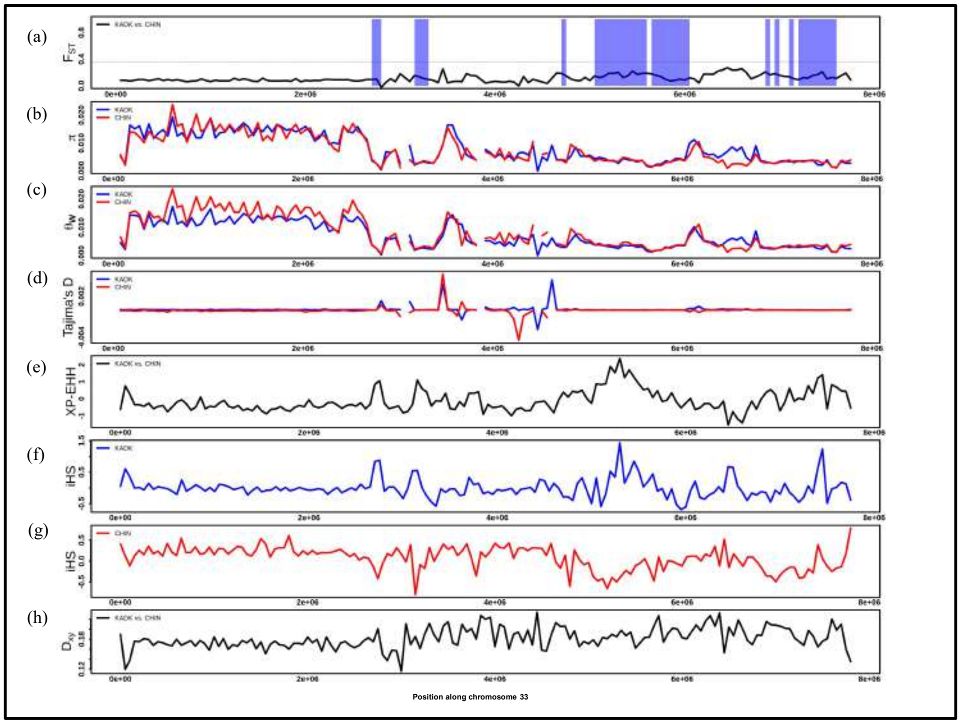


**Fig. S81. (a)** Pairwise $F_{ST}$ comparison between KADK and CHIN population along chromosome 33 using 50Kb windows. A horizontal black dotted line represents the 99 percentile $F_{ST}$ threshold. >80 percent callable region shown in transparent blue color while white color region represents <80 percent callable region. **b,c,d.** represents the π, Watterson theta, and Tajima`s D, respectively, where the solid blue line represents the KADK, and the solid red line represents the CHIN population. **(e)** Pairwise XP-EHH comparison between KADK and CHIN using 50Kb window. **f, g.** iHS results visualized in 50Kb window along the chromosome for KADK shown in solid blue color and CHIN shown in solid red color, respectively. **(h)** Dxy between KADK and CHIN.

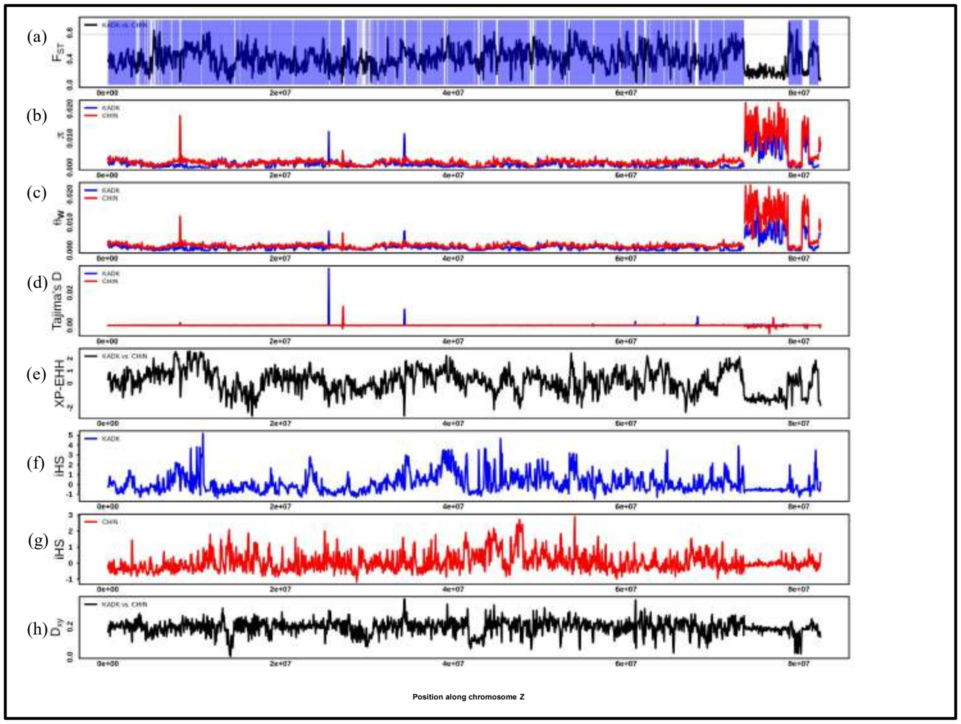


**Fig. S82. (a)** Pairwise $F_{ST}$ comparison between KADK and CHIN population along chromosome Z using 50Kb windows. A horizontal black dotted line represents the 99 percentile $F_{ST}$ threshold. >80 percent callable region shown in transparent blue color while white color region represents <80 percent callable region. **b,c,d.** represents the $\pi$, Watterson theta, and Tajima`s D, respectively, where the solid blue line represents the KADK, and the solid red line represents the CHIN population. **(e)** Pairwise XP-EHH comparison between KADK and CHIN using 50Kb window. **f, g.** iHS results visualized in 50Kb window along the chromosome for KADK shown in solid blue color and CHIN shown in solid red color, respectively. **(h)** Dxy between KADK and CHIN.

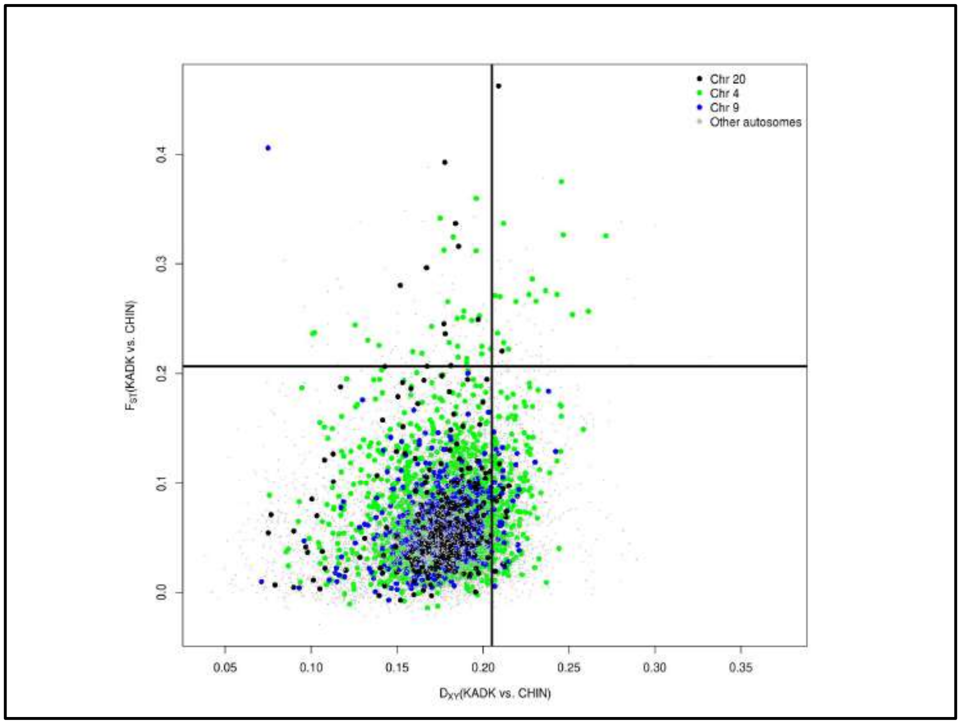


**Fig. S83**. $F_{ST}$ at Y-axis and Dxy at X-axis between KADK and CHIN population has been shown. Black color dots represent Chr 20, green color dots represents Chr 4, blue color dots represents Chr 9, and grey color dots represent other autosomes. The horizontal black color solid line represents the top1% of $F_{ST}$, while the vertical black color solid line represents the top10% of Dxy.

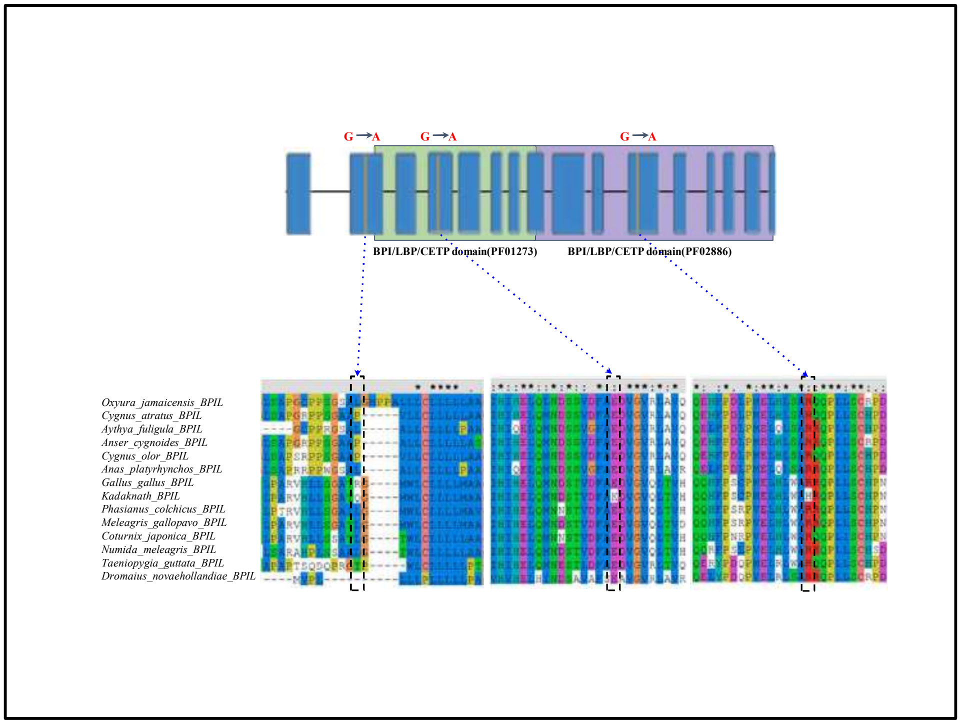


**Fig. S84.** Three nonsynonymous changes in *BPIL* gene. Rectangle boxes in blue color represent exons connected by a thin black line which represents introns. BPI superfamily domains are shown in light green and light purple colors. The zoomed view of amino acid change is shown by the blue color dotted arrow. The black color dotted rectangle represents the position of nonsynonymous changes**.**

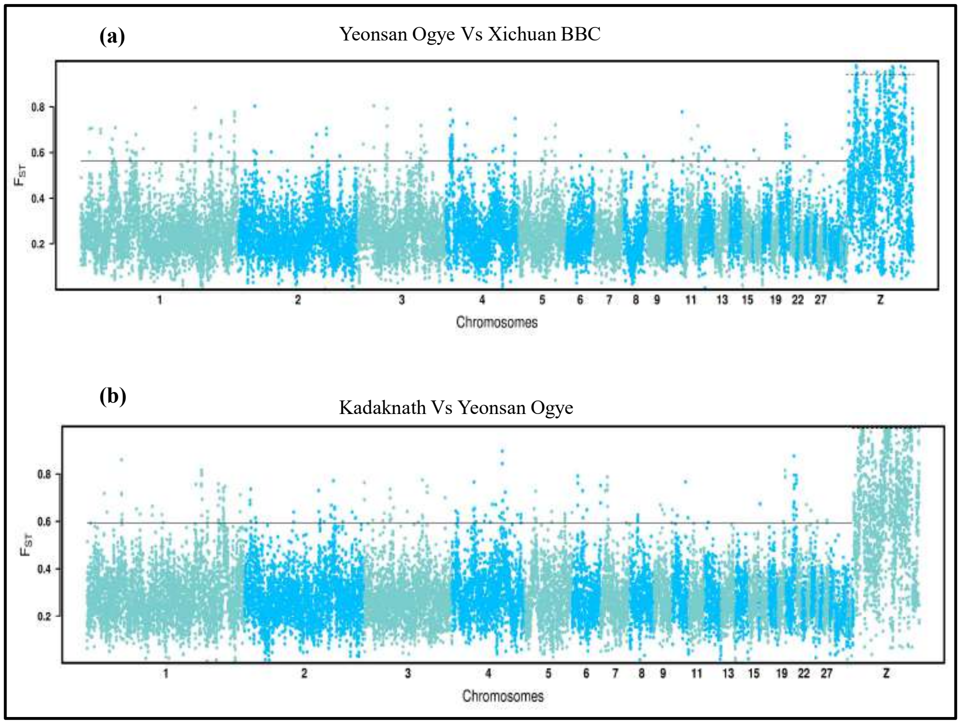


**Fig. S85:** Genome-wide landscape of pairwise genetic differentiation ($F_{ST}$) **(a)** between Yeonsan Ogye and Xichuan black-bone chicken **(b)**. between Kadaknath and Yeonsan Ogye in 50Kb non-overlapping windows. The dark slate gray and deep sky blue colors represent the alternative chromosomes. The dotted horizontal black line marks the 99th percentile outlier of estimated $F_{ST}$ for autosome and the Z chromosome, respectively.

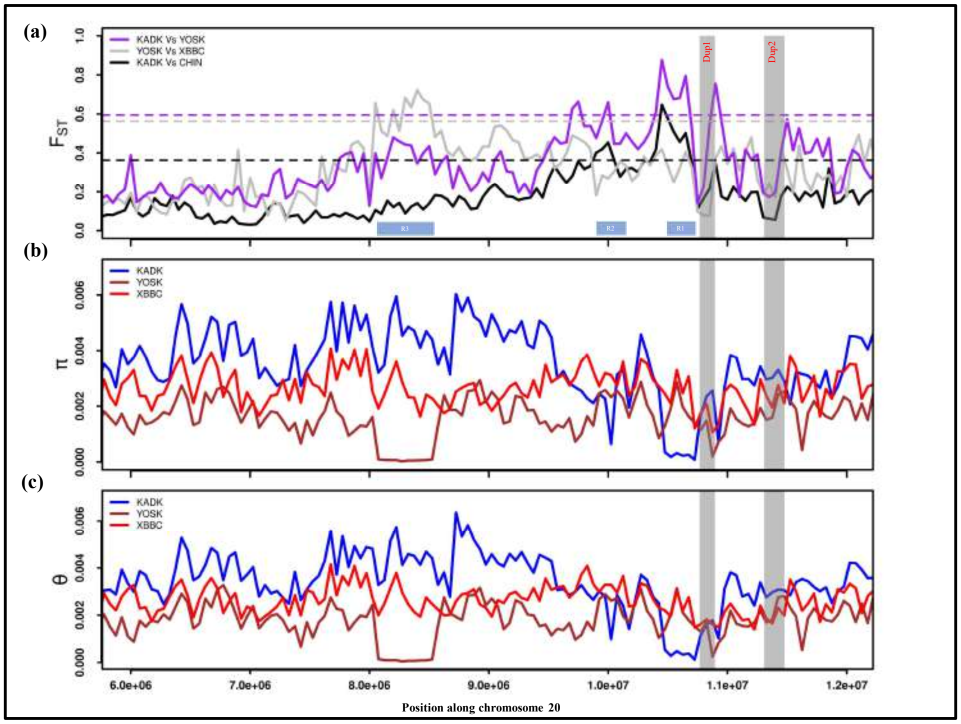


**Fig. S86: (a)**.Pairwise $F_{ST}$ comparisons between three population pairs along chromosome 20, where KADK vs. CHIN is in the solid black line, YOSK vs. XBBC in the solid grey line, and KADK vs. YOSK in the purple color solid line. The dotted horizontal lines of the same color represent the 99 percentile outlier for each population comparison. The Blue boxes represent the regions(R1 and R2) with high $F_{ST}$ identities, while the vertical transparent grey color represents the Dup1 and Dup2 regions in all panes. **(b)** within population pairwise nucleotide diversity (π) and **(c)** genetic diversity. The blue line represents the KADK population, the brown represents the YOSK population, and the red represents the XBBC population.

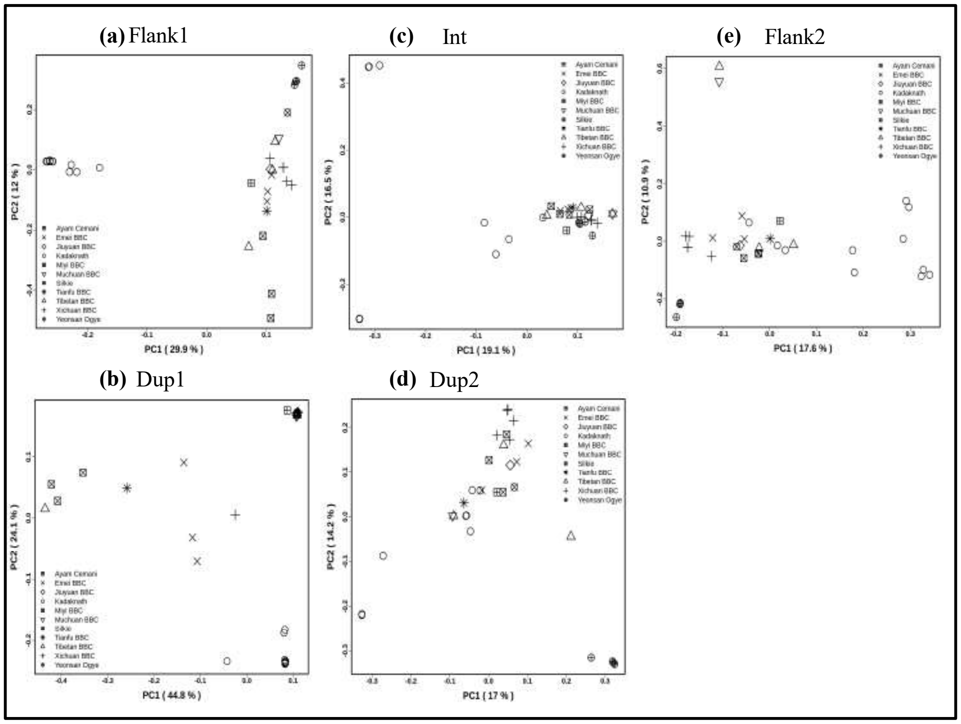


**Fig. S87.** Principal component analysis (PCA) of *Fm* region for 34 black-bone chickens with the first two principal groups (PC1 and PC2). **(a)** Flank1 **(b)** Dup1 **(c)** Int **(d)** Dup2 **(e)** Flank2. Each breed is shown in black color with different shapes for each breed.

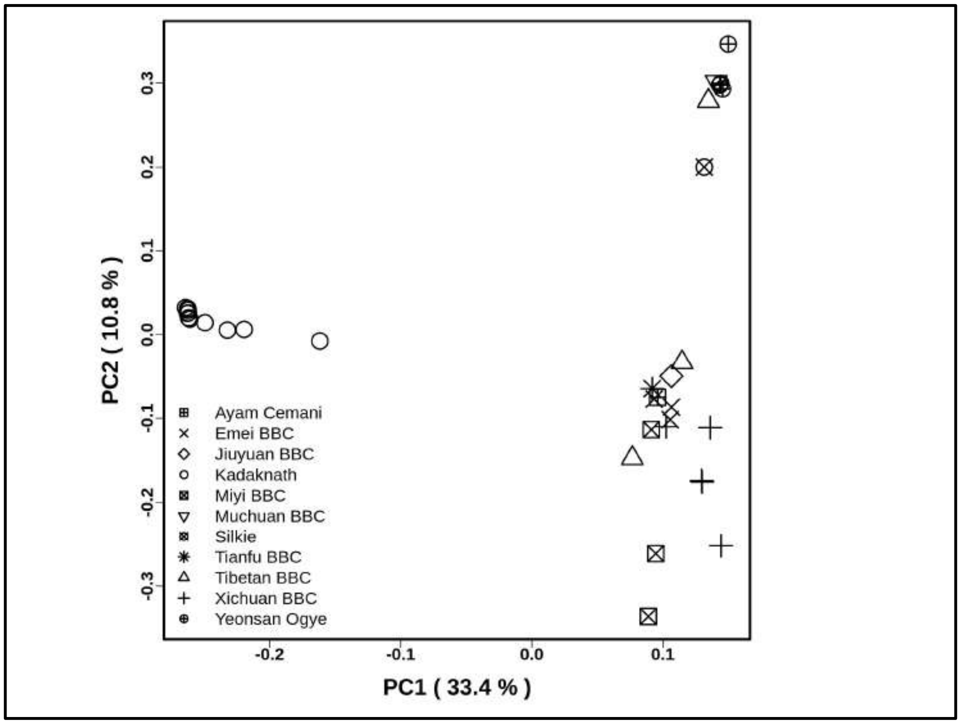


**Fig. S88.** Principal component analysis (PCA) of R1 region for 34 black-bone chickens with the first two principal groups (PC1 and PC2). Each breed is shown in black color with different shapes for each breed.

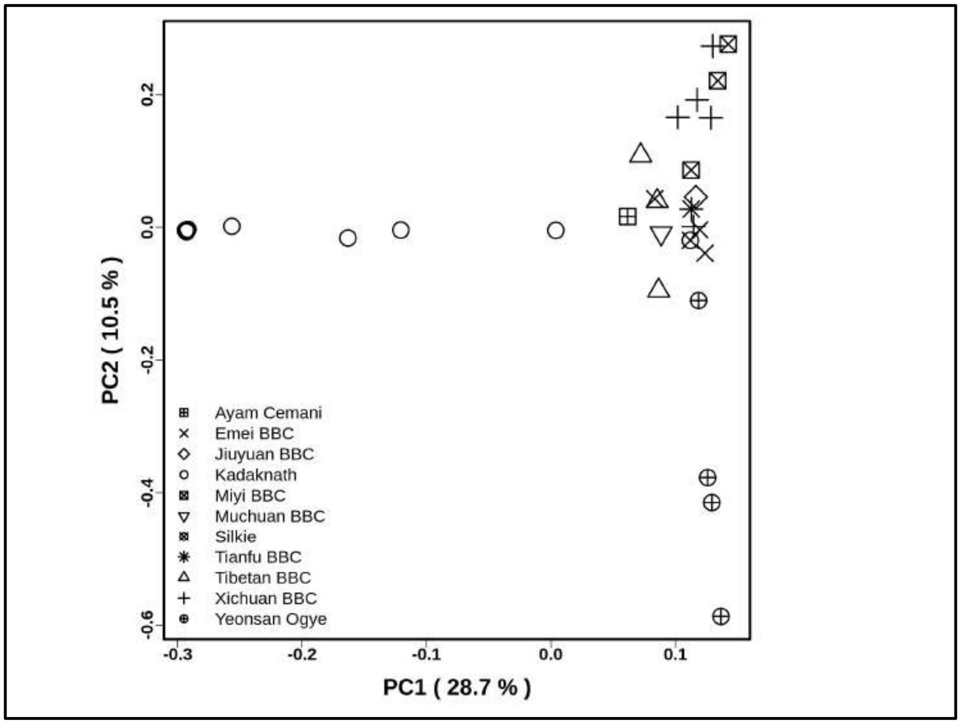


**Fig. S89.** Principal component analysis (PCA) of R2 region for 34 black-bone chickens with the first two principal groups (PC1 and PC2). Each breed is shown in black color with different shapes for each breed.

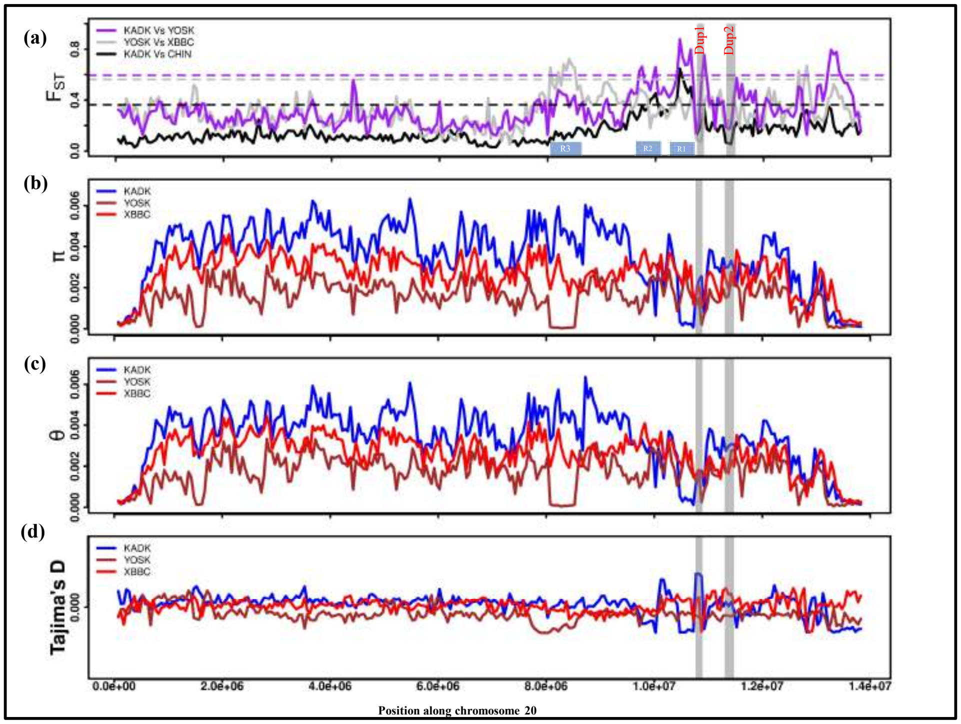


**Fig. S90: (a)**.Pairwise $F_{ST}$ comparisons between three population pairs along complete chromosome 20, where KADK vs. CHIN is in the solid black line, YOSK vs. XBBC in the solid grey line, and KADK vs. YOSK in the purple color solid line. The dotted horizontal lines of the same color represent the 99 percentile outlier for each population comparison. The Blue boxes represent the regions (R1, R2, R3) with high $F_{ST}$ identities, while the vertical transparent grey color represents the Dup1 and Dup2 regions in all panes. **(b)** within population pairwise nucleotide diversity (π),**(c)** genetic diversity and **(d)** Tajima's D . The blue line represents the KADK population, the brown represents the YOSK population, and the red represents the XBBC population.
